# Benchmarks in antimicrobial peptide prediction are biased due to the selection of negative data

**DOI:** 10.1101/2022.05.30.493946

**Authors:** Katarzyna Sidorczuk, Przemysław Gagat, Filip Pietluch, Jakub Kała, Dominik Rafacz, Laura Bąkała, Jadwiga Słowik, Rafał Kolenda, Stefan Rödiger, Legana C H W Fingerhut, Ira R Cooke, Paweł Mackiewicz, Michał Burdukiewicz

**Author notes:** These authors contributed equally to this work (Katarzyna Sidorczuk, Przemysław Gagat).

## Abstract

Antimicrobial peptides (AMPs) are a heterogeneous group of short polypeptides that target microorganisms but also viruses and cancer cells. Due to their lower selection for resistance compared to traditional antibiotics, AMPs have been attracting the ever-growing attention from researchers, including bioinformaticians. Machine learning represents the most cost-effective method for novel AMP discovery and consequently many computational tools for AMP prediction have been recently developed. In this article, we investigate the impact of negative data sampling on model performance and benchmarking. We generated 660 predictive models using 12 machine learning architectures, a single positive data set and 11 negative data sampling methods; the architectures and methods were defined on the basis of published AMP prediction software. Our results clearly indicate that similar training and benchmark data set, i.e. produced by the same or a similar negative data sampling method, positively affect model performance. Consequently, all the benchmark analyses that have been performed for AMP prediction models are significantly biased and, moreover, we do not know which model is the most accurate. To provide researchers with reliable information about the performance of AMP predictors, we also created a web server AMPBenchmark for fair model benchmarking. AMPBenchmark is available at http://BioGenies.info/AMPBenchmark.

## Introduction

Antimicrobial peptides (AMPs) are short polypeptides, generally composed of up to 50 amino acids that are widespread in all forms of life, from microorganisms, i.e. bacteria, archaeans and one-celled eukaryotes, to multicellulars [1, 2]. In microorganisms, they participate in self-protection and microbial competition [3, 4]; in multicellulars, they are part of the first line of defence against microorganisms but also target viruses and cancer cells [5, 6]. Despite their diversity in the primary amino acid sequence, AMPs are rich in cationic and hydrophobic residues. The positive charge and hydrophobicity allow them to fold into amphipathic secondary structures that preferentially disrupt negatively-charged microbial/cancer cell membranes but not the healthy eukaryotic ones; the latter contain stabilizing cholesterol and their outer leaflet is composed of neutral phospholipids. AMPs can trigger transient membrane disruption by forming pores and micellization but, depending on the concentration, they may lead to cell death by osmotic shock [7, 8, 9, 10]. The alternative mechanisms of action, especially for the larger AMPs (about 100 amino acids long or longer), include binding to specific cytosolic macromolecules and thereby inhibiting synthesis of proteins, nucleic acids and components of the cell wall [11, 12].

AMPs have also been demonstrated to have lower selection for resistance compared to traditional antibiotics. A traditional antibiotic specifically targets a single enzyme but AMPs, most of all, interact non-specifically with many components of the cell membrane. This makes it more difficult for bacteria to develop resistance against them [13, 14, 15].

According to the World Health Organization, the antibiotic resistance is currently behind the death of at least 700,000 people each year; however, the forecast of the death toll of 10 million annually by 2050 makes the race for alternative therapeutics of the utmost importance [16]. In light of their medical potential, AMPs are viewed as hopeful candidates for further experimental research. Consequently, we have recently observed a boom in computational tools for AMP prediction with the machine learning algorithms leading the way [17].

Traditionally, biological problems have first been approached by conventional, i.e. non-deep machine learning-based methods, such as random forests (RF) or support vector machines (SVM), which were then followed by more complex deep learning algorithms [17]. In order to produce reliable predictions, the algorithms first require labelled training data to build a predictive model. The training data include a positive and a negative data set, in our case AMPs and non-AMPs, respectively. In order to make the sequences readable for machine learning, they have to be transformed into informative features (feature vectors) and this process is known as feature extraction. Depending on the method of feature extraction, the obtained feature space may require additional reduction, and consequently an appropriate feature selection method is applied, e.g. for AmpGram the initial feature set amounted to 33,620 n-grams (amino acid motifs of n elements) but was decreased with Quick Permutation Test to 13,087 most informative descriptors [18].

There are many databases with thousands of experimentally validated AMP sequences, such as DBAASP [19], APD [20], CAMP [21], DRAMP [22] or dbAMP [23]; therefore, it is possible to create a representative positive data set. However, the authors of AMP classifiers, except for ampir [24], do not take into account that there might be two types of sequences deposited in these databases: mature AMPs and precursor AMPs with cleavable N-terminal signal peptides; AMPs are mostly secretory proteins. Since the databases seem to contain generally mature AMPs, and moreover the developers often restrict the sequence length in their data sets, the algorithms are mainly trained on mature AMPs. Consequently, they are good at detecting mature AMPs but might have problems classifying longer sequences, including the precursor proteins [25].

The issue of identification of precursor and longer AMPs can be satisfactorily addressed because the data about these sequences are available in public databases, e.g. in UniProt [26]. The real problem with AMP prediction lies with the negative data set as there are hardly any sequences annotated as non-AMPs. Interestingly, the lack of reliable negative samples also concerns other areas related to bioinformatics, e.g. prediction of disease genes [27, 28], microRNAs [29], bacterial virulence factors [30]; identification of protein-protein [31], protein-RNA/DNA [32, 33] and protein-drug interaction sites [32, 34]; as well as inferring protein sequence-function relationships [35].

In all these cases, the developers have to resort to: (i) one-class classification, (ii) positive-unlabelled learning or (iii) to somehow build a negative data set. In the first case, the model is trained on the positive sample whereas in the second on the positive and unlabelled data; the unlabelled set includes both positive and negative examples. These two approaches, aim at solving the problem of the negative sample by either not using it at all or applying a wide variety of strategies to obtain negative cases from the unlabelled set based on the positive sample, e.g. using distance measures (for details, see [36, 37]). Interestingly, neither the one-class classification nor the positive-unlabelled learning have attracted the attention of developers working on AMP prediction. The majority of them created their negative sets by performing non-probability sampling on sequences deposited in UniProt [26] or other databases (e.g. PDB [38]) though they do not define it as such. In this approach, the negative examples are selected on the basis of clearly defined criteria dictated by the researcher, and these criteria represent a sampling method (for details, see section Materials and methods and Table 1). In contrast to positive-unlabelled learning, dividing UniProt sequences between AMPs and non-AMPs does not require any complex methodology and is independent of the positive sample but for the length and number of sequences for some sampling methods. It is simply made by sequence filtering and then randomly selecting peptides for the final negative data set. For clarity purposes in this article, the name of the sampling method is always preceded by an abbreviation: SM (sampling method), TSM (sampling method used to generate the training set) or BSM (sampling method used to generate the benchmark set) and colon, e.g. SM:AmpGram, TSM:AmpGram and BSM:AmpGram, respectively.

**Table 1.** A comprehensive summary of the negative sampling methods implemented for AMP prediction.

| Sampling method | Excluded keywords | Sequence lengths | CD-HIT | Additional filtering | Balanced classes | Reference |
| --- | --- | --- | --- | --- | --- | --- |
| Wang et al. | secreted, antimicrobial | similar to the length distribution of positive dataset | 0.7 |  | no | [41] |
| CS-AMPpred | antimicrobial | 16 - 90 | 0.4 |  | yes | [42] |
| iAMP-2L | antimicrobial, antibiotic, fungicide, defensin | 5 - 100 | 0.4 |  | no | [43] |
| Gabere&Noble (DAMPD dataset) | antimicrobial | equal to the length distribution of positive dataset | no | localisation: golgi, cytoplasm, endoplasmic reticulum or mitochondria | no | [25] |
| AMAP | antimicrobial | similar to the length distribution of positive dataset | 0.4 |  | no | [44] |
| AMPScanner V2 | antimicrobial, antibiotic, antiviral, fungicide, secreted | equal to the length distribution of positive dataset | no | localisation: cytoplasm; removed sequences similar to any AMPs with BLAT [45] | yes | [46] |
| dbAMP | transmembrane, toxin, secreted, defensin, antimicrobial, antibiotic, antiviral, fungicide | 10 - 100 | 0.4 |  | no | [23] |
| Witten&Witten | antimicrobial, antibiotic, antiviral, fungicide, secreted | equal to the length distribution of positive dataset | 0.4 | localisation: cytoplasm; only cysteine-free substrings | yes | [47] |
| AmpGram | antimicrobial, antibacterial, antiviral, fungicide, secreted, transit peptide | equal to the length distribution of positive dataset | no |  | yes | [18] |
| ampir-mature | antimicrobial | 10 - 40 | 0.9 |  | no | [24] |
| AMPlify | antimicrobial, antibiotic, defense, defensin, bacteriocin, fungicide | equal to the length distribution of positive dataset | no | removed potential AMPs* | yes | [48] |

\*UniProt sequences with keywords: antimicrobial, antibiotic, defense, defensin, bacteriocin, fungicide

The aim of this study was to elaborate on the impact of negative data sampling on model performance and benchmarking. We decided to explore this issue because each developer of an AMP predictor tested only one sampling method to build the optimal non-AMP class despite knowing that machine learning models heavily depend on the data sets they are trained on. They all overlooked the fact that various sampling methods could generate statistically different samples, thereby affecting the predictive power of their models. Moreover, and more importantly, we investigated how machine learning architectures perform when they are trained on a given data set but tested on a different one, a commonplace in model benchmarking (Figure 1A). This particular issue is of vital importance not only for the comparison of AMP predictors but for the evaluation of all machine learning models in general. We define the machine learning architecture as an approach to solve the problem of AMP prediction with all its parameters involved in the machine learning cycle. The architectures for our study were developed based on published models that we were able to reuse or reimplement; some might slightly deviate from the original methods (for details, see section Materials and methods and Table 2, S2). For clarity purpose, the name of a given architecture always begins with a letter ‘A’ and colon, e.g. A:AmpGram. By the term machine learning model, we understand one specific instance of a given architecture, i.e. an architecture trained on the same positive and one of negative samples. Consequently, what we did was to generate 660 machine learning models using (i) 12 defined architectures, (ii) the same positive training data set and (iii) 11 different negative sampling methods each run five times (Figure 1B). To our knowledge, this was the first kind of such a research project undertaken, and moreover on such a scale.

**Fig. 1.**
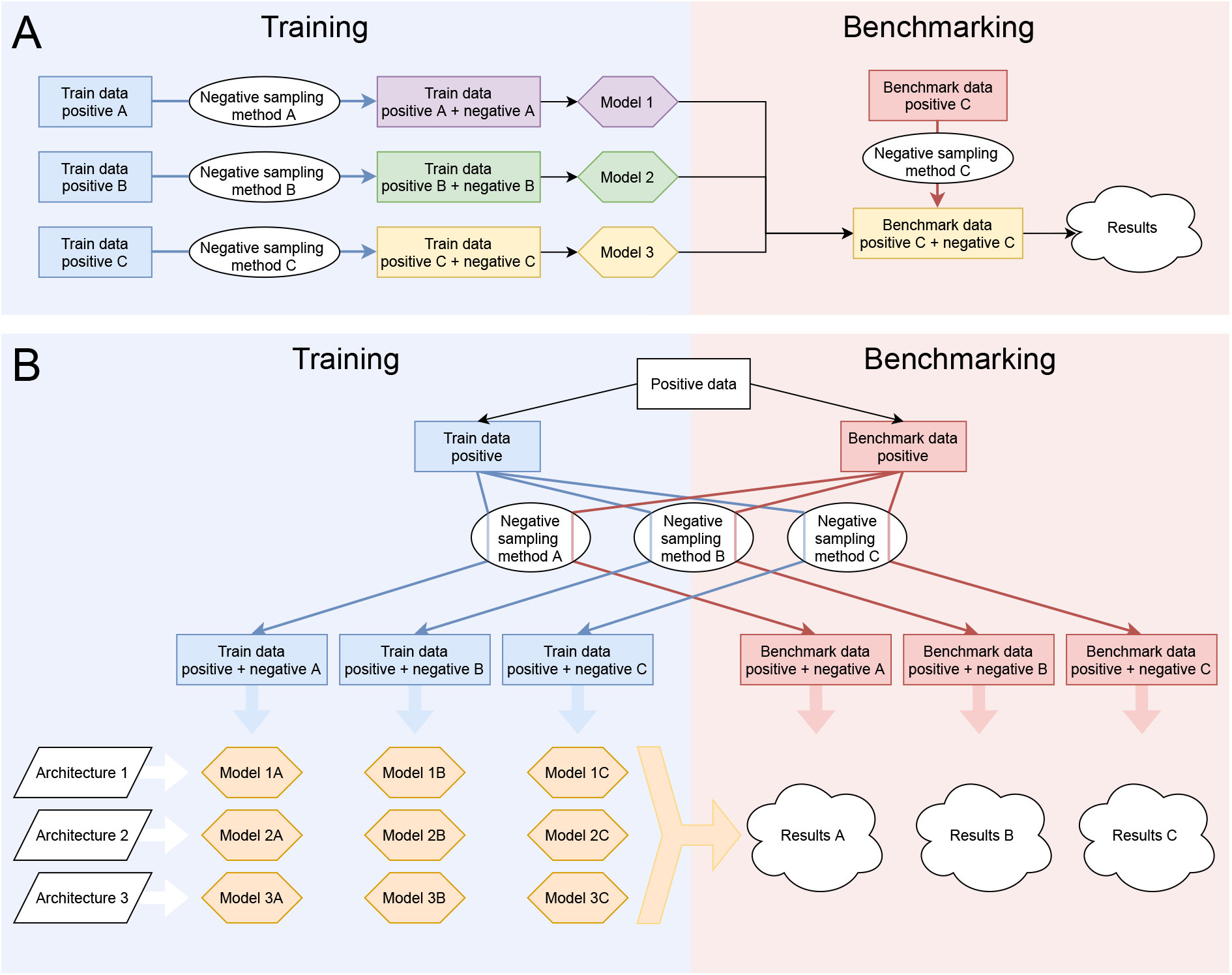
Schematic representation of the traditional model benchmarking (A) and the methodology employed in comparing the impact of different negative data sampling methods on model performance (B). Model 1, 2 and 3 (colourful hexagons) were trained on data set A, B and C (colourful rectangles), respectively. Each data set was generated by an appropriate negative sampling method (white ovals) and a positive sample (blue rectangles). In the evaluation process, the models were compared only on the benchmark set C, built with the same method as the training set C, thereby introducing some bias in favour of model 3 in the benchmark analysis (A). Architectures were developed based on published models, and they represent the algorithm with all its parameters involved in the machine learning cycle (white parallelograms). Each architecture was trained on the same positive data set (the white rectangle) and a negative sample generated by one of 11 negative sampling methods (white ovals) five times to verify the repeatability. The training and benchmark sample are indicated as blue and red rectangles, respectively. The models (orange hexagons) represent instances of architectures trained on given data sets and were validated on each benchmark sample. The results of model performance were indicated as white clouds (B).

**Table 2.**
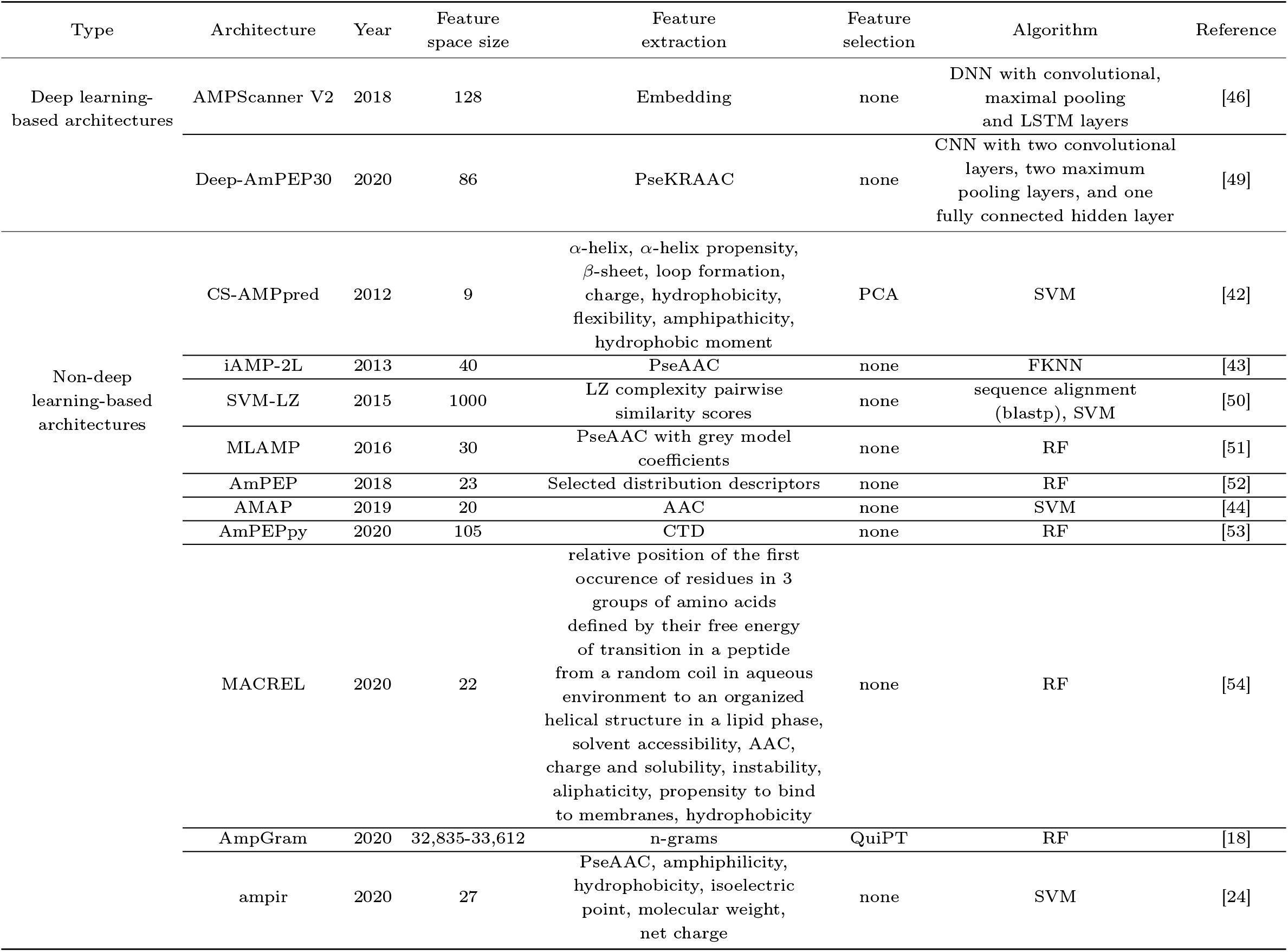
A comprehensive summary of the implemented architectures for AMP prediction.

## Materials and methods

### Data sets

To create the positive data set, we used DBAASP v3.0 [19], a manually-curated database for experimentally verified peptides with antimicrobial properties. We selected sequences with activity against Gram positive or Gram negative bacteria. Next, we removed those with non-standard amino acids or shorter than five. We used CD-HIT version 4.8.1 [39, 40] to reduce the redundancy and eliminated sequences with the identity threshold higher than 90%. This threshold was most frequently used for the reduction of positive data in the algorithms selected for the reimplementation of negative sampling methods (Table S1). In total, we obtained 5,190 AMP sequences. To prevent the information leakage, the positive data set was split, before sampling the negative set, into a training sample (80%, 4151 sequences) and a benchmark sample (20%, 1039 sequences).

The negative data set used for sampling was created using sequences available in the UniProt database. The reviewed protein sequences (563,972) and their annotations were downloaded from the UniProtKB release 2020_06 [26].

We considered 26 methods of negative data sampling from literature (Table S2) and selected 11 well-described ones for reuse/reimplementation (Table 1). Each method was run on the negative data set of UniProt sequences five times to create five replicates of the training and benchmark samples (Table S4, S5) to verify their repeatability. Some selected methods required modifications, e.g. removal of sequences with non-standard amino acids to make them readable for all architectures. We also took measures to prevent information leakage between the training and benchmark sets, especially for the sampling methods that did not depend on the positive data. The full description of changes is provided in the Supplementary Data.

The selected sampling methods use combinations of keywords to search the negative data set of UniProt sequences. Generally the keywords are repetitive among the methods used. They allowed to filter out AMPs by naming their typical functions, e.g. ‘antimicrobial’, ‘antibacterial’, ‘antiviral’, and/or restrict the cellular compartment to cytoplasm since AMPs are mostly secretory proteins (Table 1). The latter is, however, unfortunate because the predictive models focus then on differences between cytoplasmic and secretory peptides instead of detecting AMPs, i.e. they neglect the issue of (i) distinguishing AMPs from secretory non-AMPs and (ii) cytosolic AMPs. The number of the filtered-out sequences was small if only the function keywords were considered, e.g. about 1% of the negative data set of UniProt sequences for SM:CS-AMPPred, SM:AMAP and SM:iAMP-2L, but increased when the cytosolic or additional location, especially experimentally verified, was included to 65%, 70% and 98% for SM:Gabere&Noble, SM:AMPScannerV2 and SM:Witten&Witten, respectively.

For five methods, SM:AmpGram, SM:AMPlify, SM: AMPScannerV2, SM:CS-AMPPred and SM:Witten&Witten, the number of sequences in the negative sample depended on the positive one, i.e. the data sets were balanced. Two methods: SM:AMAP and SM:dbAMP generated only slightly imbalanced samples, the former due to the reduction of sequence redundancy with CD-HIT [40] at the end of the sampling process but the latter by accident. The remaining methods produced imbalanced (SM:iAMP-2L, SM:Wang et. al) or highly imbalanced (SM:Gabere&Noble) sets with the predominance of non-AMPs. The exception was SM:ampir-mature with minority of non-AMPs (Table 1, S4, S5).

Five negative sampling methods: SM:AmpGram, SM: AMPlify, SM:AMPScannerV2, SM:Gabere&Noble and SM: Witten&Witten produced non-AMPs that exactly matched in length peptides and proteins contained in the positive set, and they were mostly up to 50 amino acids long though the sequence maximum length was 190 amino acids. The negative samples of SM:AMAP and SM:Wang et. al were only similar in terms of length distribution to the AMP set because of CD-HIT [39, 40] reduction at the end of sequence filtering. The SM:ampir-mature generated only short sequences, and each sequence length within the range of 10 to 40 amino acids was approximately equally represented (Table 1, Figure S1). The other sampling methods focused on longer peptides and proteins though at the same time they rejected sequences longer than about 100 amino acids. These methods included SM:dbAMP, SM:CS-AMPPred and SM:iAMP-2L and their sequence length distribution resembled that of an upside-down isosceles triangle (Table 1, Figure S1).

All the negative data sampling methods generating sets with equal or similar length distribution to the positive sample selected their non-AMPs from peptide/protein fragment or fragments while the other methods from uncut sequences of the negative UniProt data set.

In order to avoid overrepresentation of highly similar sequences in the non-AMP samples, we used the clustering algorithm CD-HIT version 4.8.1 [40] for seven sampling methods according to their description. We removed sequences above a certain identity threshold, and mostly it was 40% (Table 1).

Interestingly, SM:AMPlify, SM:AMPScanner V2 and SM:ampir-mature additionally verified whether the negative data set did not, by chance, contain sequences from the positive data set (Table 1). This might arise as a result of: (i) non-AMPs generation from a protein fragment or fragments, and (ii) an improper/lack of annotation in UniProt [26] for sequences that are indeed antimicrobial.

The sampling methods generated sequences that greatly differed from those in the positive data set both in the amino acid composition (Figure S2-S6) and physicochemical properties (Figure S7). There were also some pronounced differences among the negative sets, but the five iterations of each method always produced similar samples (Figure S2-S8).

### Model architectures

We considered 26 model architectures for the prediction of AMPs from literature (Table S2), and selected 12 for reimplementation, two deep learning and ten non-deep learningbased methods, the latter represent mainly RF and SVM algorithms (Table 2). In our research, we chose algorithms described in detail that could be run locally and do not require any usage of web servers or other software for feature selection. Moreover, we focused on the classification task, i.e. the model ability to divide sequences into AMPs or non-AMPs. Consequently, we did not consider software that is trained using MIC (minimum inhibitory concentration) values, e.g. Witten&Witten or multiclass models. However, if the multiclass algorithm was composed of two models: one predicting if a given sequence is or is not an AMP, and the second classifying an AMPs into functional groups, we did reimplement the first model, e.g. for A:iAMP-2L and A:MLAMP.

For each model, sequences in the data sets were transformed (encoded) into features (descriptors) using an appropriate feature extraction method (Table 2). In the case of nondeep learning architectures, the features can be defined by the researcher, based on the knowledge of AMP properties, whereas the deep learning architectures can automatically learn high-level features from the training data sets though A:Deep-AmPEP30 also employed developer-defined features.

The simplest feature extraction method was used by A:AMAP, and it was based on the amino acid composition. Consequently, its feature space contained 20 descriptors, each reflecting the occurrence frequency of one of the 20 amino acids in a peptide sequence (Table 2). Other architectures, such as A:iAMP-2L, A:MLAMP, A:ampir and A:Deep-AmPEP30 used features based on pseudo-amino acid composition. Beyond the simple amino acid counts, they included various physicochemical and structural properties of amino acids to incorporate the information about the sequence order; their feature space increased accordingly (Table 2). Four architectures: A:CS-AMPPred, A:MACREL, A:AmPEP and A:AmPEPpy used features based on structural and physicochemical properties of peptide sequences, e.g. their *α*-helix propensity, charge and hydrophobicity (Table 2). A:SVM-LZ used pairwise similarity scores, and A:AmpGram n-grams. In the case of A:AMPScanner V2 the feature information was extracted in the embedding layer, and then the obtained embeddings fed the subsequent layers of the models (Table 2).

Five non-deep machine learning architectures used feature selection for feature space optimisation, but we implemented it for only two architectures: A:CS-AMPpred and A:AmpGram. The former used principal component analysis and the latter Quick Permutation Test (Table 2). For A:AmPEP and A:ampir, we used the reduced feature space indicated by their developers without computing Pearson correlation coefficients and rigorous recursive feature elimination, respectively. For A:AmPEPpy, we abandoned the reduction of feature space by stepwise feature selection because according to the authors it did not improve the predictive power of the model but only its size.

Five of the selected architectures required only slight modifications in their already available codes, and seven were implemented based on the information provided by the authors either in their articles or personal communication. A comprehensive description of the implementations and the applied modifications are provided in the Supplementary Data, including Table S3.

## Results

### The impact of data sampling on benchmarks

In order to evaluate the impact of data sampling on benchmark analysis, the ROC (receiver operating characteristic) curves were plotted (Figure S9-S69) and values of the area under the ROC curve (AUC) were calculated for each of 660 models on each benchmark data set and then averaged for the appropriate architecture. The results of the analysis are presented in Figure 2A, S70 and S71. They clearly indicate that all but two architectures, A:AmPEP and A:iAMP-2L, performed much better when the training and benchmark samples were generated by the same sampling method. A:SVM-LZ, A:AmpGram and A:CS-AMPPred showed only small improvement of 2.3%, 3.6% and 4.4% respectively; however, A:AmpGram by far outperformed the other architectures. The mean value of AUC for A:AMAP and A:MACREL increased 7.5%, for A:ampir and A:AmPEPpy about 9.5%, and for the rest architectures soared more than 10% (Table S6). A:AmPEP and A:iAMP-2L were the only architectures, which preferred dissimilar sampling methods for training and benchmarking, but both generally performed very poorly with mean AUC amounting to 0.65. The calculated differences were statistically significant for all comparisons but for iAMP-2L and SVM-LZ (Kruskal-Wallis test with Bonferroni correction, p-value < 0.05, Table S7).

**Fig. 2.**
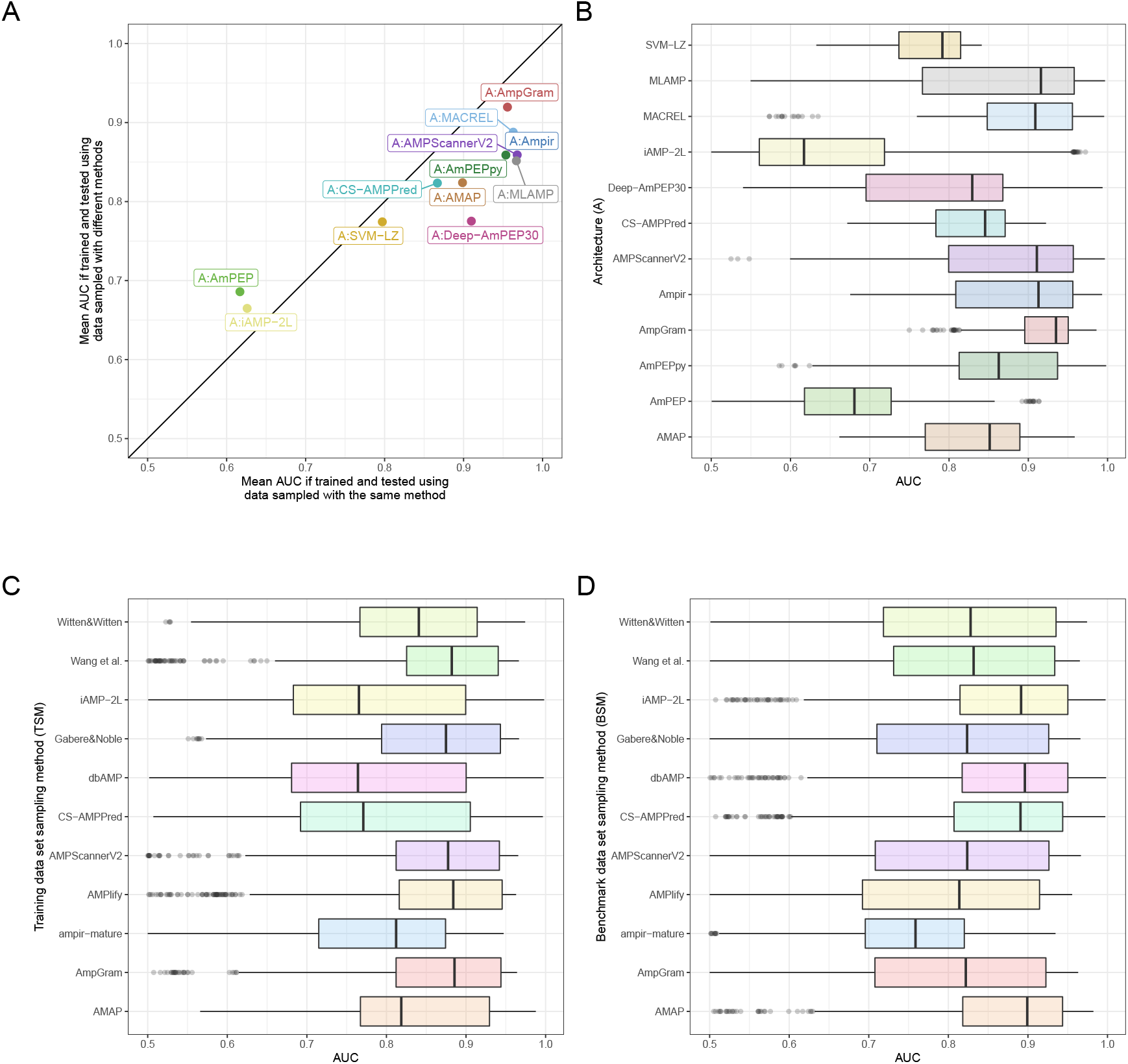
Model performance depending on the architecture and negative data sampling method used for training and benchmarking. The x-axis represents mean AUC for architectures trained and tested on sets generated by the same negative data sampling method. The y-axis represents mean AUC for architectures trained and tested on sets generated by different negative data sampling methods. The architectures on the right of the diagonal perform better when the training and benchmark sample are produced by the same method while the architectures on the left when the methods are different (A). Box plots with median and interquartile range differences in AUC for architectures (B), training data set sampling method (C) and benchmark data set sampling method (D).

The main conclusion from these analyses is that similar training and benchmark data set positively affect model performance. Accordingly, there was significant negative correlation between mean AUC value and the difference in amino acid composition between the training and benchmark sets, measured as the square root of the sum of the squared differences in the frequency of individual amino acids (Spearman correlation coefficient, *ρ* = −0.53, p-value < 2.2e-16). There was also smaller but still significant negative correlation for mean AUC and the absolute difference between median length of the sets (*ρ* = −0.44, p-value < 2.2e-16).

### The impact of architecture, training and benchmark data sampling method on model performance

To visualise which of the three components of the machine learning model: (i) architecture, (ii) training or (iii) benchmark data sampling method, bears the greatest importance for model performance, we compared box plots of AUC distribution for each of these features (Figure 2B, C, D). The plots clearly indicate the greatest variation of AUC for data grouped according to the architecture. We also calculated the ratio of between-group MAD (median absolute deviation) to within-group MAD to verify if the AUC dispersal between different architectures or training/benchmark data sampling methods is much greater than the AUC dispersal found inside a single architecture or method. The MAD ratios amounted to 1.29, 0.48 and 0.29 for architectures, training and benchmark data sampling methods, respectively, and express in numbers the relative AUC variation presented in the graphical form in the box plots (Figure 2B, C, D). Additionally, we conducted pairwise Wilcoxon test for paired samples between groups of the three components (Table S8-S10). The statistically significant differences (after Bonferroni correction) were indicated for 86%, 60% and 62% comparisons for groups of architectures, training and benchmark data sampling methods, respectively.

Unquestionably, the greatest differences in AUC are associated with the architecture indicating that this component is more important than the training and benchmark data sampling method for model performance. Among the five architectures with the median AUC value greater than 0.9, there were three using RF (A:AmpGram, A:MACREL and A:MLAMP), one SVM (A:ampir) and one deep learning algorithm (A:AMPScannerV2) (Figure 2B). The results emphasize the power of RF-based architectures in tackling the problem of AMP prediction. The only RF architecture that stood out with small median AUC value was A:AmPEP, which most probably results from the reduction of its feature space to 23; A:AmPEPy, a python implementation of A:AmPEP, with the full feature set of 105 performed quite well. The best architecture was A:AmpGram with the median AUC value of 0.93 and the narrowest box indicating low variance of AUC obtained for various combination of training and benchmark data (Figure 2B). About 73% models of this architecture obtained AUC greater than 0.9. The A:AmpGram performance indicates that short amino acid motifs have greater discriminatory power than amino acid composition or physicochemical and structural properties of amino acids and peptides. This seems true considering that the vast majority of AMPs form cationic, amphipathic *α*-helices, i.e. structures determined by an appropriate placement of amino acids with certain properties.

We also noticed a certain trend in the distribution of AUC for the training data sampling methods (Figure 2C). The AUC values calculated for TSM:AmpGram, TSM:AMPlify, TSM:AMPScannerV2, TSM:Gabere&Noble and TSM:Wang et. al were generally higher than for TSM:AMAP, TSM:ampir-mature and TSM:Witten&Witten, and the lowest AUC values were for TSM:CS-AMPPred, TSM:dbAMP and TSM:iAMP-2L. Interestingly, the first five sampling methods produced data sets similar in terms of length distribution (Figure S1), amino acid composition (Figure S2-S6) and physicochemical properties (Figure S7) that deviated from the other sets, especially those with the lowest median AUC. Given that there was significant negative correlation between mean AUC value and the difference in amino acid composition and median length between the training and benchmark sets (see above), it is not surprising that architectures trained and benchmarked on the five similar sampling methods outperformed the others. They simply were advantaged classifying benchmark sequences in accordance with our finding that similar training and benchmark sample positively affect model performance.

Contrary to the results presented in the prior paragraph, the sampling methods that performed worse as training sets (SM:dbAMP, SM:iAMP-2L, SM:CS-AMPPred and also SM:AMAP), turned out with the highest AUC as benchmark samples (Figure 2D). This can be explained by the fact that these methods generate sequences that not only differ from AMPs of the positive set in the amino acid content and physicochemical properties but are also generally much longer (Figure S1). The median values for sequences of SM:dbAMP, SM:iAMP-2L, SM:CS-AMPPred, SM:AMAP and the positive sample are: 79, 79, 72, 36 and 18, respectively. Accordingly, we found significant positive correlation between mean AUC and differences in the median length of the benchmark negative data sets and the benchmark positive sample (Spearman correlation coefficient, *ρ* = 0.74, p-value = 8.63e-11).

### Repeatability of prediction for the replicates of data sets

To verify the repeatability of prediction, each architecture was trained and benchmarked on five replicates of the training and benchmark sample. Despite the fact that the replicates were very similar in terms of length (Figure S1), amino acid composition (Figure S2-S6) and physicochemical properties (Figure S7), they did affect the performance of our investigated architectures (Figure 3), especially A:iAMP-2L, A:AMPScannerV2 and A:Deep-AmPEP30, for which mean standard deviation (SD) of AUC value amounted to 0.035, 0.019 and 0.014, respectively (Table S11). A:iAMP-2L generally resulted in poorly performing models and the majority of them were characterised by low repeatability. Similarly, A:Deep-AmPEP30 was not a very robust architecture and some models trained on the replicated data also produced significantly different AUC values, e.g. those trained on SM:ampir-mature, SM:AMPScannerV2 and SM:AMPlify, and benchmarked on SM:CS-AMPPred, SM:dbAMP and SM:iAMP-2L. In contrast, A:AMPScannerV2 was rather at the forefront of the investigated architectures, especially if the training and benchmark set were generated by the same sampling method (Figure 2A, B). In the case of A:AMPScannerV2, there is a clear pattern of overlapping low AUC and high SD distribution for models trained on replicated data generated by SM:ampir-mature, SM:CS-AMPPred, SM:dbAMP, SM:iAMP-2L and also SM:Witten&Witten (Figure 3). It looks like these training sets were not enough for deep learning models to learn features necessary to classify AMPs and non-AMPs correctly. This result concurs with other works reporting that shallow models have similar performance to deep ones for AMP prediction [55].

**Fig. 3.**
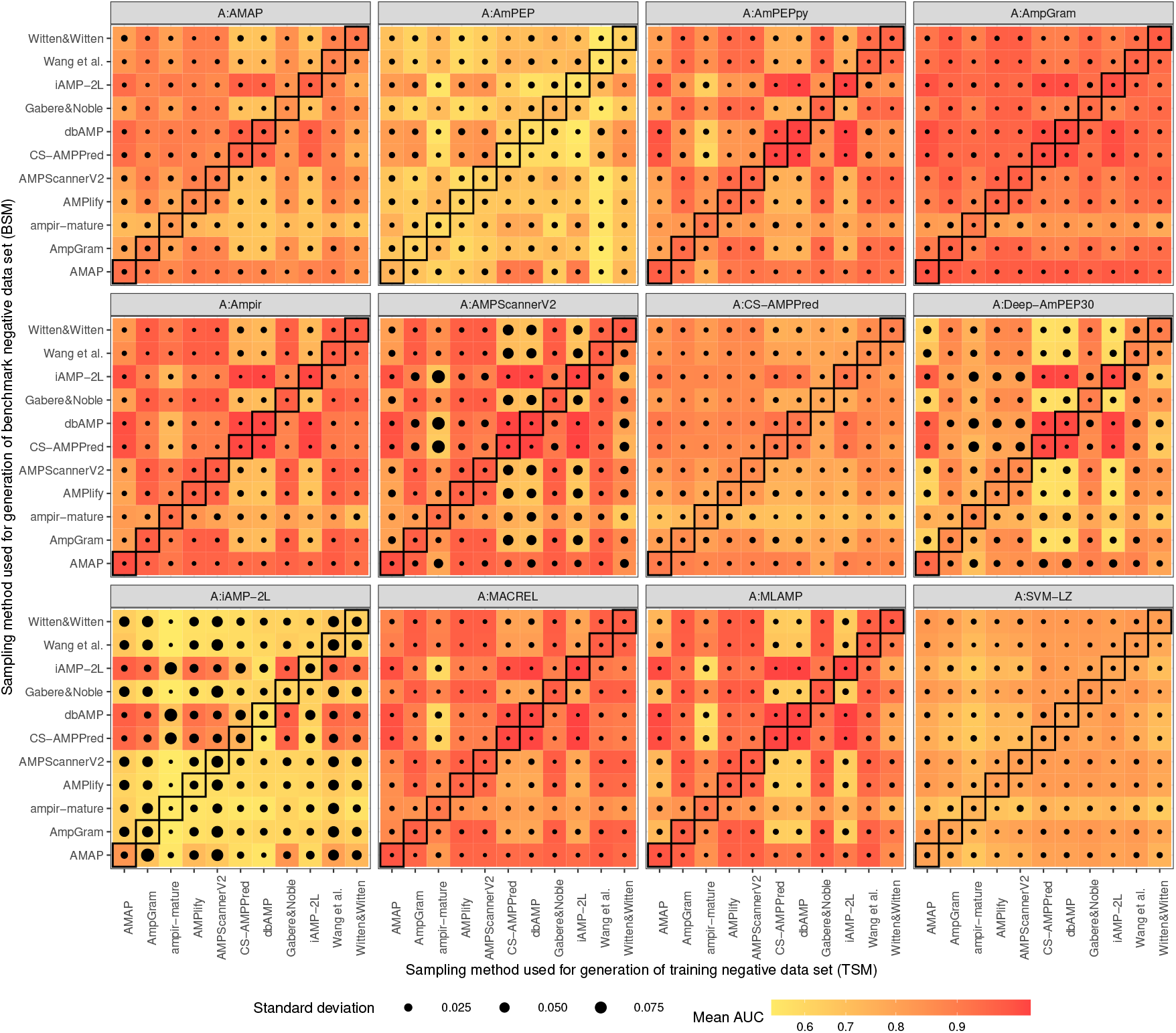
Architecture performance depending on the negative data sampling method used for training and benchmarking. Each of 12 heat maps represents an architecture, the x- and y-axis describe the training and benchmark method of negative data sampling, respectively. Each architecture was trained and benchmarked on five replicates of the training and benchmark sample. The mean value of AUC for the five replicates is indicated as shades of red, orange and yellow, and the standard deviation as black dots of varying sizes. The diagonals mark results for architectures trained and benchmarked on the data generated by the same sampling method.

It is worth emphasising that the most stable architectures included also the best ones: A:AmpGram, A: MACREL and A:ampir, as well as A:AmPEPpy (Table S11, Figure 2A, 2B, 3, S70). Their mean SD of AUC value amounted to about 0.004. A:ampir implemented SVM and the rest RF algorithm indicating their superiority over deep learning architectures (A:AMPScannerV2 and A:Deep-AmPEP30) in tackling our data.

We also noticed a certain trend in the distribution of AUC reflecting a previously formulated conclusion that similar training and benchmark data set positively affect model performance. It was less noticeable for poorly performing architectures: A:iAMP-2L, A:AmPEP and A:SVM-LZ, and A:AmpGram representing the top architecture in our studies (Figure 3).

## Discussion and conclusions

Machine learning represents the most cost-effective method for novel AMP discovery. As a result, many computational tools for AMP prediction have been developed in recent years [17] and each subsequent state-of-the-art model claims to outperform its predecessors. As a rule, the state-of-the-art model is evaluated with other software on a benchmark sample generated by the same method that was also used to produce its training set (Figure 1A). According to the presented research, this is a source of statistically significant bias in favour of the state-of-the-art model because the more similar the training and benchmark data set are the better the model performance (Figure 2A, 2B, 3, S70). Consequently, we came to logical conclusions that (i) all the benchmark analyses that have been published for AMP prediction tools are unfair and (ii) we do not know which model is the most accurate.

To provide researchers with reliable information about the performance of AMP predictors, we created a web server AMPBenchmark for fair benchmarking of AMP prediction models. Similarly to Kaggle, AMPBenchmark provides developers with public and private data sets for model training and validation that contain explicit and hidden data labels, respectively. The public data sets are the same samples that were used in the presented research. AMPBenchmark allows users to upload the prediction results for their AMP models, trained and benchmarked on the public data sets. It generates charts and tables comparing the performance of the uploaded architecture with those deposited in our database. The users can also upload prediction results for their AMP models, trained and benchmarked on the private data set, which is accessible after entering the e-mail address. The operator of AMPBenchmark will manually verify the results of the prediction and similarly provide charts and tables for comparative analysis.

Our study has also vital importance for the ongoing debate about the reproducibility crisis in science [56, 57]. In machine learning research, reproducibility means obtaining the same results to those presented in the original study using the same data and source code. Recently, Heil et al. [58] proposed three standards for computational reproducibility: bronze, silver and gold, reflecting the time needed to recreate research. The minimal and most time-consuming bronze standard requires: (i) data, (ii) models and (iii) source code to be published and downloadable. From the 26 models for AMP prediction that we considered, only eight met the minimal bronze standard (Table S2). This means that about 70% models represented non-reproducible work and consequently are unreliable. Interestingly, this number is very consistent with the survey published in the journal Nature [57] indicating that more than 70% researchers failed to reproduce other group’s experiments. Among the implemented models, five met the bronze standard: AmPEP, AmPEPpy, AmpGram, ampir and MACREL, and AmpScannerV2 was accessible upon request. These architectures, with the exception of A:AmPEP, also represent the top architectures investigated though A:AmpGram was clearly the most accurate and best at generalizing to other data sets.

The developers that do not reveal all the details necessary to recreate their models, not to mention reuse them, shoulder the blame for the lack of fair benchmarks for AMP prediction software. Consequently, progress in the field is slowed, mistrust to bioinformatics is spreading and resources that could have been allocated to other projects are wasted. Our study represents the first unbiased approach to compare models for AMP prediction, and moreover, we made reproducible another six model architectures for further research. In total, we built a staggering number of 660 machine learning models from 12 architectures. Therefore, being fully aware of the difficulty of the task, we highly recommend all researchers to embrace the notion of fair benchmarking and reproducibility by using AMPBenchmark web server and the recommendations provided by Heil et al. [58].

## Key Points

- We review the performance of existing machine learning models for identification of AMPs.
- Our benchmark highlight the major methodological flaw in the construction of benchmarks as the data sampling impacts the quality of AMP prediction.
- We propose a solution for fair benchmarking of AMP-predicting models.

## Code and software availability

The code necessary to reproduce the whole analysis is available at https://github.com/BioGenies/NegativeDatasets. All architectures are located at https://github.com/BioGenies/NegativeDatasetsArchitectures. AMPBenchmark web server is available at http://BioGenies.info/AMPBenchmark.

## Acknowledgments

This work was supported by the National Science Centre grant 2017/26/D/NZ8/00444 to P.G., the National Science Centre grant 2018/31/N/NZ2/01338 to K.S. and the National Science Centre grant 2019/35/N/NZ8/03366 to F.P. M. B. was supported by the Maria Zambrano grant funded by the European Union-NextGenerationEU. We thank Henrik Nielsen (Technical University of Denmark) and Simon Rasmussen (University of Copenhagen) for their critical review of the draft and fruitful discussions.

**Katarzyna Sidorczuk** received the M.Sc. degree in biotechnology from the University of Wroclaw, Poland, in 2019. She is currently pursuing the Ph.D. degree in biological sciences at the University of Wrocław. Her research focuses on bioinformatics and machine learning approaches for the analysis and prediction of peptide functions, protein targeting sequences, and bacterial adhesins.

**Przemysław Gagat** is an Assistant Professor in the Faculty of Biotechnology at the University of Wroclaw in Poland, where he received his B.Sc., (2006) M.Sc. (2008) and PhD (2014) in biotechnology. He has been involved in research projects in the field of bioinformatics, molecular phylogeny, molecular biology and evolution. His research interests include particularly endosymbioses of primary and complex plastids, and recently antimicrobial and anticancer peptides.

**Filip Pietluch** graduated with both a Masters and a Bachelors degree in Biotechnology from the University of Wrocław, Poland. He is currently a PhD student at the University of Wrocław, exploring phylogenetics and AMPs evolution. His main research topics are in computational and evolutionary biology.

**Jakub Kała** received the M. Sc. Eng. degree in Data Science from the Warsaw University of Technology in 2021. His main research focuses on applications of machine learning in protein function prediction.

**Dominik Rafacz** is a M.Sc. student with his major in Data Science at the Warsaw University of Technology. He has contributed to several projects related to predicting properties of proteins. He is an active open source developer, author and coauthor of several packages for the R programming language, including a framework for biological sequences storage and processing.

**Laura Bąkała** is another Data Science major at the Warsaw University of Technology, as well as a M.Sc. student. She has worked on a few open source R packages together with Dominik Rafacz, among those being the aforementioned framework for handling biological sequences.

**Jadwiga Słowik** received her master’s in data science from Warsaw University of Technology. She is passionate about software development and competitive programming.

**Rafał Kolenda** graduated in veterinary medicine from Wroclaw University of Environmental and Life Sciences, Poland, in 2013. He received Dr. med. vet. degree from Freie Universität, Berlin, Germany, in 2018. He is currently working as a research associate at the Department of Biochemistry and Molecular Biology of Wroclaw University of Environmental and Life Sciences, Poland. In his research, he uses molecular microbiology and omics techniques to investigate the pathogenic actions of bacteria. Moreover, he is involved in the development of new tools for *Escherichia coli* adhesiome analysis.

**Stefan Rödiger** is Privatdozent at the Brandenburg University of Technology. He studied pharamabiotechnology and forensics and received the doctoral degree from the Charité. His research includes personalized medicine, bioinformatics, and pharmacology.

**Legana C H W Fingerhut** graduated with a B.Sc degree in Ecology and Conservation and Zoology followed by a B.Biomed.Sc(Hons1) degree in Molecular Biology at James Cook University, Australia. She is currently a PhD candidate in bioinformatics examining antimicrobial peptide classification. She is the first author of the genome-wide antimicrobial peptide prediction R package, ampir.

**Ira R Cooke** received his PhD from the Australian National University in 2006 and is now a senior lecturer in bioinformatics at James Cook University, Australia. He is interested in the evolution and function of small secreted proteins in marine taxa, especially cephalopods and corals.

**Paweł Mackiewicz** is a professor at the University of Wroclaw in Poland. He is an author and co-author of more than 180 scientific publications. He is involved in many interdisciplinary research projects in the field of bioinformatics, genomics, phylogenetics and molecular evolution. His research interests focus on endosymbiosis, genetic code optimization and evolution, computer simulation of genome evolution, as well as phylogeny at the level of genes, proteins, genomes and organisms including ancient DNA samples.

**Michał Burdukiewicz** received Ph.D. in biochemistry from the University of Wrocław in 2019. He is currently working as a post-doc at the Institute of Biotechnology and Biomedicine at the Autonomous University of Barcelona and a research assistant in the Centre for Clinical Research at the Medical University of Białystok. His research interests cover machine learning applications in the functional analysis of peptides and proteins, focusing on amyloids. Moreover, he is co-developing tools for proteomics, mainly hydrogen-deuterium exchange monitored by mass spectrometry.

