## Supplementary material for "Benchmarks in antimicrobial peptide prediction are biased due to the selection of negative data": supp.pdf

### 1 Modifications to negative data sampling methods

In order to ensure fair benchmarking of negative data sampling methods, we introduced a few general changes. All sampling methods were run on the same negative data set created from sequences downloaded from UniProt [1]; CS-AMPpred originally used sequences from PDB database [2]. For SM:AMAP, SM:dbAMP, SM:CS-AMPpred, SM:Witten&Witten and SM:Gabere&Noble, we added a step removing sequences with ambiguous letters or non-standard amino acids because some model architectures do not handle such characters.

While filtering with keywords, we used only those present on the UniProt keyword list as identifiers to allow automatic filtering of sequences in the file with annotations downloaded from UniProt [1]. Consequently, some keywords were changed to match those present on the identifiers' list, such as: 'microbicidal' to 'antimicrobial' for SM:AMPlify, 'excreted' to 'secreted' for SM:AmpGram and SM:AMPScanner V2, 'antifungal' to 'fungicide' for SM:AmpGram and SM:AMPlify, 'toxic' to 'toxin' for SM:dbAMP, 'membrane' to 'transmembrane' for SM:dbAMP, 'secretory' to 'secreted' for SM:dbAMP. Some sampling methods were also modified by not including keywords absent from the list of UniProt identifiers and without any equivalents, such as: 'anticancer' for SM:dbAMP and SM:AMPlify; 'effector' for SM:AMPScanner V2; and 'antiparasitic', 'antimalarial', 'antiprotist', 'cathelicidin' and 'histatin' for SM:AMPlify.

Further changes were necessitated by the input data requirements. The simplest sampling methods demanded only a set of sequences with UniProt annotations whereas others depended on the positive data set, e.g. to ensure equal set size and/or equal sequence length distribution. For most methods: SM:Wang et al., SM:AmpGram, SM:Witten&Witten,

SM:AMPScanner V2, SM:Gabere&Noble and SM:AMAP, we ran negative data sampling separately for training and benchmarking using corresponding positive data sets. To prevent information leakage in methods that needed only sequences with annotations: SM:iAMP-2L, SM:dbAMP, i.e. did not depend on the positive data set, we generated single data sets and then split them into the training and benchmark sample.

We also introduced a few method-specific changes. In the case of SM:ampir-mature, we generated a negative data set and then divided it for training and benchmarking. Next, to ensure that the positive and negative set do not overlap, we filtered out sequences from the negative set using sequences from the corresponding positive samples. To ensure equal positive and negative set size for SM:CS-AMPpred, we used the number of sequences in the positive set to create a single negative sample and then split it for training and benchmarking.

Additionally, a few discrepancies occurred during the implementation of some sampling methods. First, to obtain equal length distribution in the positive and negative data set for SM:AMPlify, non-AMP sequences were generated from peptide/protein fragment or fragments instead of being selected from whole peptides/proteins. Second, for SM:AmpGram, we filtered out secretory proteins using the keyword 'secreted'. Third, for SM:AMPScanner V2 the step of CD-HIT reduction was omitted.

### 2 Modifications to model architectures

For all models requiring calculation of PseAAC, we used the implementation available in protr R package [3]. In the case of A:AMAP, we assumed default SVM hyperparameters; they were not provided by the authors. PseKRAAC in A:DeepAmPEP30 were calculated assuming k=1 and the default values of lambda and gap; they were not provided in the article. The architecture description of DeepAmPEP30 indicated that there is no padding both in the pooling and convolution layer. However, in the schema, the dimensionality of vector output does not change and therefore we did apply padding. AmpGram and ampir were modified to handle sequences shorter than 10 amino acids. In the case of AmpGram, we created an implementation that worked on 5-mers instead of 10-mers. For ampir, we changed the minimum sequence length to 5 after communication with the authors. The full description of changes is provided in the Table S3.

### 3 Tables

Table S1: Thresholds used for CD-HIT homology reduction of positive data set in the original papers describing negative data sampling methods we reimplemented.

| Method | Threshold |
|---|---|
| AMAP | 0.4[a] |
| AmpGram | 0.9 |
| ampir-mature | 0.9 |
| AMPlify | 1 |
| AMPScannerV2 | 0.9 |
| CS-AMPpred | - |
| dbAMP | 0.4 |
| Gabere&Noble | 0.9 |
| iAMP-2L | 0.4[b] |
| Wang et al. | 0.7 |
| WittenWitten | - |

[a]Homology partitioning at 40% for cross-validation
[b]Reduction performed only for selected subsets

Table S2: List of considered negative data sampling methods and architectures for AMP prediction. Models fulfilling the minimal standard for computational reproducibility according to [4] are marked with asterisk. SM - negative data sampling method.

| Software | Implemented SM | SM comment | Implemented Architecture | Architecture comment | Reference |
|---|---|---|---|---|---|
| ACEP* | No | Data set from AMPScanner V2 | No | Requires generation of PSSMs | [5] |
| AMAP | Yes | | Yes | | [6] |
| amPEP* | No | | Yes | | [7] |
| AmPEPpy* | No | Data set from amPEP | Yes | | [8] |
| AMP-GAN | - | | No | Not enough information | [9] |
| AmpGram* | Yes | | Yes | | [10] |
| ampir* | Yes | Precursor SM was not used since the selected model architectures are designed for mature proteins, the size of the precursor sample (significantly imbalanced) would cause problems in the result analysis | Yes | | [11] |
| AMPlify* | Yes | | No | We were not able to install dependencies for old versions of tensorflow-gpu needed to run the available code | [12] |
| AMPScanner V2 | Yes | | Yes | | [13] |
| ANFIS | No | Requires usage of Phobius | No | Requires usage of Tango software | [14] |
| AntiBP2 | No | Unclear information about sequence processing | No | Not enough information about SVM parameters | [15] |
| CAMPR3 | No | Uses experimentally proven non-AMPs and randomly generated sequences | No | Not enough information about generation of features | [16] |
| ClassAMP | No | Unclear information about acquisition of sequences | No | Link to scripts for feature selection does not work | [17] |
| CS-AMPpred | Yes | Added a step selecting proteins that contain only standard amino acids | Yes | | [18] |
| dbAMP | Yes | Added a step selecting proteins that contain only standard amino acids | No | Not enough information about feature selection | [19] |
| Deep-AmPEP30 | No | Due to the presence of longer sequences in the positive data set it is impossible to follow all steps of this SM | Yes | Model considers only short-length (<= 30 amino acid) AMPs | [20] |
| Gabere&Noble | Yes | Only DAMPD negative data set, added a step selecting proteins that contain only standard amino acids. Modified version of APD negative data set (added CD-HIT step) is used by AMAP | - | | [21] |
| iAMP-2L | Yes | | First model | The first model is responsible for AMP/nonAMP classification | [22] |
| IAMPE | No | Unclear information about sequence source | No | Requires a lot of work, including manual rewriting tables | [23] |
| iAMPpred | No | Multiclass data sets | No | Requires usage of Tango software | [24] |
| Maccari et. al | No | Unclear information about partitioning of the space into alpha and non-alpha peptides | No | Not enough information about algorithm parameters | [25] |
| MACREL* | No | Data set from amPEP | Yes | | [26] |
| MAMPs-pred | No | Uses Pfam families | No | Uses SVM-Prot for feature selection | [27] |
| MLAMP | No | Data set from iAMP-2L | First model | The first model is responsible for AMP/nonAMP classification | [28] |
| SVM-LZ | No | Data set from Wang et al. | Yes | | [29] |
| Wang et al. | Yes | Implemented only second data set | No | | [30] |
| Witten&Witten* | Yes | Added a step selecting proteins that contain only standard amino acids | No | Architecture based on MIC values (regression problem) | [31] |

Table S3: List of modifications to implemented architectures. Code sources marked with asteriks were not used during implementations.

| Architecture | Modifications | Code source |
|---|---|---|
| AMAP | Written based on the information found in the article; assumed default SVM parameters (not provided in a paper) | |
| AmPEP | Written in R based on the information found in the article as we were not able to run MATLAB version; used 23 distribution features selected by authors | https://sourceforge.net/projects/axpep/files/AmPEP_MATLAB_code* |
| AmPEPpy | Feature selection was not performed because (i) random forest itself performs regularization that should be sufficient for such number of features, (ii) authors did not show a significant gain in the performance using feature selection, (iii) authors indicated only 3 features out of 105 that were not significant for the model performance | https://github.com/tlawrence3/amPEPpy |
| AmpGram | Changed length of analysed k-mers (from 10 to 5) to handle sequences shorter than 10. | https://github.com/michbur/AmpGram-analysis |
| ampir | Changed minimal sequence length to 5 to handle sequences shorter than 10 | https://github.com/Legana/ampir; https://github.com/Legana/AMP_pub |
| AMPScanner V2 | Changed sequence maximum length from 200 to the length of the longest sequence in the training data set | Provided by the authors on the request |
| CS-AMPpred | Written based on the information found in the article | |
| Deep-AmPEP30 | Written based on the information found in the article; PseKRAAC were calculated assuming k=1; applied padding in pooling and convolutional layer (description did not indicate it but the schema showed that dimensionality of vector output does not change); the code provided by the authors was found at the end of our analyses | https://sourceforge.net/projects/axpep/files/Deep-AmPEP30_datasets* |
| iAMP-2L | Written based on the information found in the article; implemented only the first model responsible for discrimination between AMPs and non-AMPs; PseAAC were calculated using APAAC function from the protr R package | |
| MACREL | Used the available code without modifications | https://github.com/BigDataBiology/macrel |
| MLAMP | Written based on the information found in the article; implemented only the first model responsible for discrimination between AMPs and non-AMPs | |
| SVM-LZ | Written based on the information found in the article | |

Table S4: Number of sequences in the training data sets: the positive sample and five replicates of a given negative sampling method.

| Data set | 1 | 2 | 3 | 4 | 5 |
|---|---|---|---|---|---|
| Positive | 4151 | | | | |
| TSM:AMAP | 3999 | 4005 | 3998 | 4037 | 4007 |
| TSM:AmpGram | 4151 | 4151 | 4151 | 4151 | 4151 |
| TSM:ampir-mature | 2856 | 2854 | 2859 | 2863 | 2866 |
| TSM:AMPlify | 4151 | 4151 | 4151 | 4151 | 4151 |
| TSM:AMPScannerV2 | 4151 | 4151 | 4151 | 4151 | 4151 |
| TSM:CS-AMPPred | 4151 | 4151 | 4151 | 4151 | 4151 |
| TSM:dbAMP | 4112 | 4112 | 4112 | 4112 | 4112 |
| TSM:Gabere&Noble | 24906 | 24906 | 24906 | 24906 | 24906 |
| TSM:iAMP-2L | 5862 | 5862 | 5862 | 5862 | 5862 |
| TSM:Wang et. al | 8316 | 8304 | 8393 | 8342 | 8430 |
| TSM:Witten&Witten | 4151 | 4151 | 4151 | 4151 | 4151 |

Table S5: Number of sequences in the benchmark data sets: the positive sample and five replicates of a given negative sampling method.

| Data set | 1 | 2 | 3 | 4 | 5 |
|---|---|---|---|---|---|
| Positive | 1039 | | | | |
| BSM:AMAP | 1470 | 1472 | 1478 | 1519 | 1474 |
| BSM:AmpGram | 1039 | 1039 | 1039 | 1039 | 1039 |
| BSM:ampir-mature | 750 | 747 | 749 | 744 | 751 |
| BSM:AMPlify | 1039 | 1039 | 1039 | 1039 | 1039 |
| BSM:AMPScannerV2 | 1039 | 1039 | 1039 | 1039 | 1039 |
| BSM:CS-AMPPred | 1039 | 1039 | 1039 | 1039 | 1039 |
| BSM:dbAMP | 1028 | 1028 | 1028 | 1028 | 1028 |
| BSM:Gabere&Noble | 6234 | 6234 | 6234 | 6234 | 6234 |
| BSM:iAMP-2L | 1466 | 1466 | 1466 | 1466 | 1466 |
| BSM:Wang et. al | 8545 | 8546 | 8420 | 8491 | 8513 |
| BSM:Witten&Witten | 1039 | 1039 | 1039 | 1039 | 1039 |

Table S6: Architecture performance depending on the negative data sampling method used for training and benchmarking.

| Architecture | Mean AUC[1] | Mean AUC[2] |
|---|---|---|
| AMAP | 0.90 | 0.82 |
| AmPEP | 0.62 | 0.69 |
| AmPEPpy | 0.95 | 0.86 |
| AmpGram | 0.96 | 0.92 |
| Ampir | 0.97 | 0.87 |
| AMPScannerV2 | 0.97 | 0.86 |
| CS-AMPPred | 0.87 | 0.82 |
| Deep-AmPEP30 | 0.91 | 0.78 |
| iAMP-2L | 0.63 | 0.66 |
| MACREL | 0.96 | 0.89 |
| MLAMP | 0.97 | 0.85 |
| SVM-LZ | 0.80 | 0.77 |

[1]Mean AUC calculated for models trained and benchmarked on sets produced by the same sampling methods
[2]Mean AUC calculated for models trained and benchmarked on sets produced by different sampling methods

Table S7: Kruskal-Wallis test with Bonferroni correction for models trained and benchmarked on sets produced by the same and different negative data sampling methods.

| Architecture | Bonferroni corrected $p$-value |
|---|---|
| AMAP | **2.11e-09** |
| AmPEP | **1.54e-06** |
| AmPEPpy | **2.59e-17** |
| AmpGram | **4.29e-10** |
| Ampir | **4.72e-18** |
| AMPScannerV2 | **8.29e-18** |
| CS-AMPPred | **3.85e-06** |
| Deep-AmPEP30 | **1.09e-16** |
| iAMP-2L | 1 |
| MACREL | **1.47e-16** |
| MLAMP | **3.33e-17** |
| SVM-LZ | 0.141 |

Table S8: Pairwise Wilcoxon test for paired samples between groups of model architectures.

| Architecture | AMAP | AmPEP | AmPEPpy | AmpGram | Ampir | AMPScannerV2 | CS-AMPPred | Deep-AmPEP30 | iAMP-2L | MACREL | MLAMP |
|---|---|---|---|---|---|---|---|---|---|---|---|
| AmPEP | **5.75e-15** | | | | | | | | | | |
| AmPEPpy | **8.38e-11** | **2.89e-19** | | | | | | | | | |
| AmpGram | **9.02e-20** | **9.02e-20** | **9.89e-12** | | | | | | | | |
| Ampir | **6.56e-16** | **2.1e-19** | **0.00116** | **0.0025** | | | | | | | |
| AMPScannerV2 | **8.78e-13** | **1.34e-16** | 1 | **1.13e-05** | **0.0166** | | | | | | |
| CS-AMPPred | 1 | **8.96e-18** | **3.16e-08** | **9.02e-20** | **5.75e-11** | **2.96e-06** | | | | | |
| Deep-AmPEP30 | **1.05e-06** | **4.14e-07** | **2.27e-16** | **2.75e-19** | **3.44e-19** | **1.65e-17** | **0.0182** | | | | |
| iAMP-2L | **8.03e-16** | 1 | **1.14e-16** | **1.85e-19** | **9.85e-18** | **4.63e-17** | **3.26e-16** | **6.29e-09** | | | |
| MACREL | **8.59e-16** | **9.02e-20** | **5.08e-17** | **0.0016** | **0.0231** | **0.00292** | **9.82e-16** | **6.71e-17** | **2.3e-17** | | |
| MLAMP | **4.82e-07** | **2.45e-14** | 1 | **0.000263** | 0.0958 | 1 | 0.0868 | **4.18e-16** | **3.33e-16** | 0.132 | |
| SVM-LZ | **5.67e-06** | **4.17e-11** | **6.14e-15** | **9.02e-20** | **9.08e-15** | **1.07e-11** | **7.21e-09** | 0.684 | **1.04e-09** | **3.76e-17** | **2.38e-06** |

Table S9: Pairwise Wilcoxon test for paired samples between groups of training data sampling methods (TSM).

| TSM | AMAP | AmpGram | ampir-mature | AMPlify | AMPScannerV2 | CS-AMPPred | dbAMP | Gabere&Noble | iAMP-2L | Wang et al. |
|---|---|---|---|---|---|---|---|---|---|---|
| AmpGram | 1 | | | | | | | | | |
| ampir-mature | 0.12 | **8.42e-11** | | | | | | | | |
| AMPlify | 1 | **0.00114** | **1.81e-10** | | | | | | | |
| AMPScannerV2 | 1 | **8.79e-06** | **7.15e-11** | 1 | | | | | | |
| CS-AMPPred | **2.42e-09** | **0.00101** | 1 | **0.00289** | **0.00255** | | | | | |
| dbAMP | **3.78e-10** | **0.000481** | 1 | **0.00126** | **0.00125** | **0.00041** | | | | |
| Gabere&Noble | 1 | 1 | **1.86e-11** | 1 | 1 | **0.000754** | **0.000199** | | | |
| iAMP-2L | **9.27e-10** | **0.00111** | 1 | **0.00278** | **0.00267** | **0.00779** | 1 | **0.000264** | | |
| Wang et al. | 1 | 1 | **1.87e-10** | 1 | 0.5 | **0.000109** | **4.74e-05** | 1 | **8.91e-05** | |
| Witten&Witten | 1 | **1e-07** | **3.67e-06** | **0.000248** | **2.26e-06** | 0.403 | 0.244 | **2.77e-05** | 0.367 | **1.92e-07** |

Table S10: Pairwise Wilcoxon test for paired samples between groups of benchmark data sampling methods (BSM).

| BSM | AMAP | AmpGram | ampir-mature | AMPlify | AMPScannerV2 | CS-AMPPred | dbAMP | Gabere&Noble | iAMP-2L | Wang et al. |
|---|---|---|---|---|---|---|---|---|---|---|
| AmpGram | **7.3e-06** | | | | | | | | | |
| ampir-mature | **2.34e-18** | **1.25e-07** | | | | | | | | |
| AMPlify | **4.12e-09** | **1.98e-14** | **9.81e-06** | | | | | | | |
| AMPScannerV2 | **0.000563** | **1.73e-13** | **2.44e-08** | **9.55e-15** | | | | | | |
| CS-AMPPred | 1 | 0.652 | **9.37e-13** | 0.106 | 1 | | | | | |
| dbAMP | 1 | 0.151 | **3.45e-13** | **0.0141** | 0.347 | **1.4e-08** | | | | |
| Gabere&Noble | **0.000298** | **7.61e-14** | **3.03e-08** | **1.13e-14** | **0.0281** | 1 | 0.319 | | | |
| iAMP-2L | 1 | 0.401 | **4.43e-13** | 0.0517 | 0.864 | **0.000169** | **1.34e-06** | 0.817 | | |
| Wang et al. | **0.000956** | **8.1e-12** | **6.6e-12** | **1e-15** | **0.00024** | 1 | 0.903 | **5.2e-05** | 1 | |
| Witten&Witten | **0.00807** | **1.88e-14** | **1.04e-08** | **3.74e-15** | **4.53e-05** | 1 | 0.978 | **1.46e-07** | 1 | 1 |

Table S11: Mean standard deviation (SD) of AUC value for the five replicates of data sets.

| Architecture | Mean SD | Min SD | Max SD |
|---|---|---|---|
| AMAP | 0.00502 | 0.00099 | 0.01153 |
| AmPEP | 0.00813 | 0.00174 | 0.01789 |
| AmPEPpy | 0.00400 | 0.00010 | 0.01399 |
| AmpGram | 0.00353 | 0.00072 | 0.01414 |
| Ampir | 0.00370 | 0.00021 | 0.01460 |
| AMPScannerV2 | 0.01859 | 0.00058 | 0.09374 |
| CS-AMPPred | 0.00508 | 0.00143 | 0.01397 |
| Deep-AmPEP30 | 0.01440 | 0.00079 | 0.05347 |
| iAMP-2L | 0.03470 | 0.00082 | 0.09706 |
| MACREL | 0.00323 | 0.00031 | 0.01520 |
| MLAMP | 0.00428 | 0.00013 | 0.01407 |
| SVM-LZ | 0.00572 | 0.00099 | 0.01414 |

### 4 Figures

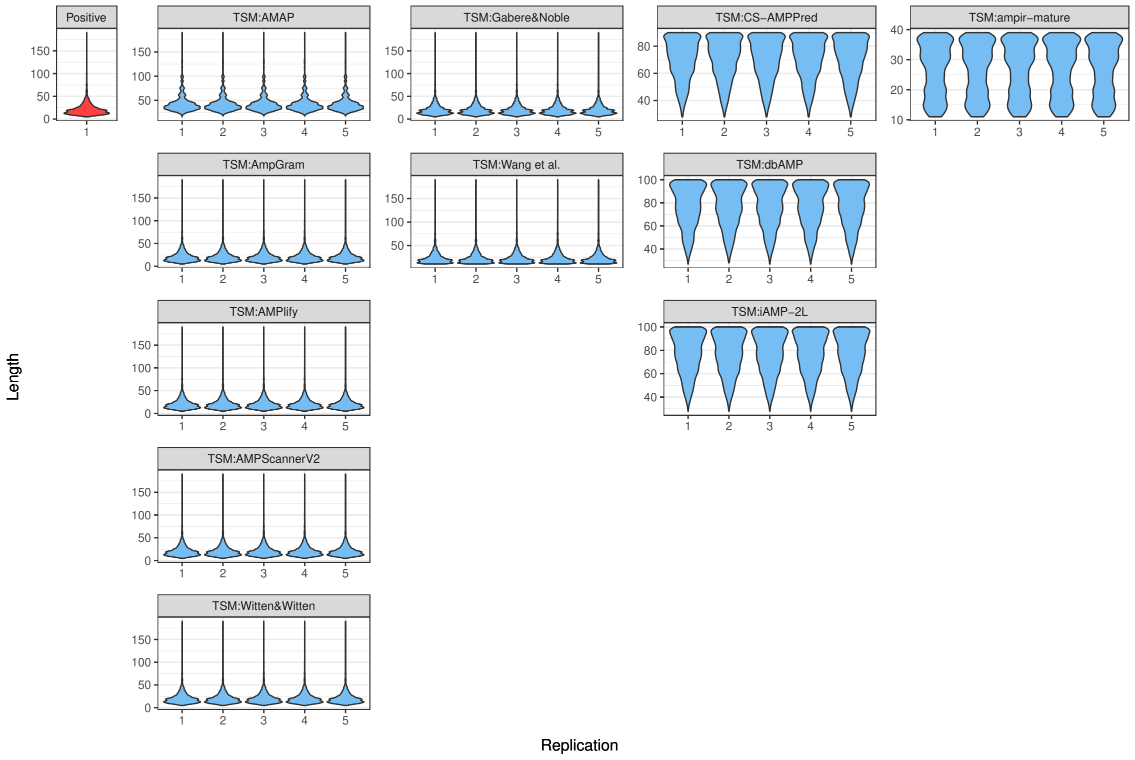

Figure S1: Length distribution of sequences in the positive and five replicates of the negative data sets.

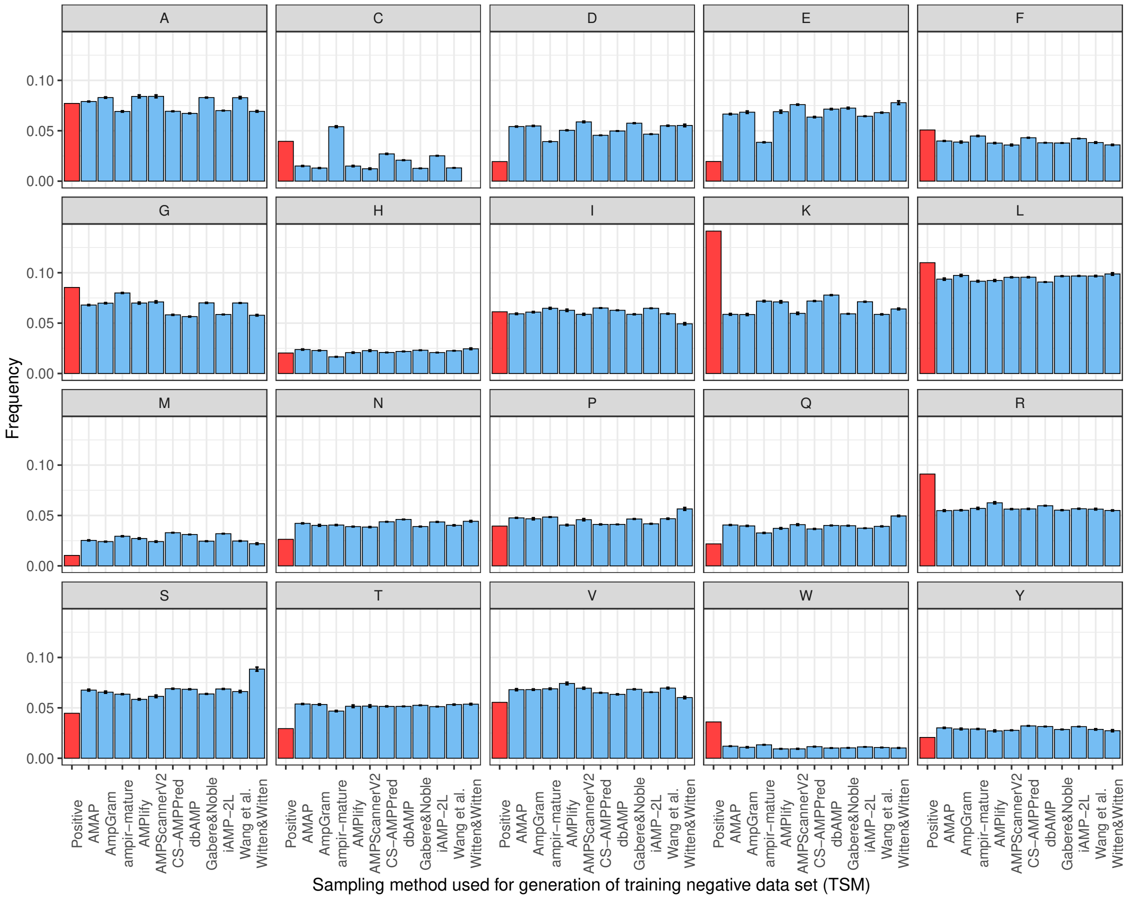

Figure S2: Amino acid composition of sequences in the positive and negative data sets. The global comparison of all sequences included in the positive sample and the negative ones indicated that AMPs are richer in positively charged amino acids: lysine (K) and arginine (R), and hydrophobic residues: tryptophan (W) and leucine (L). They were also abundant in cysteine (C), but not as much as the negative sets of ampir-mature; C is responsible for stabilization of motif and domain structure [32]. Interestingly, glycine, important for peptide conformational flexibility, was not as abundant in AMP compared to non-AMP sequences as previous studies indicated [10]. AMPs are also depleted in negatively charged amino acids: aspartate (D) and glutamate (E), and other hydrophilic ones such as: asparagine (N), glutamine (Q), serine (S), threonine (T) and tyrosine (Y). They are also poor in methionine (M) though methionine is a moderately hydrophobic amino acid that was shown to enhance antimicrobial properties at least for some peptides [33].

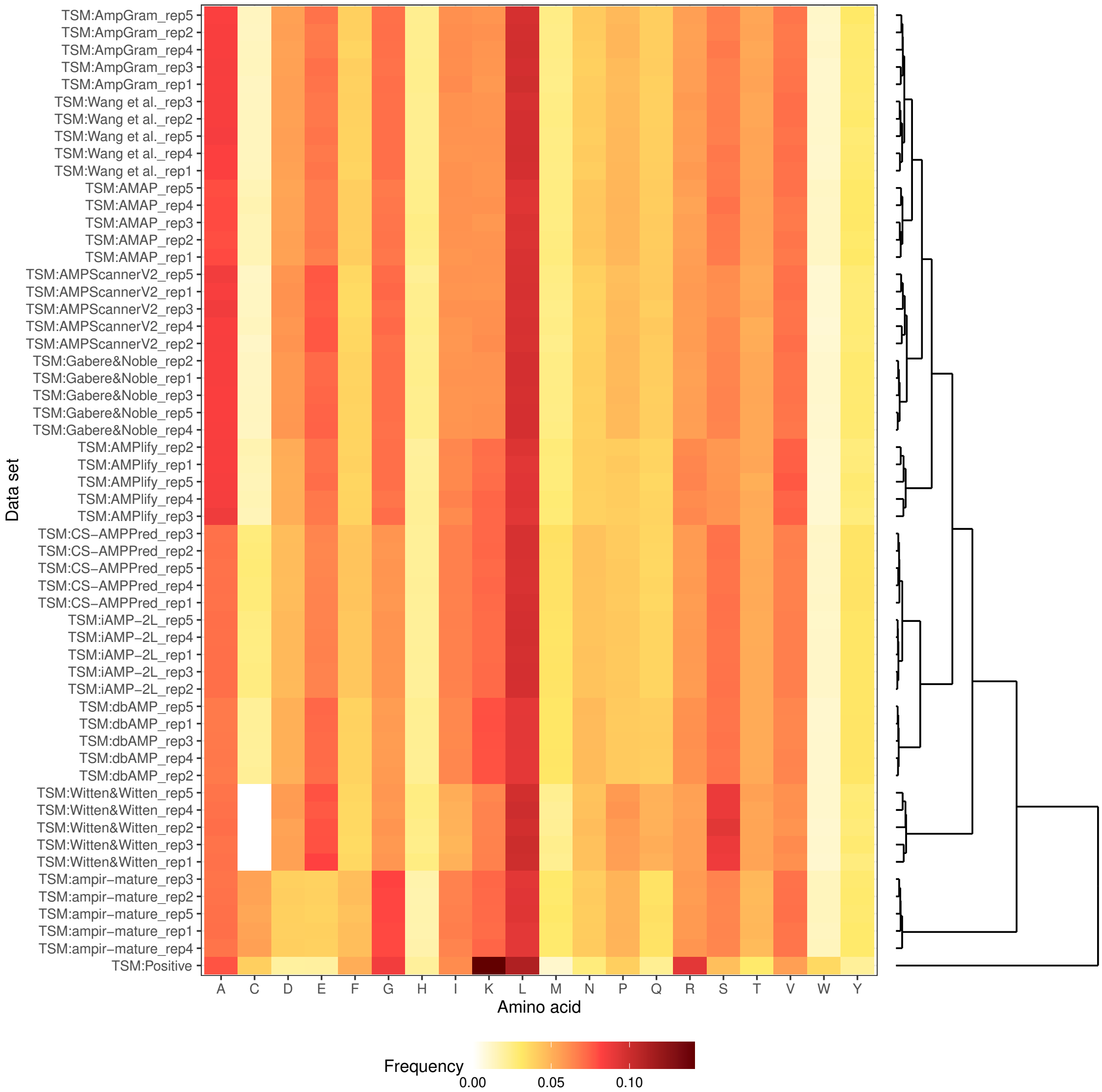

Figure S3: Hierarchical clustering using Euclidian distance performed on amino acid composition for sequences from the positive and five replicates of the negative data sets. The length of branches represents the similarity between the data sets, i.e. the shorter the branches the more similar they are. The x-axis of the heat map represents amino acids ordered alphabetically according to the one-letter code system. The y-axis represents the methods of negative data sampling (five replicates each) and the positive data set. The over- and under-representation of amino acids for each data set are indicated as shades of brown and yellow, respectively.

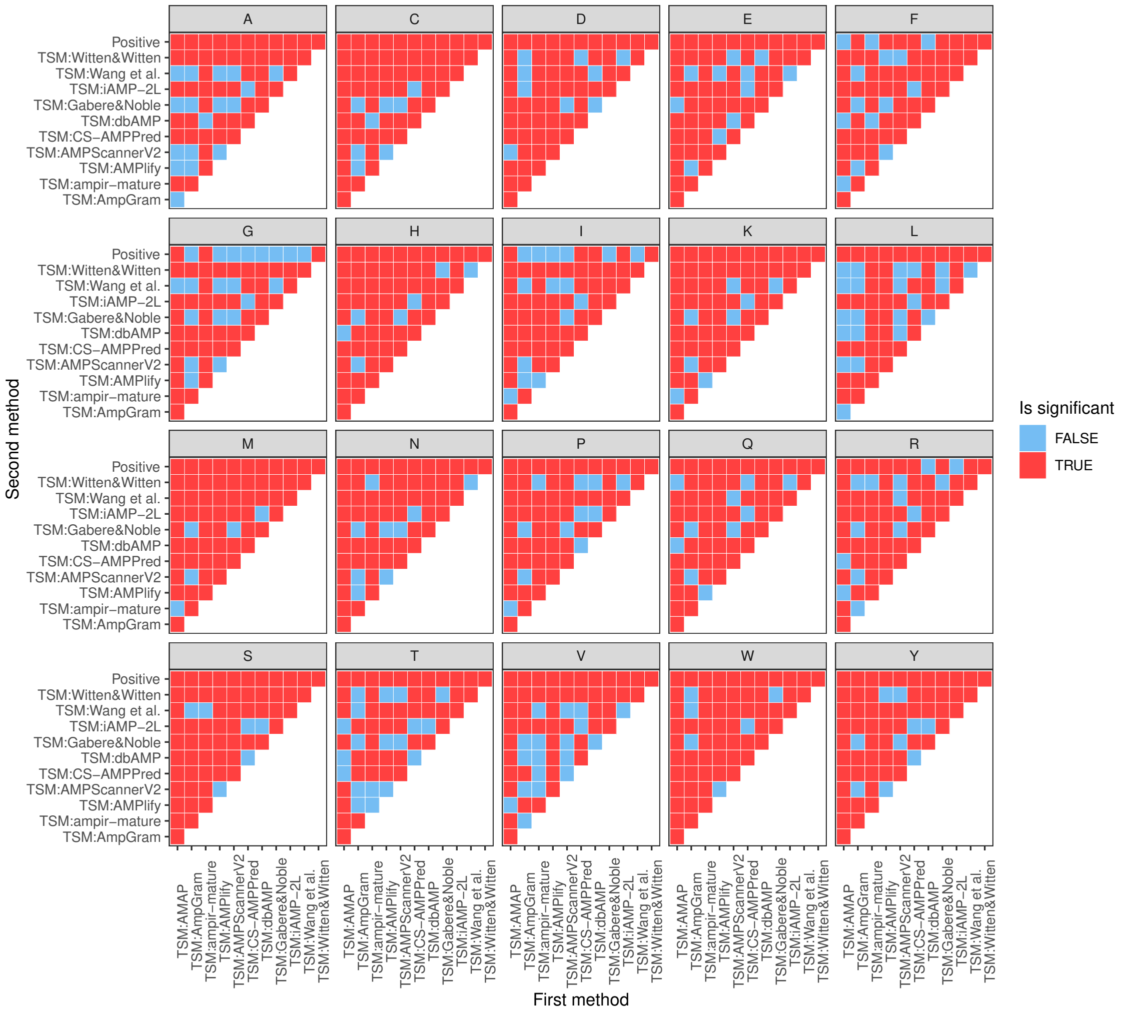

Figure S4: Mann-Whitney U test for the comparison of fractions of a given amino acid per peptide.

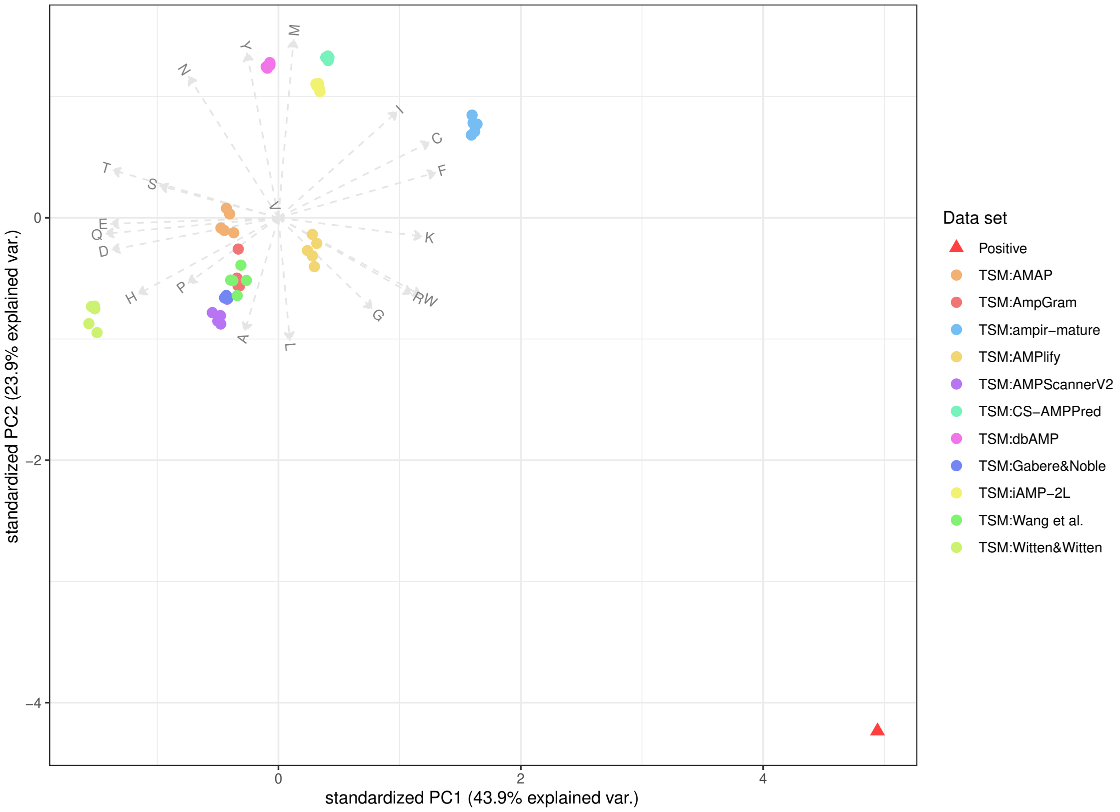

Figure S5: Principal component analysis performed on amino acid composition for sequences from the positive (red triangle) and negative data sets (colourful dots). Each dot represents a replicate of a given data set. The dots are coloured according to method of negative data sampling. The original variables, i.e. amino acid frequencies, are shown as vectors and letters according to the one-letter code name. The first (PC1) and second (PC2) principal component account for 67.8% of the variation in the data sets.

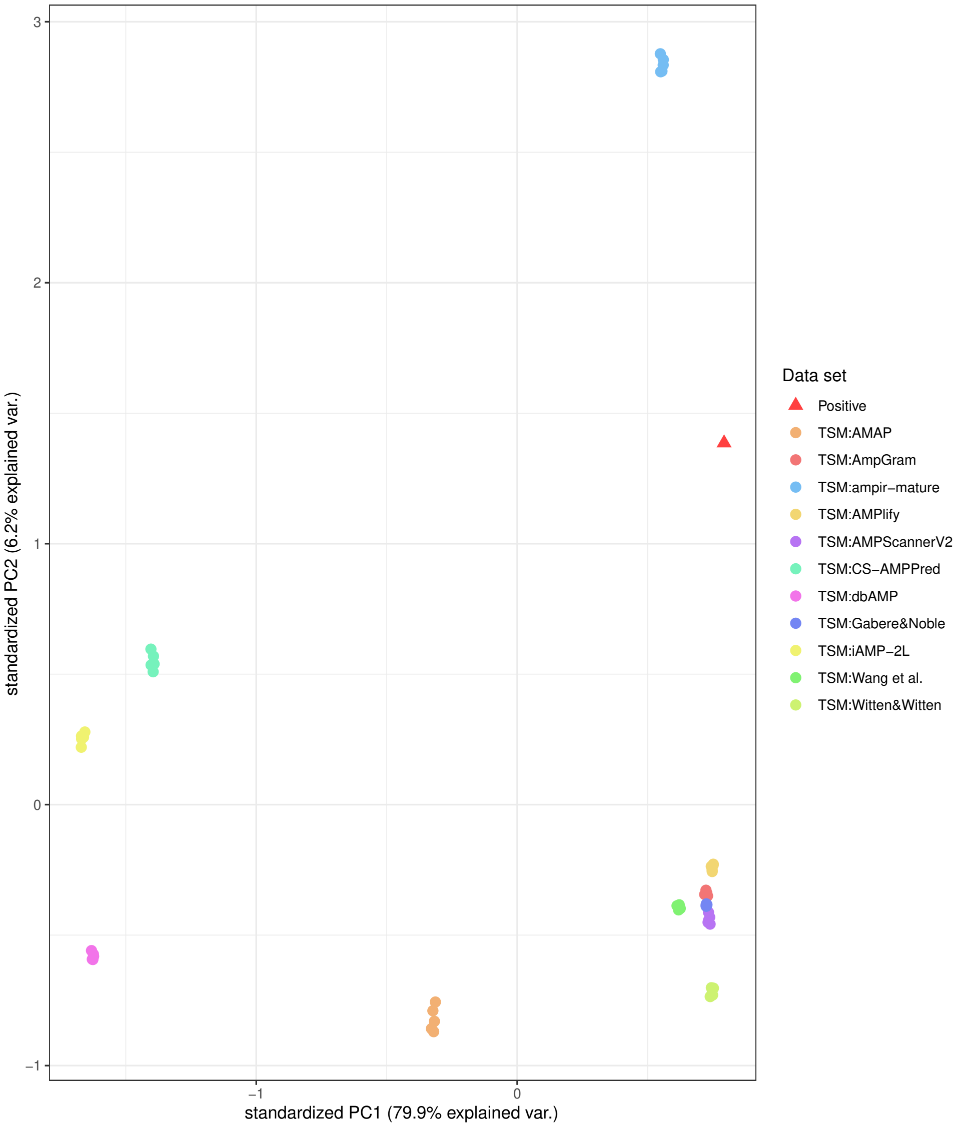

Figure S6: Principal component analysis performed on n-grams for sequences from the positive (red triangle) and negative data sets (colourful dots). N-grams are amino acid motifs of n elements. We performed the PCA on bigrams (n-gram of size 2) and trigrams (n-gram of size 3). For bigrams, we also considered n-grams with a gap length of 1, whereas the trigrams could contain only a single gap between the first and the second or the second and the third position. Each dot represents a replicate of a given data set. The dots are coloured according to method of negative data sampling. The sole first principal component (PC1) accounts for 79.9% of the variation in the data sets.

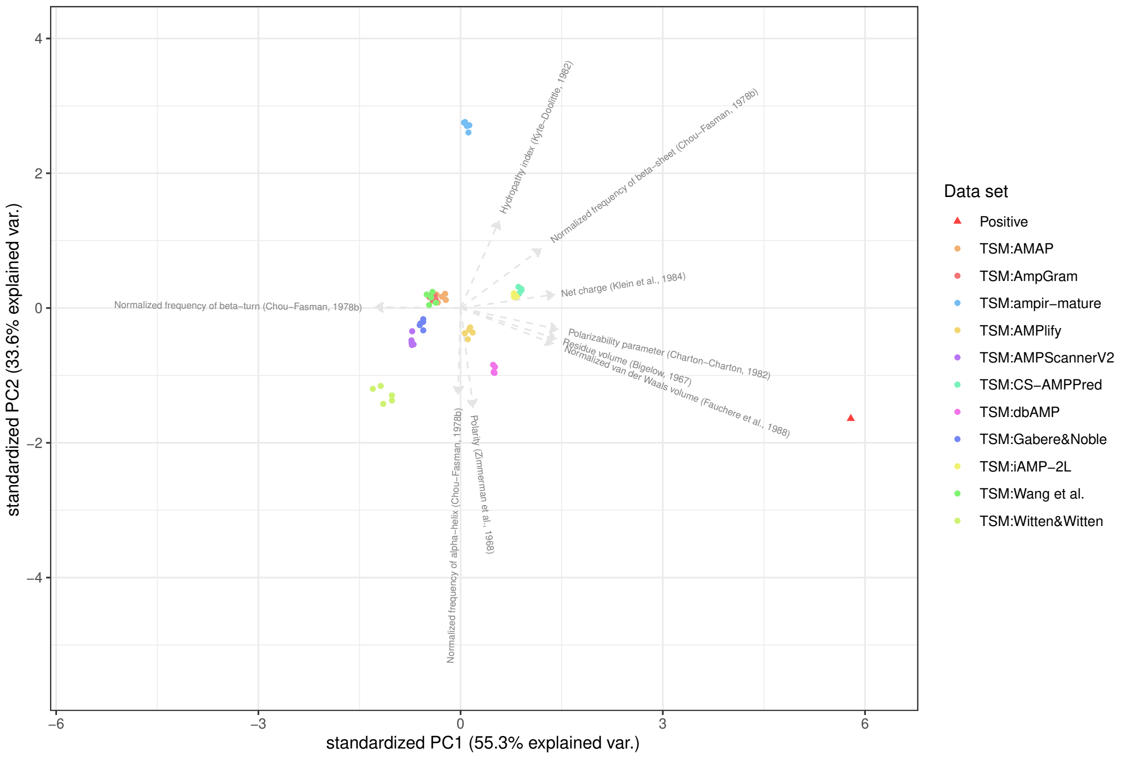

Figure S7: Principal component analysis performed on physicochemical properties for sequences from the positive (red triangle) and negative data sets (colourful dots). Each dot represents a replicate of a given data set. The dots are coloured according to method of negative data sampling. The original variables, i.e. physicochemical properties, are shown as vectors and appropriately named. The first (PC1) and second (PC2) principal component account for 88.9% of the variation in the data sets.

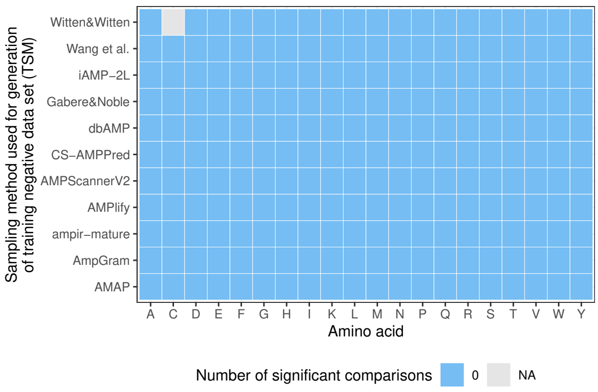

Figure S8: Mann-Whitney U test for the comparison of amino acid composition among the five replicates of each negative data sampling method.

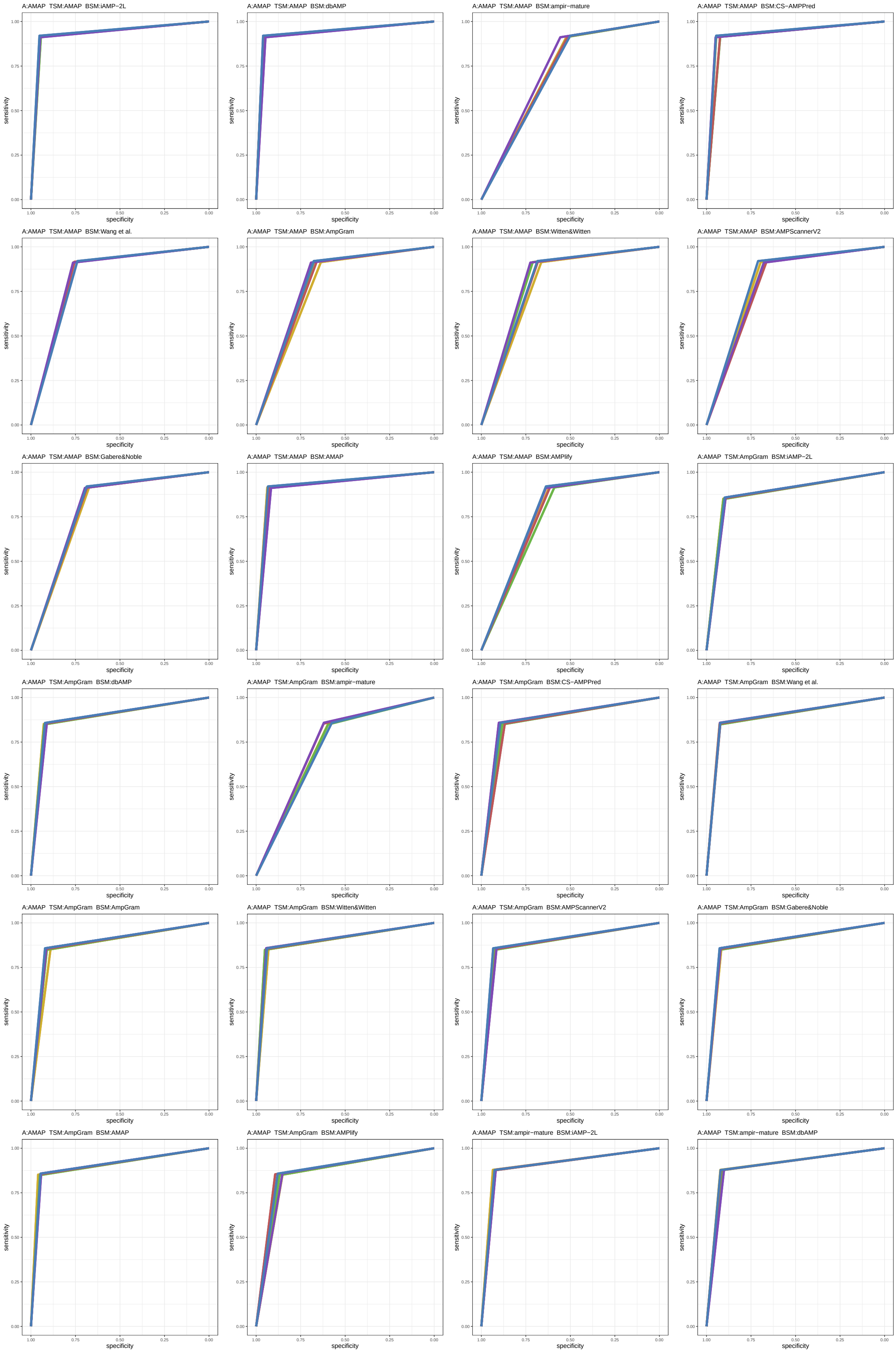

Figure S9: ROC curves 1-24 of 1452. Each subplot presents results for five replications indicated by different line colors.

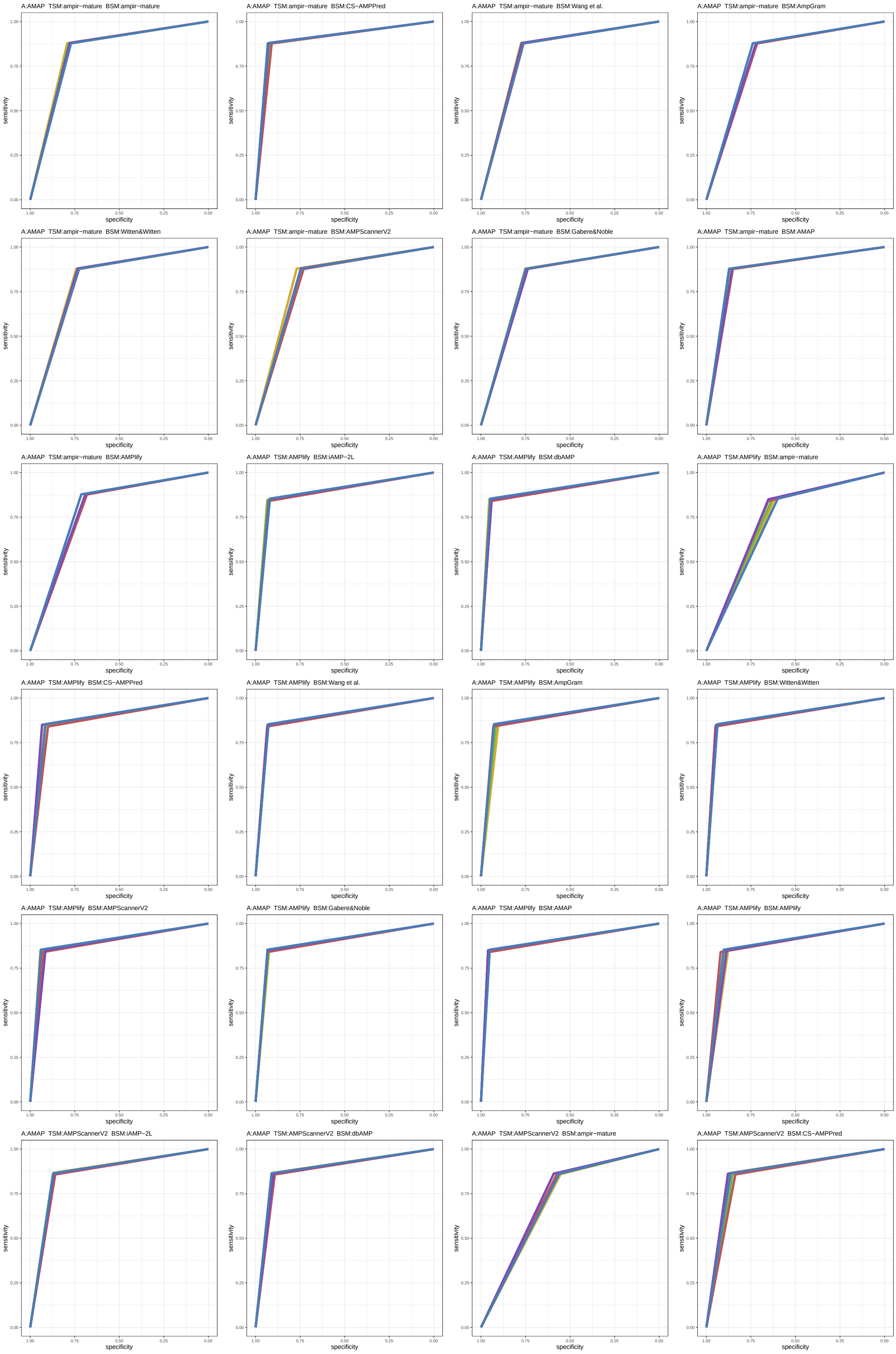

Figure S10: ROC curves 25-48 of 1452. Each subplot presents results for five replications indicated by different line colors.

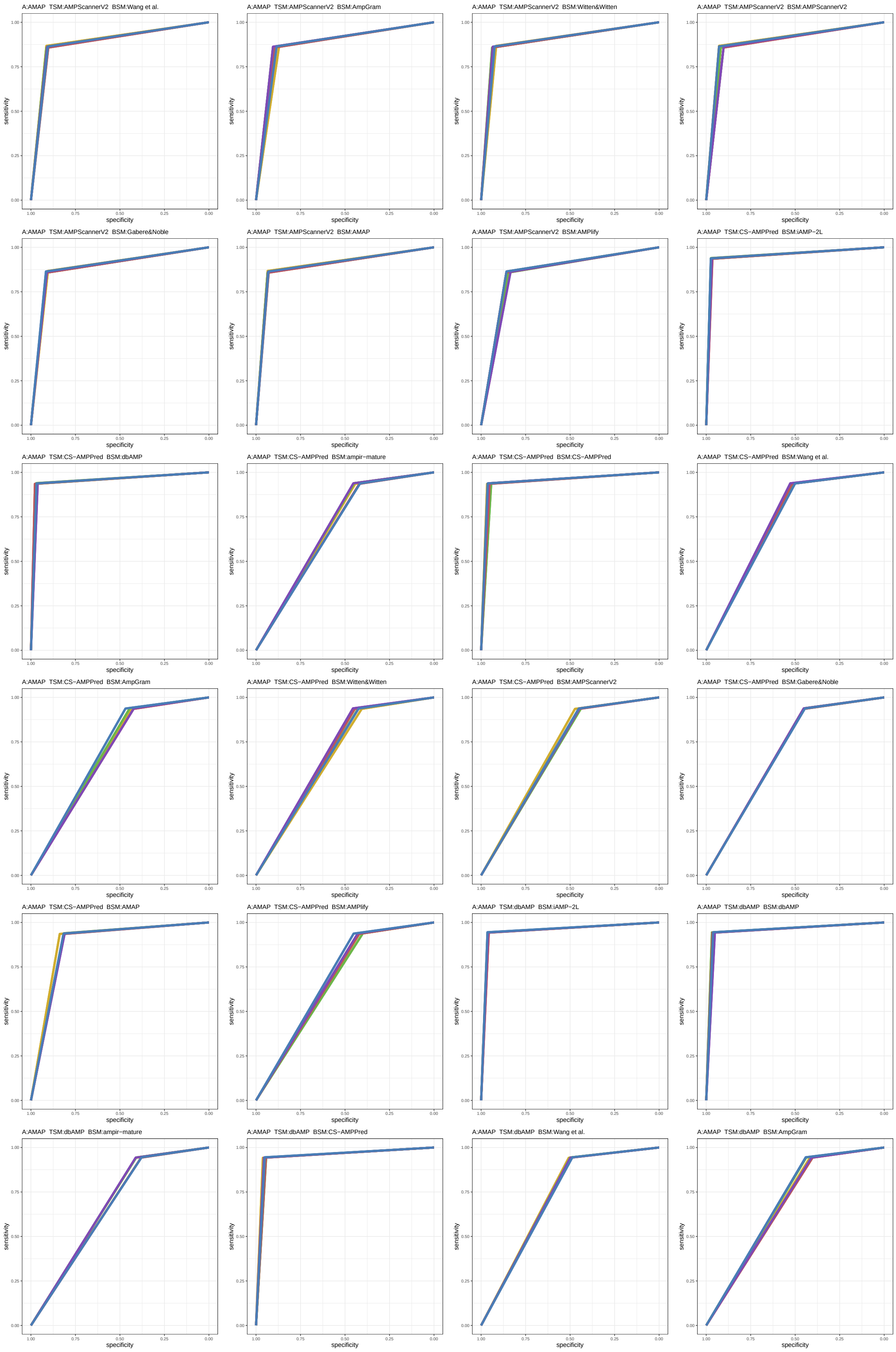

Figure S11: ROC curves 49-72 of 1452. Each subplot presents results for five replications indicated by different line colors.

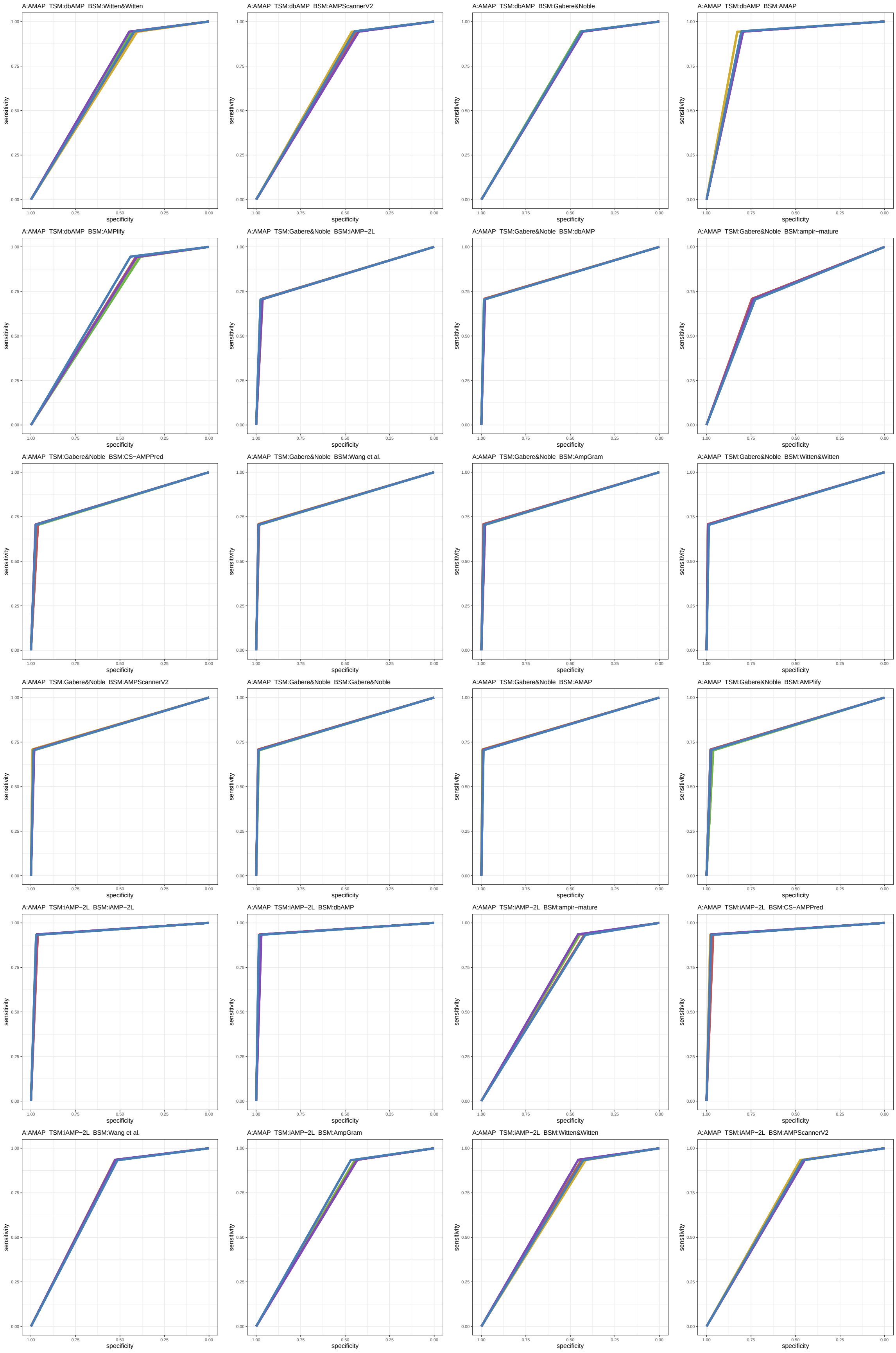

Figure S12: ROC curves 73-96 of 1452. Each subplot presents results for five replications indicated by different line colors.

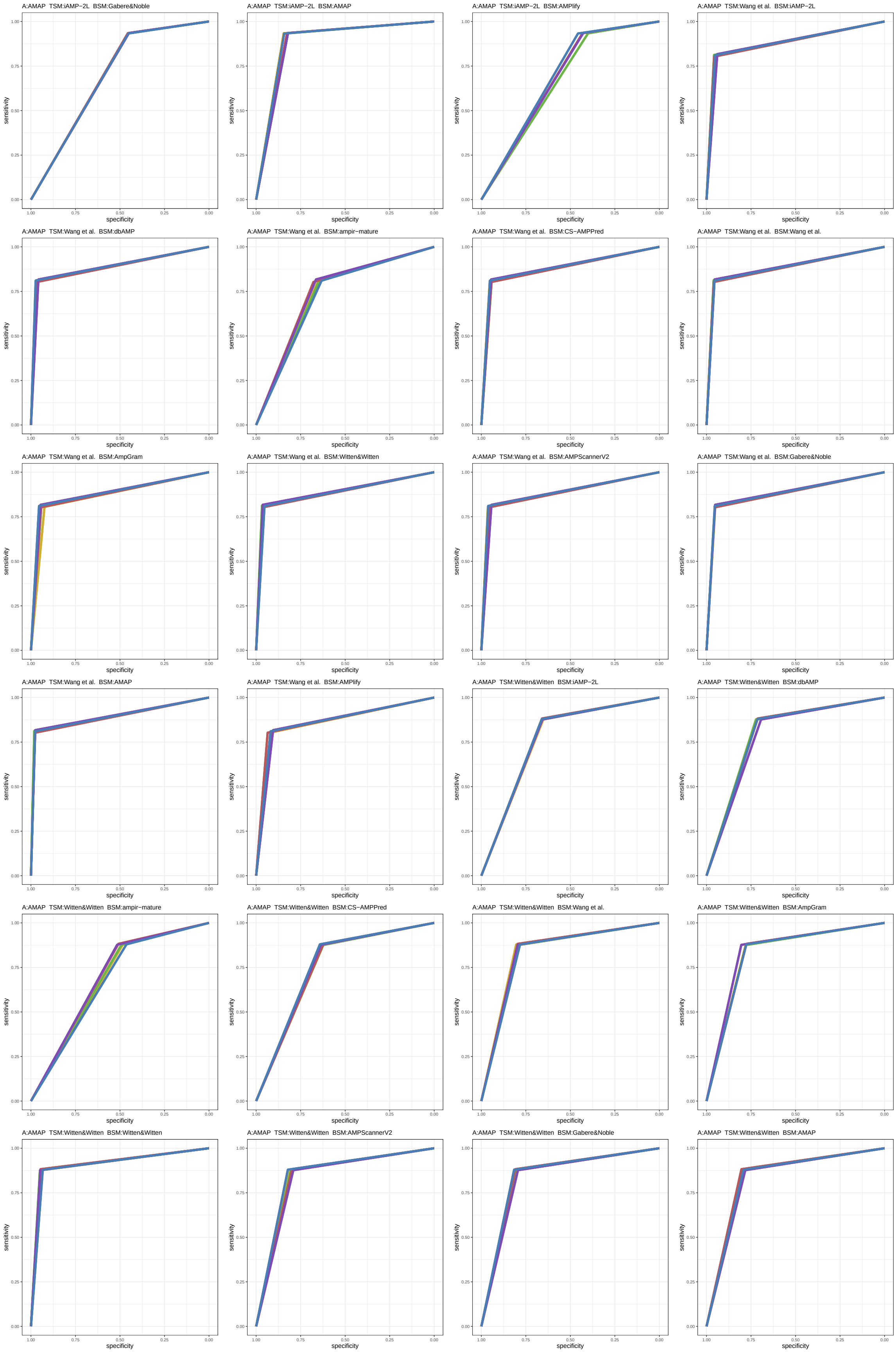

Figure S13: ROC curves 97-120 of 1452. Each subplot presents results for five replications indicated by different line colors.

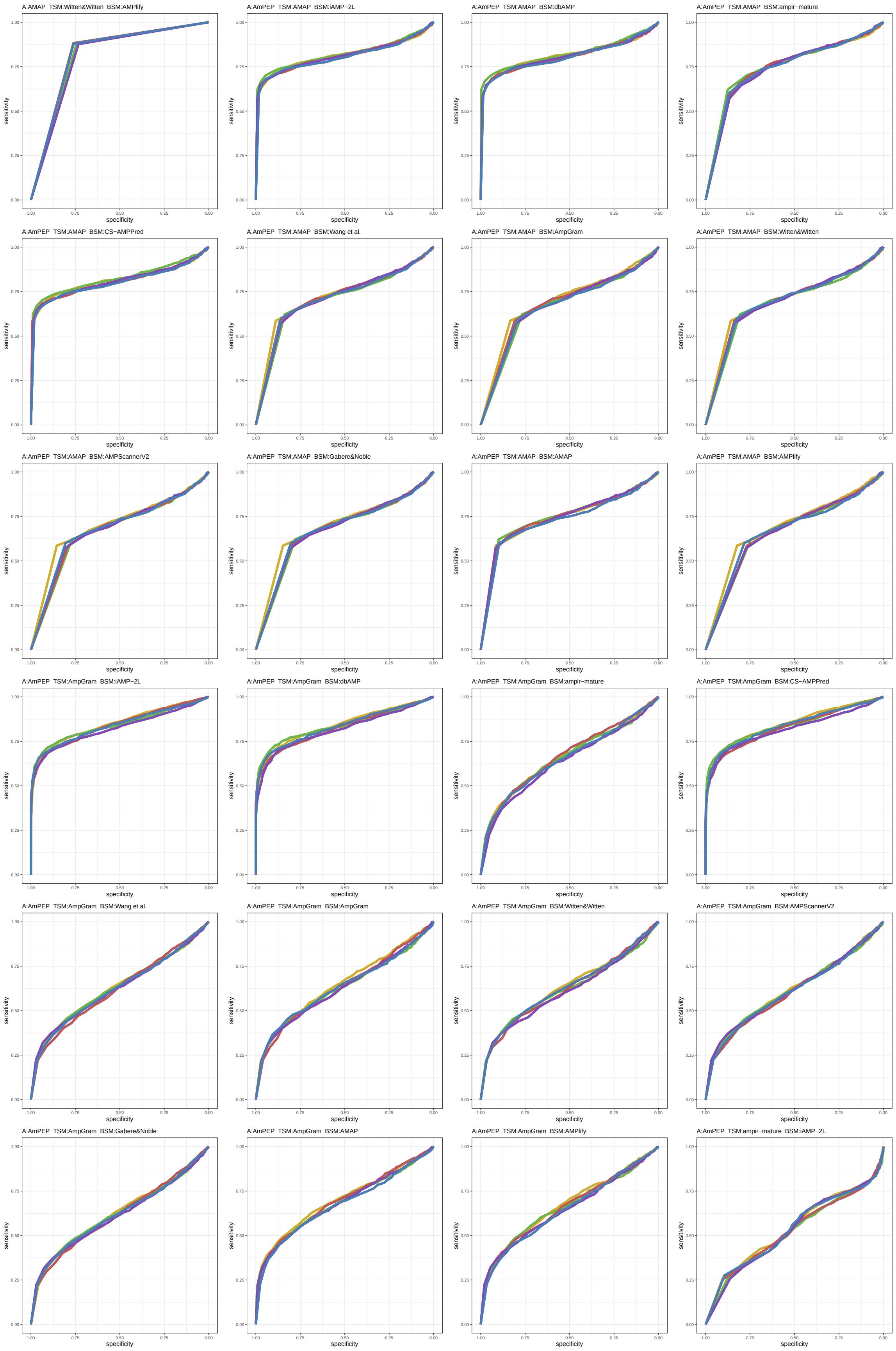

Figure S14: ROC curves 121-144 of 1452. Each subplot presents results for five replications indicated by different line colors.

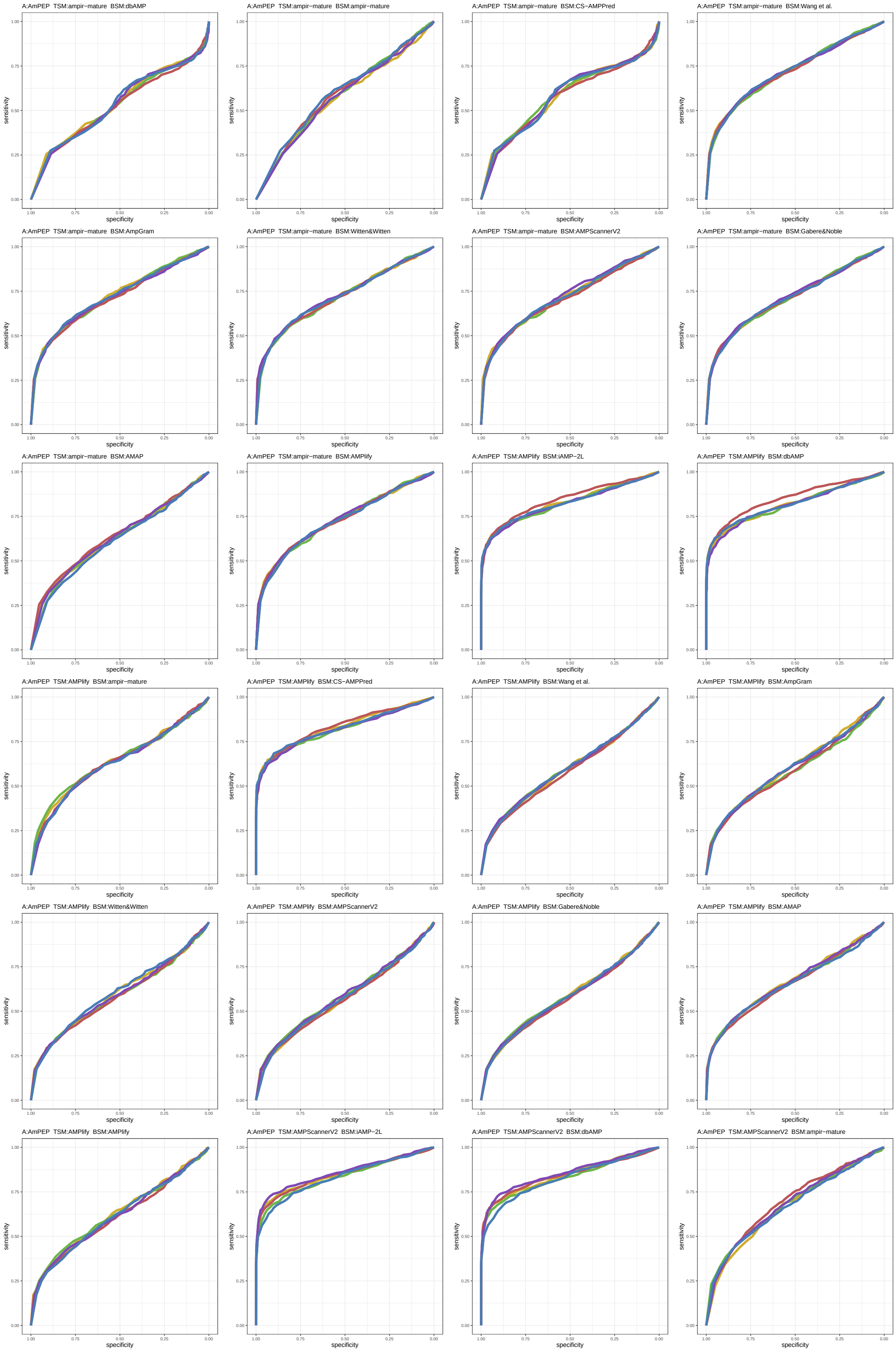

Figure S15: ROC curves 145-168 of 1452. Each subplot presents results for five replications indicated by different line colors.

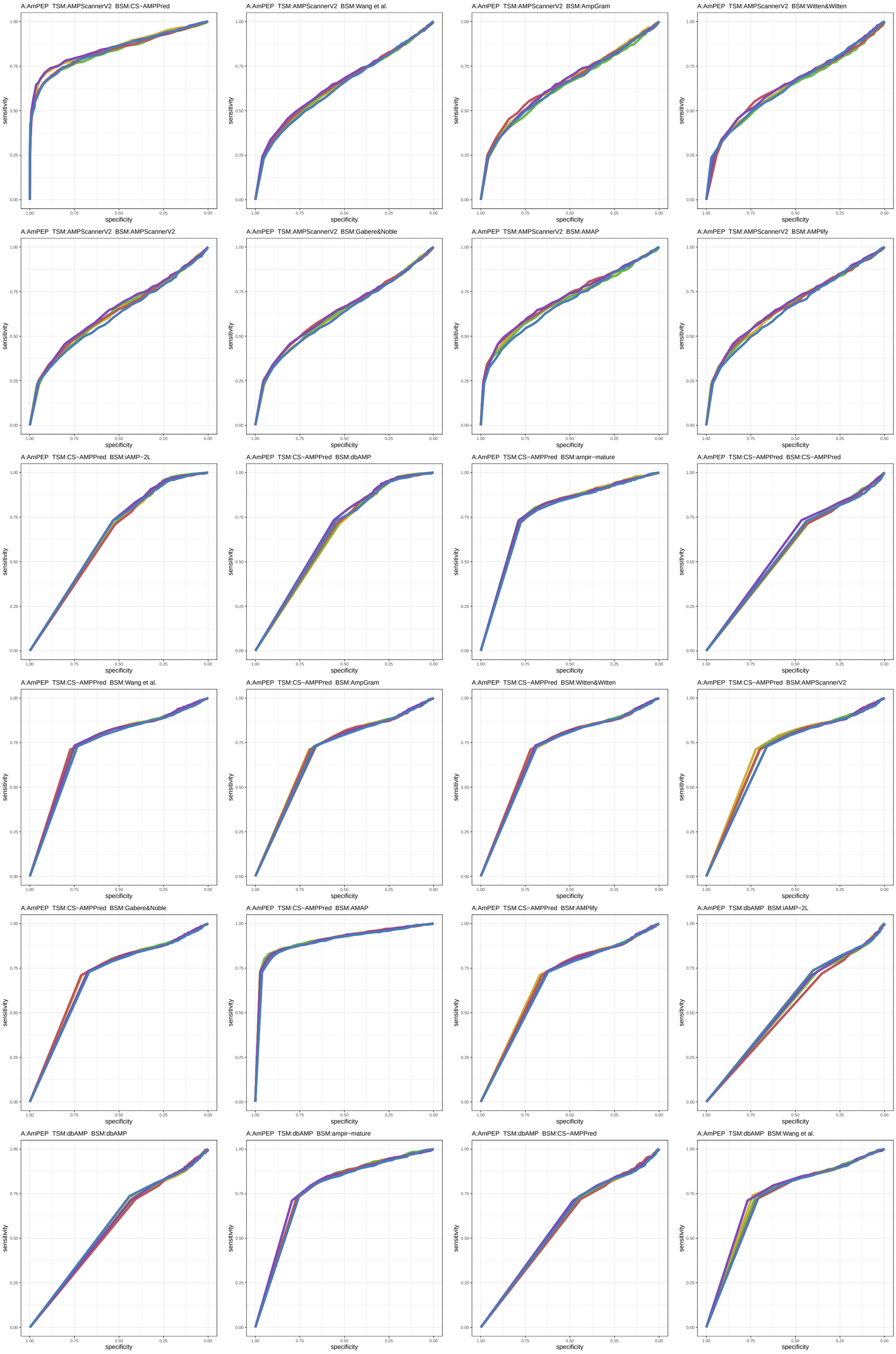

Figure S16: ROC curves 169-192 of 1452. Each subplot presents results for five replications indicated by different line colors.

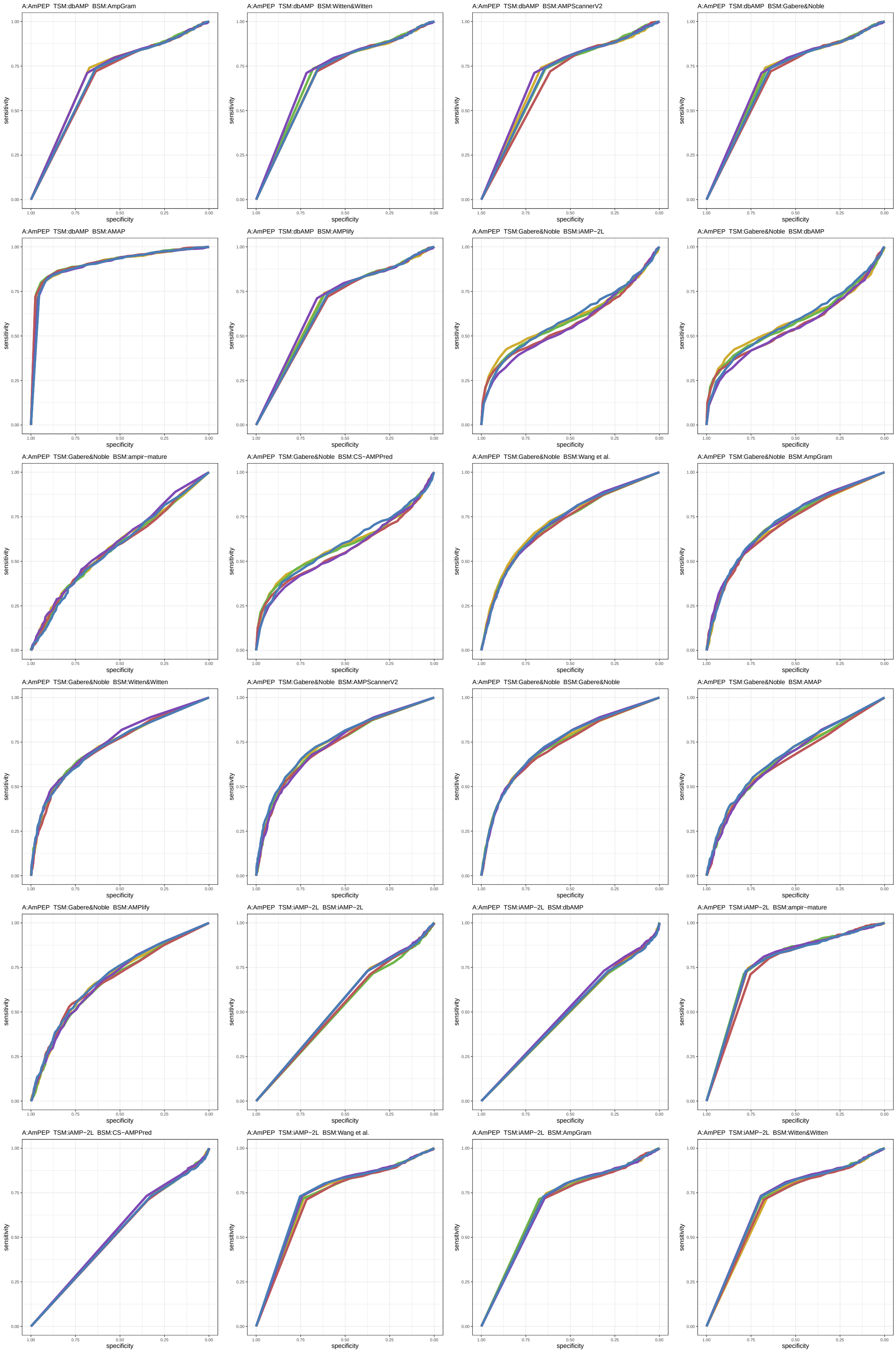

Figure S17: ROC curves 193-216 of 1452. Each subplot presents results for five replications indicated by different line colors.

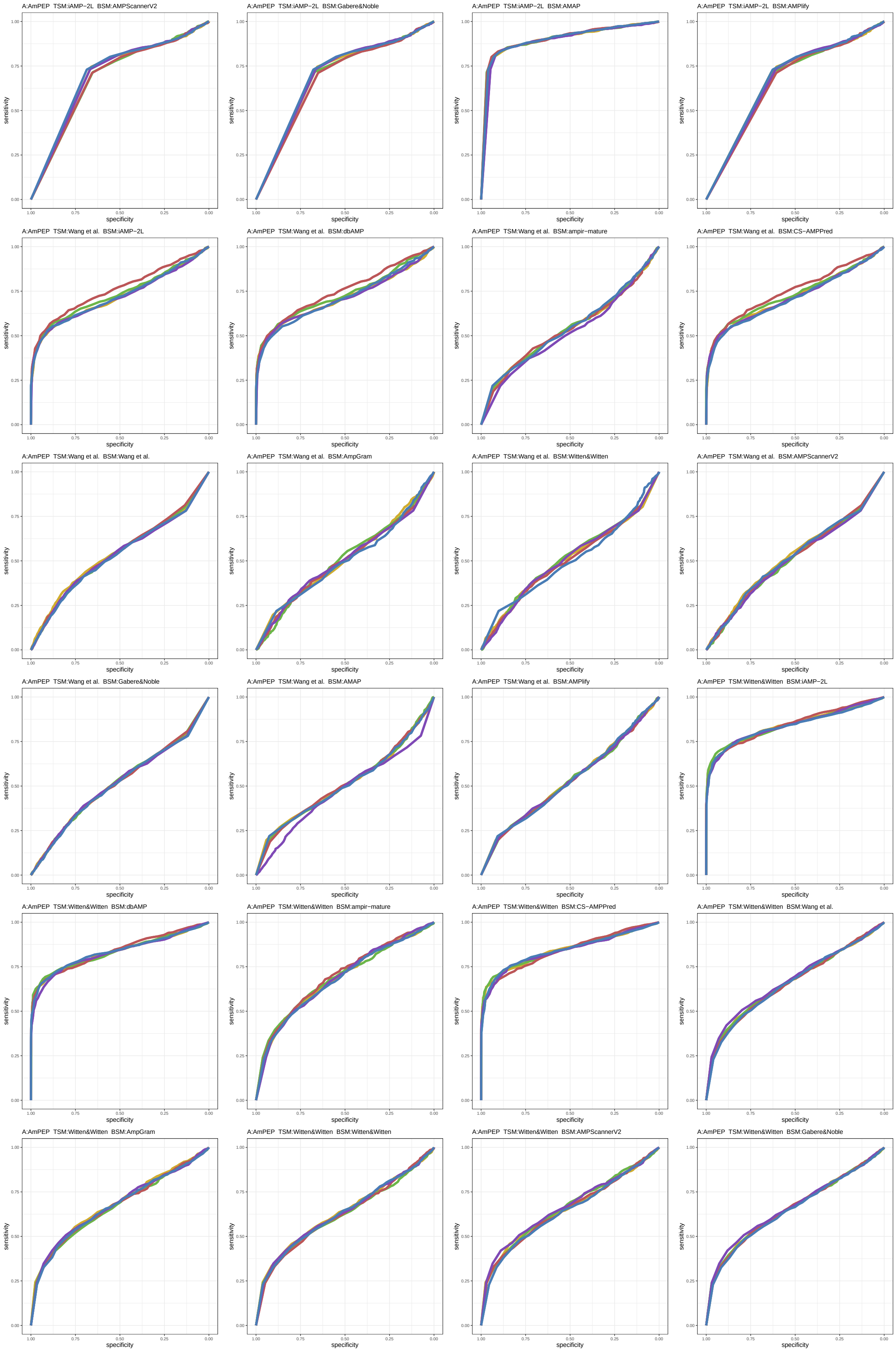

Figure S18: ROC curves 217-240 of 1452. Each subplot presents results for five replications indicated by different line colors.

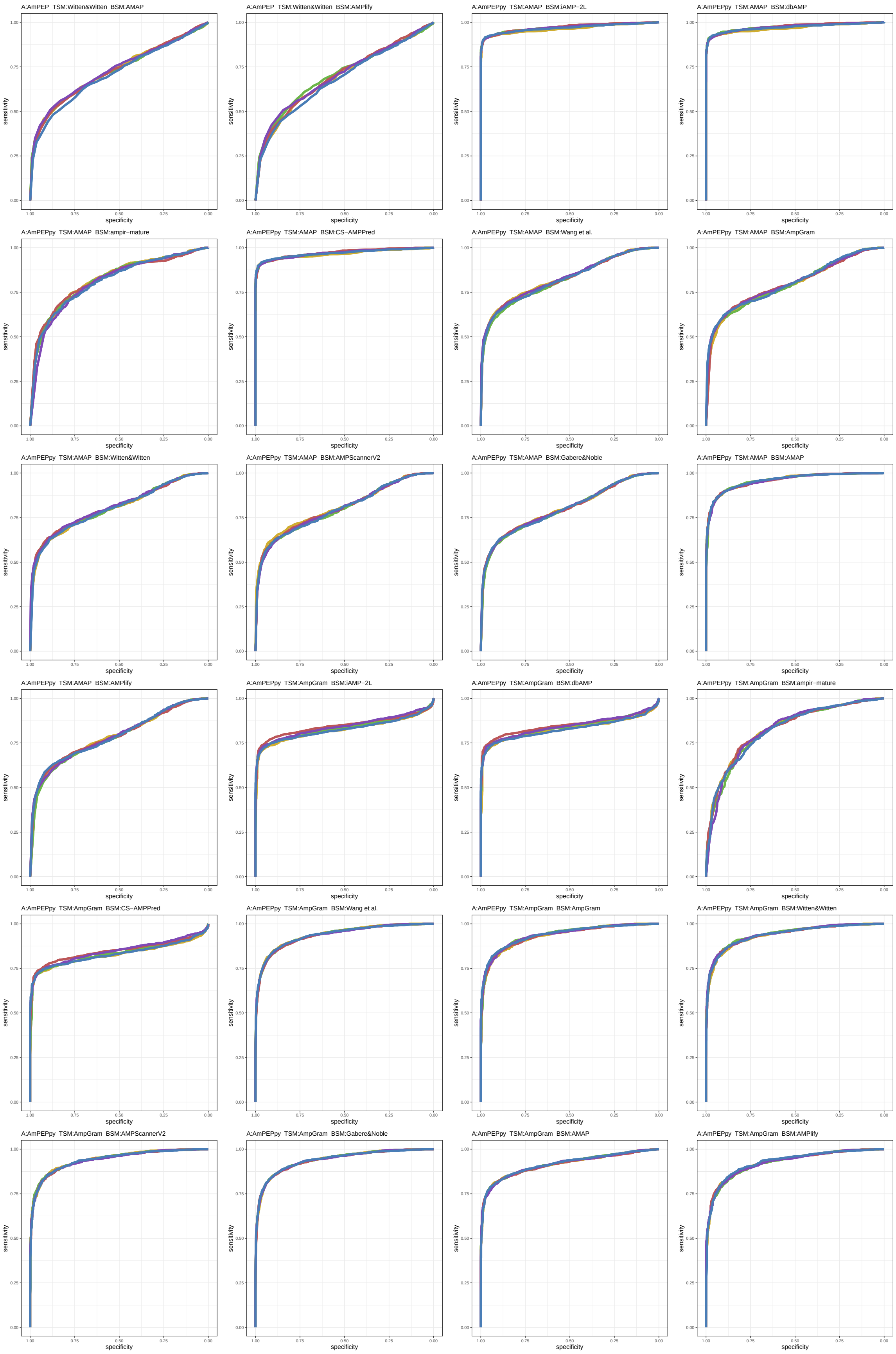

Figure S19: ROC curves 241-264 of 1452. Each subplot presents results for five replications indicated by different line colors.

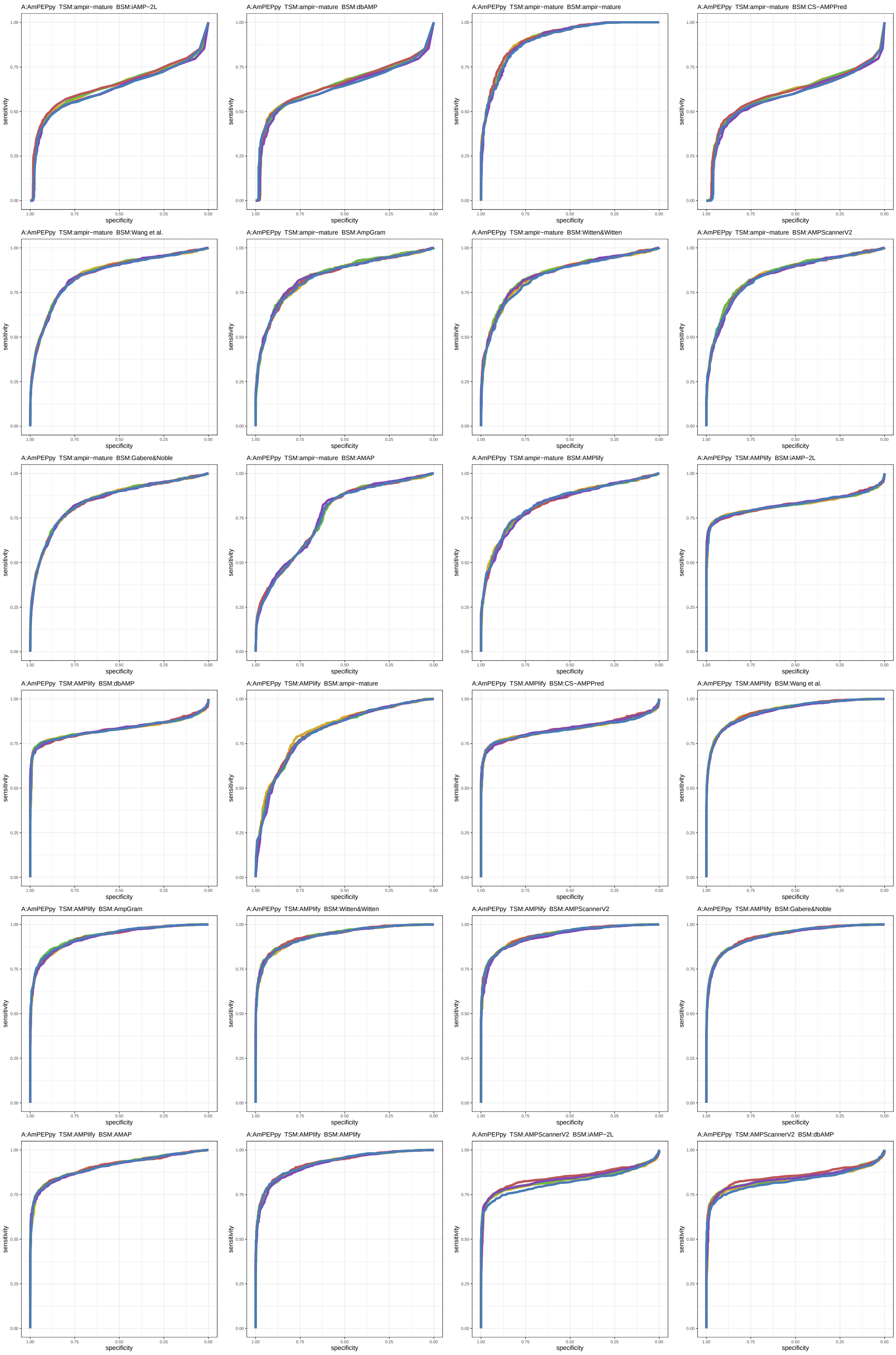

Figure S20: ROC curves 265-288 of 1452. Each subplot presents results for five replications indicated by different line colors.

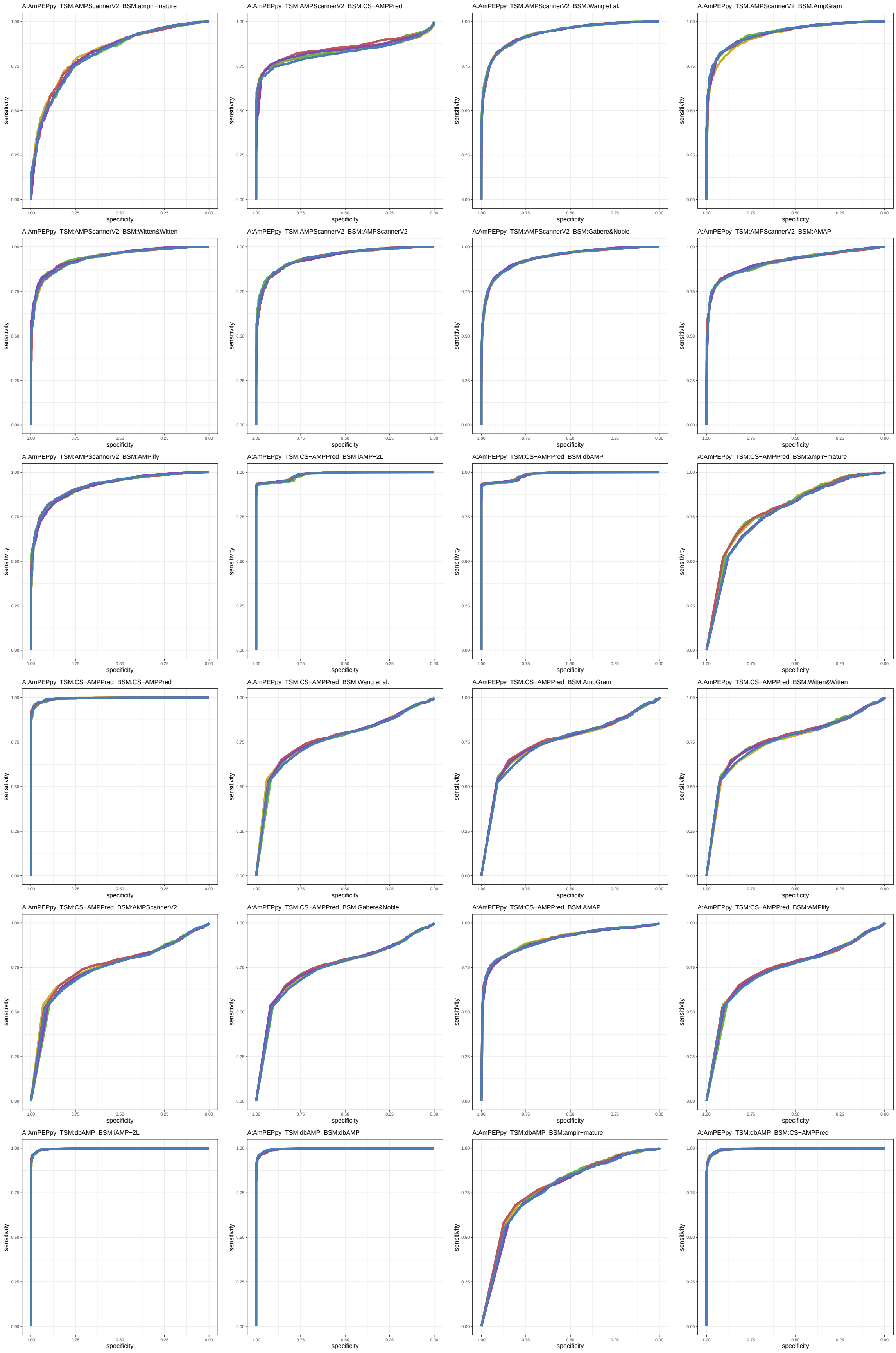

Figure S21: ROC curves 289-312 of 1452. Each subplot presents results for five replications indicated by different line colors.

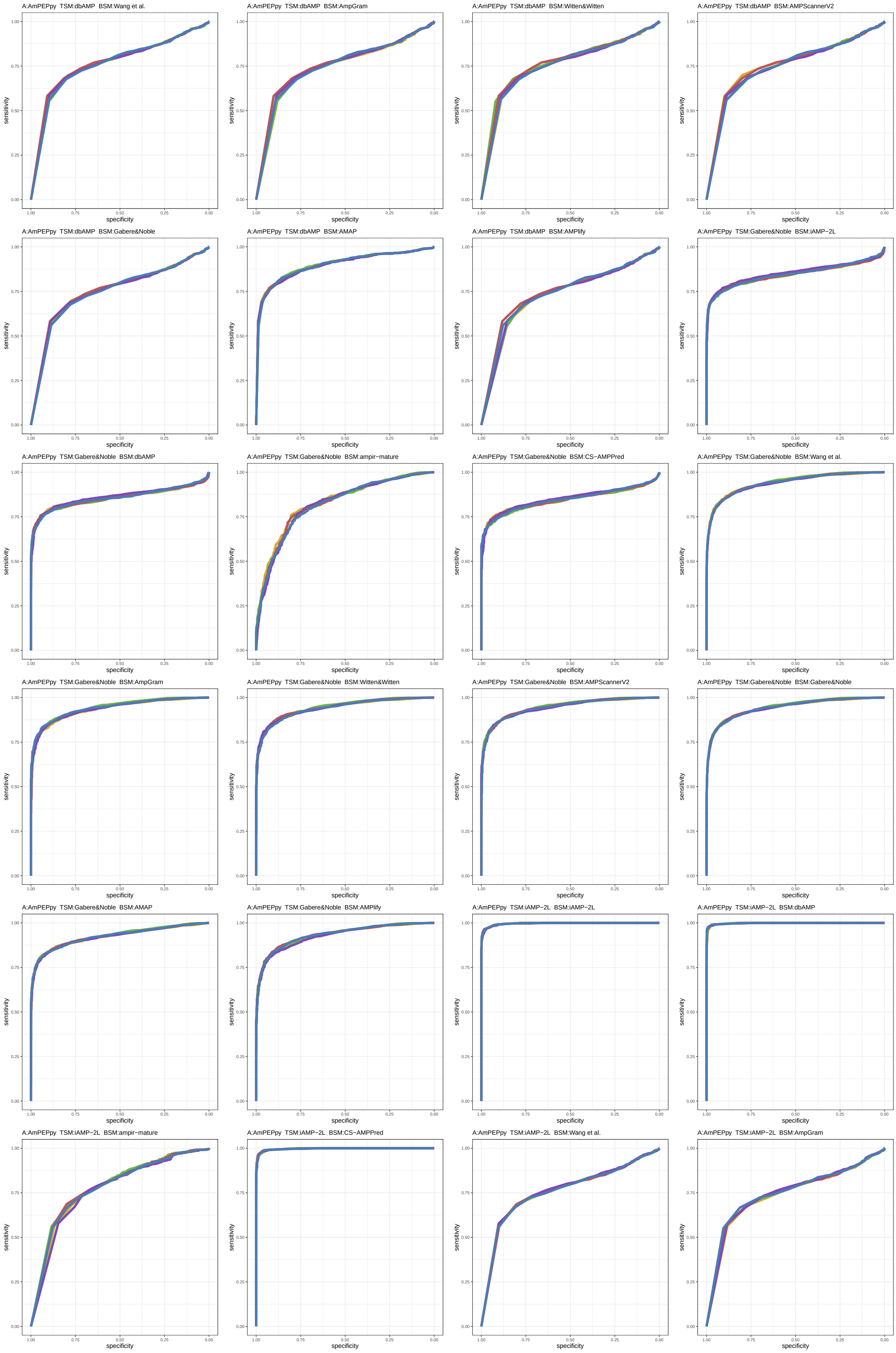

Figure S22: ROC curves 313-336 of 1452. Each subplot presents results for five replications indicated by different line colors.

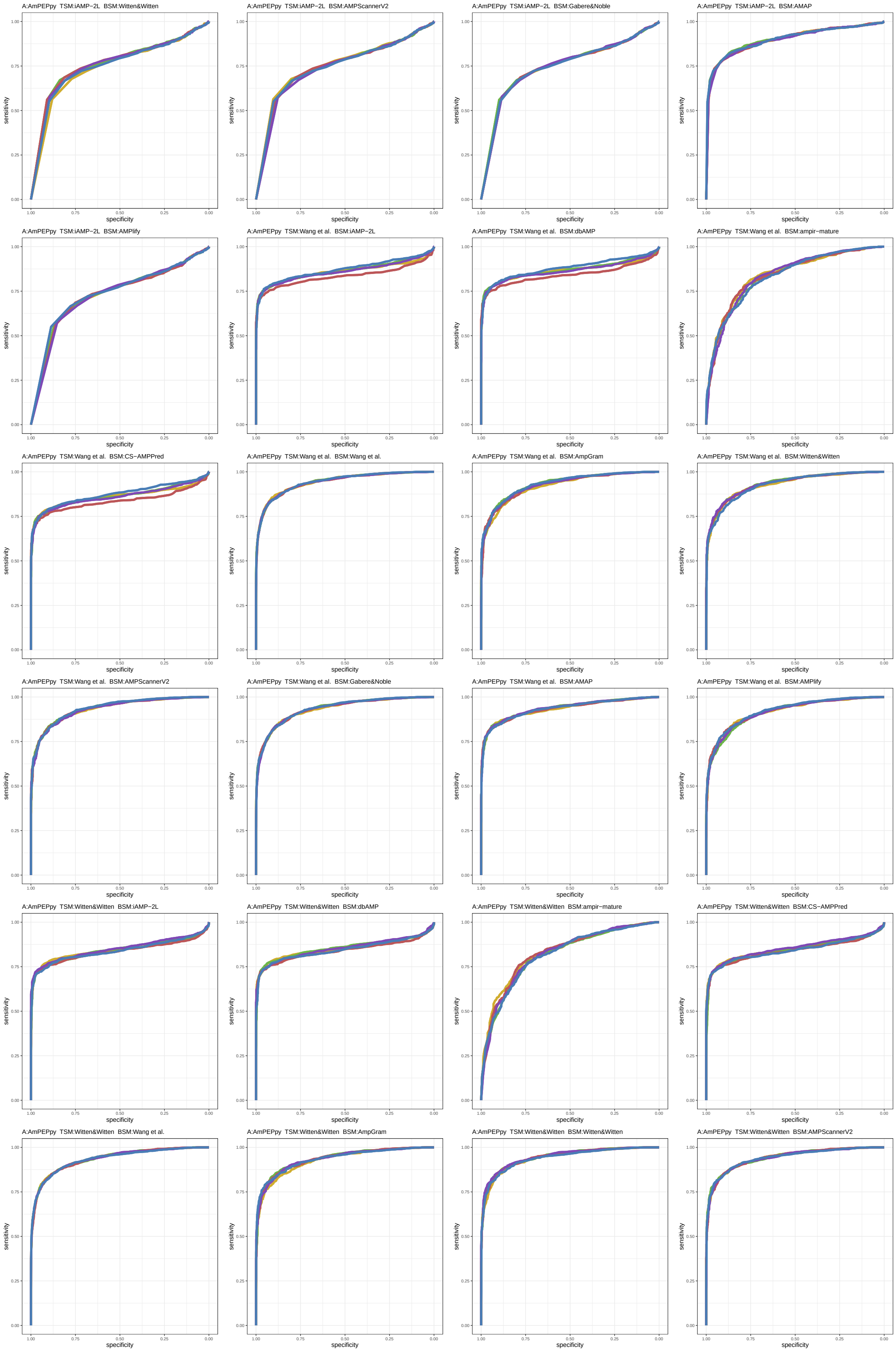

Figure S23: ROC curves 337-360 of 1452. Each subplot presents results for five replications indicated by different line colors.

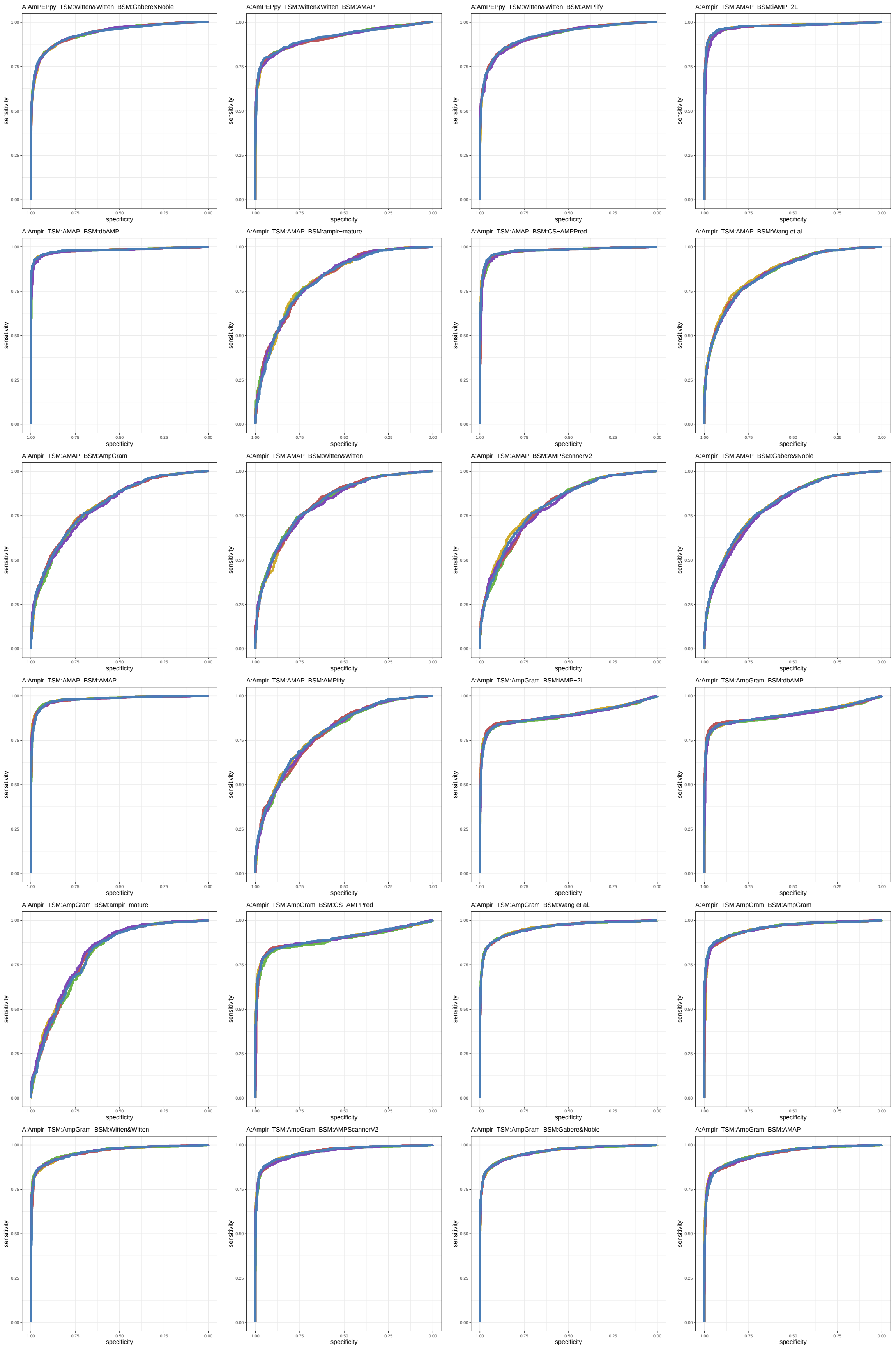

Figure S24: ROC curves 361-384 of 1452. Each subplot presents results for five replications indicated by different line colors.

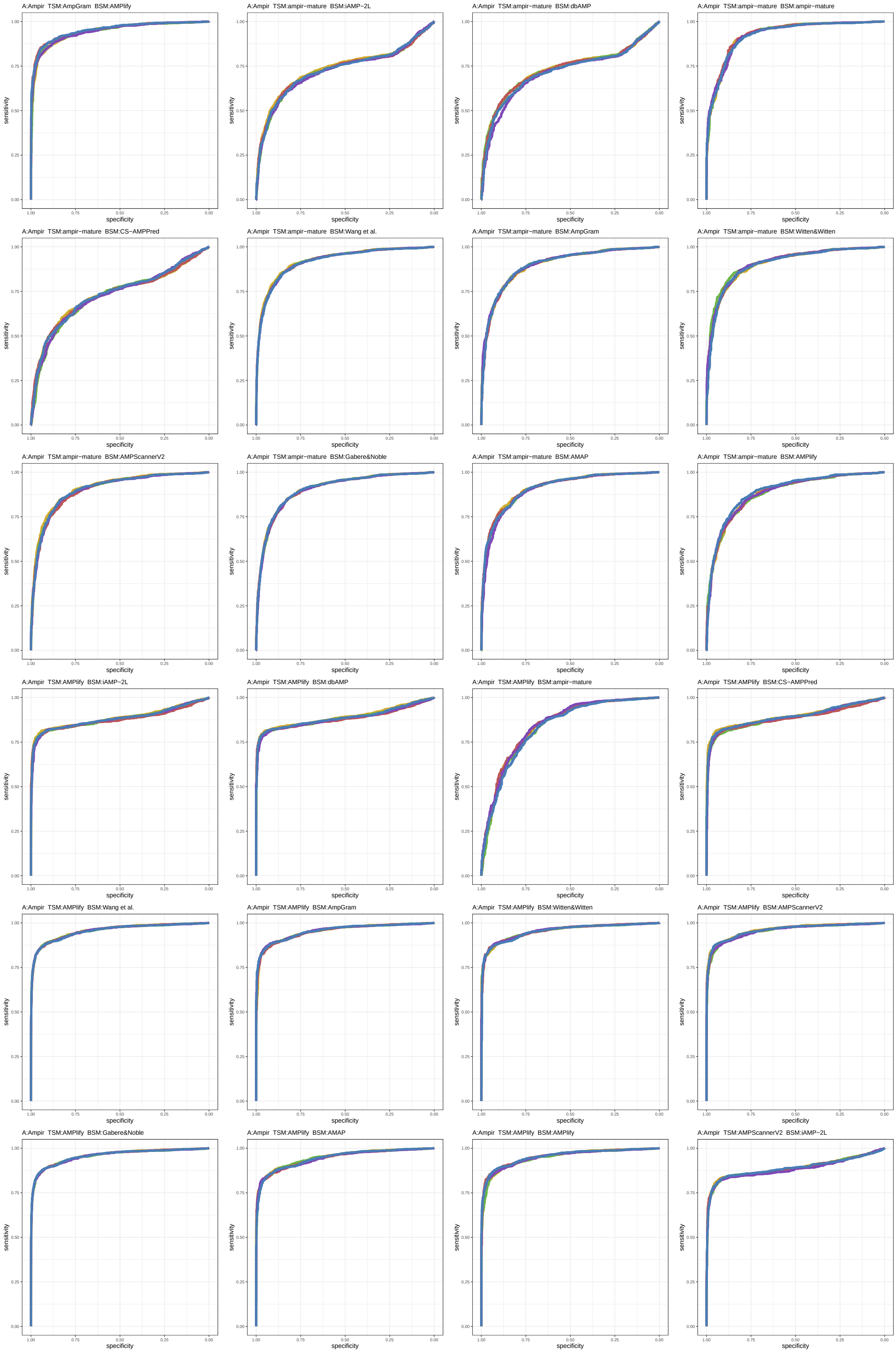

Figure S25: ROC curves 385-408 of 1452. Each subplot presents results for five replications indicated by different line colors.

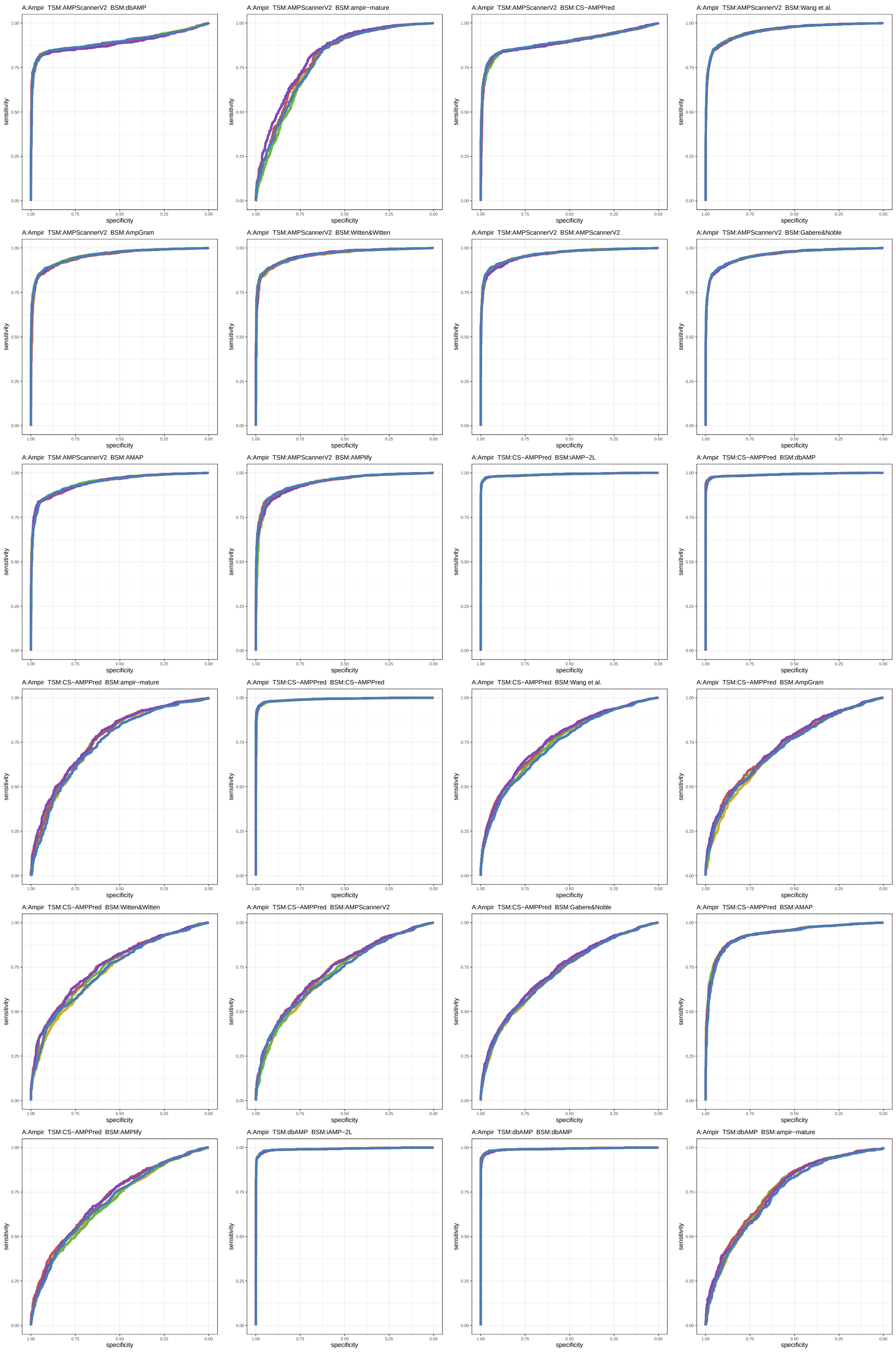

Figure S26: ROC curves 409-432 of 1452. Each subplot presents results for five replications indicated by different line colors.

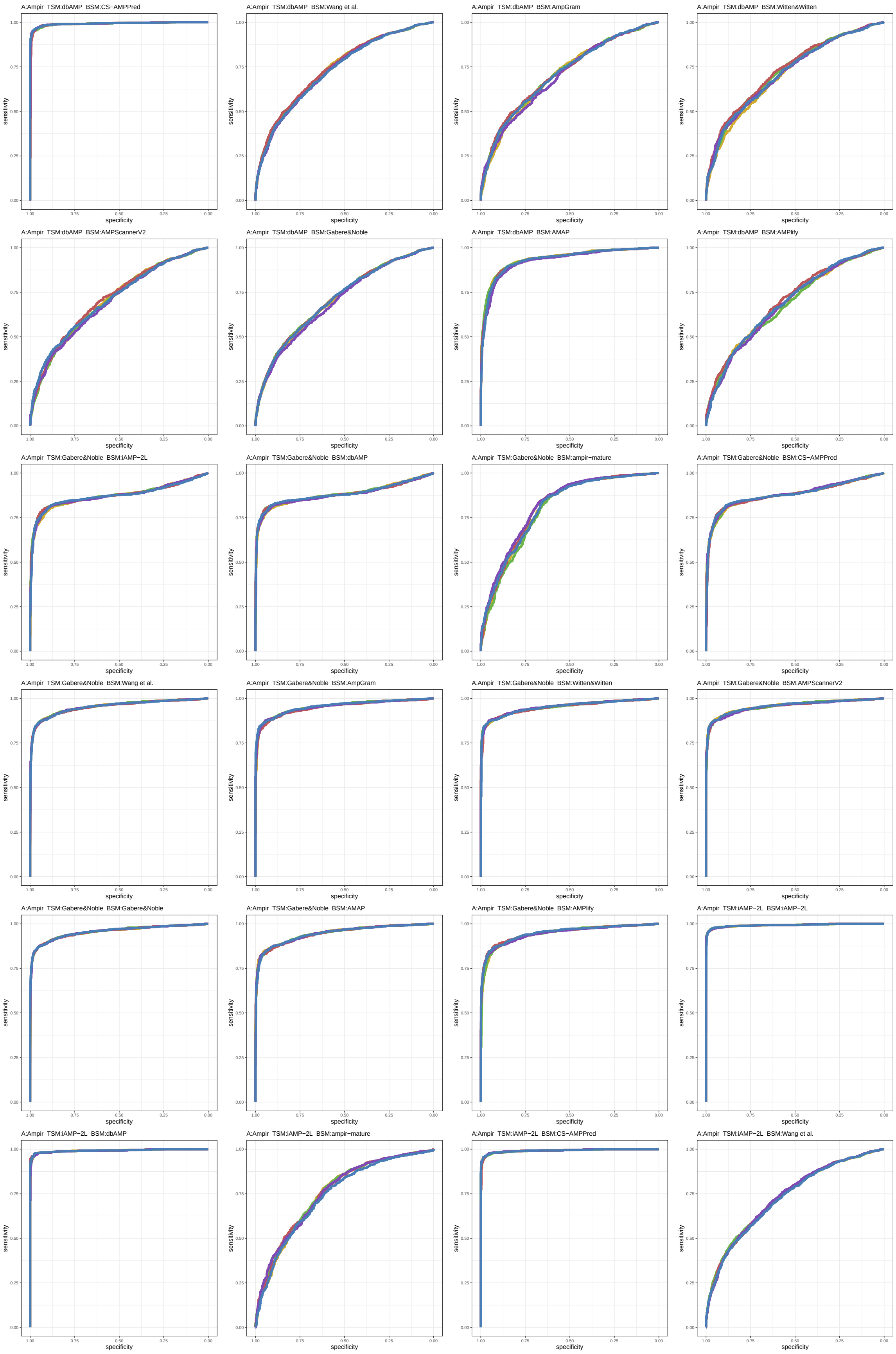

Figure S27: ROC curves 433-456 of 1452. Each subplot presents results for five replications indicated by different line colors.

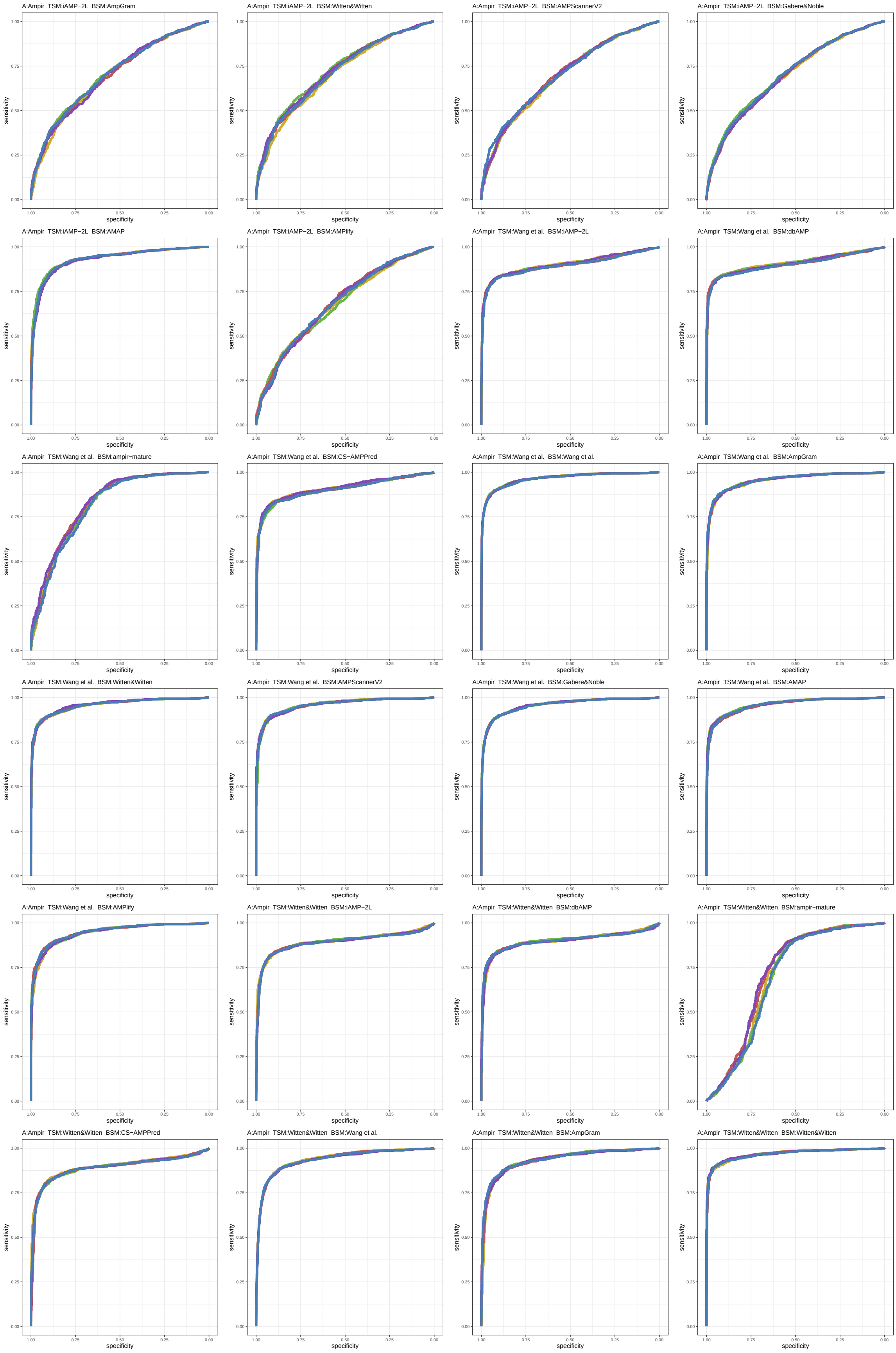

Figure S28: ROC curves 457-480 of 1452. Each subplot presents results for five replications indicated by different line colors.

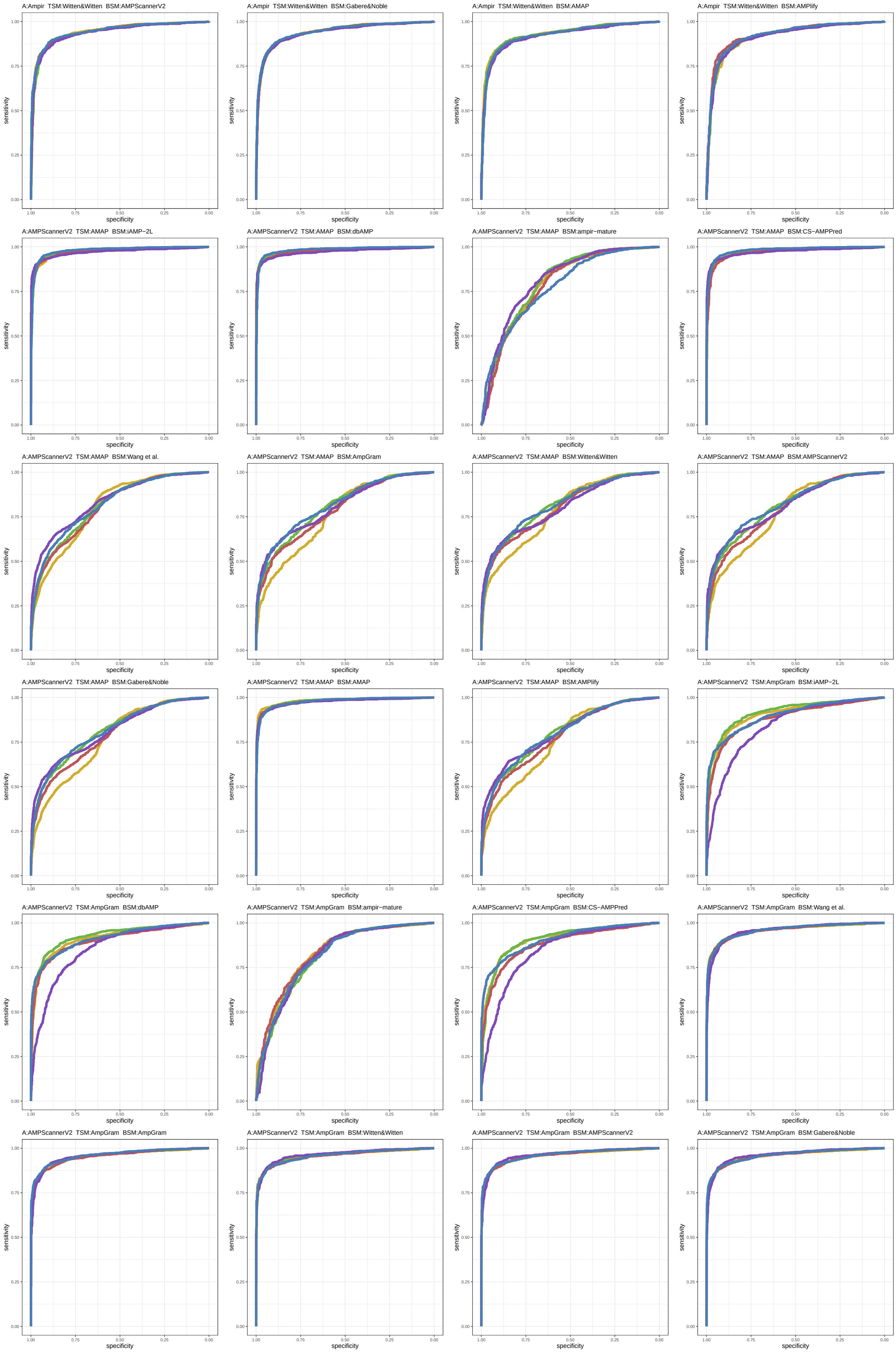

Figure S29: ROC curves 481-504 of 1452. Each subplot presents results for five replications indicated by different line colors.

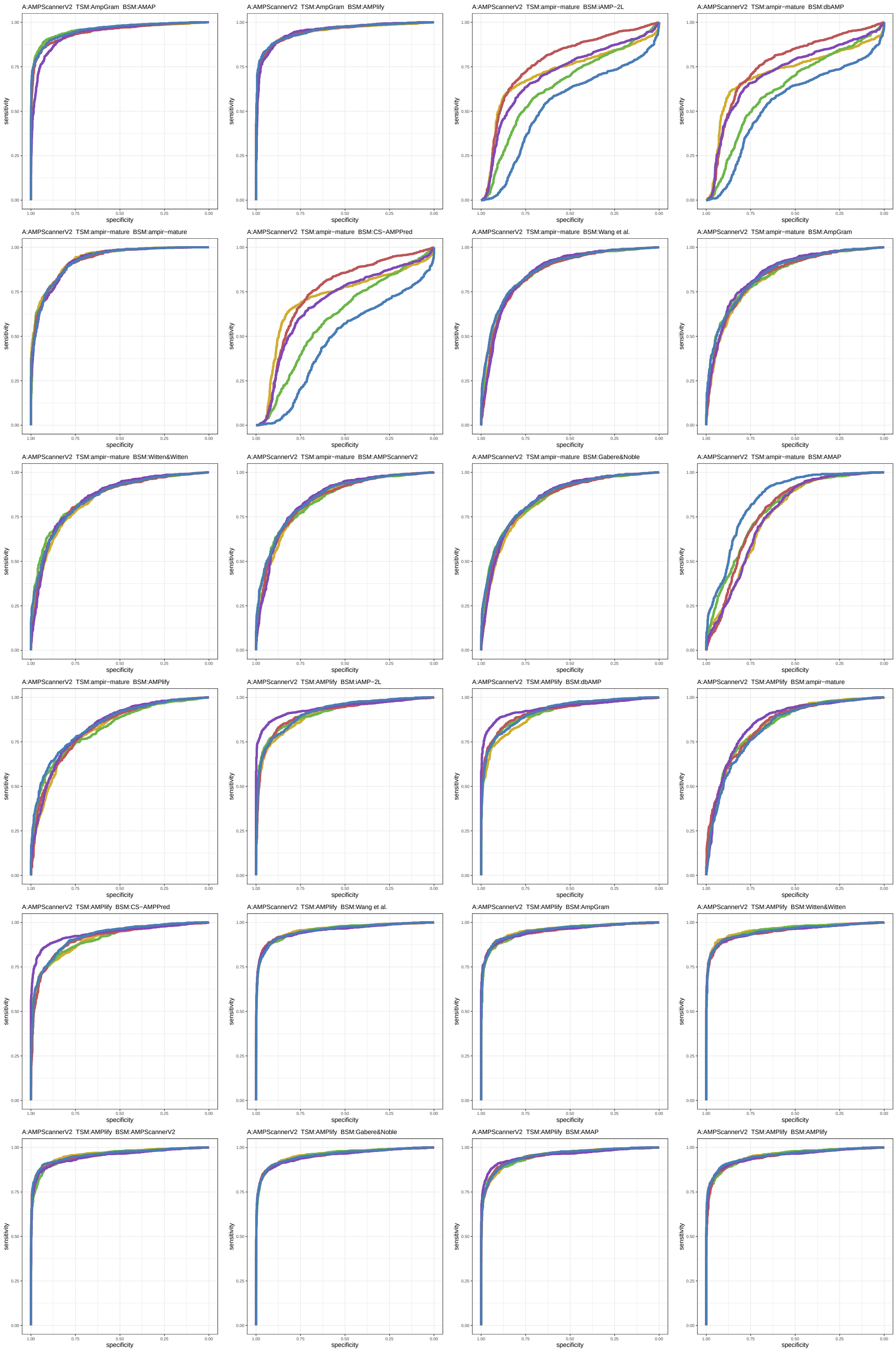

Figure S30: ROC curves 505-528 of 1452. Each subplot presents results for five replications indicated by different line colors.

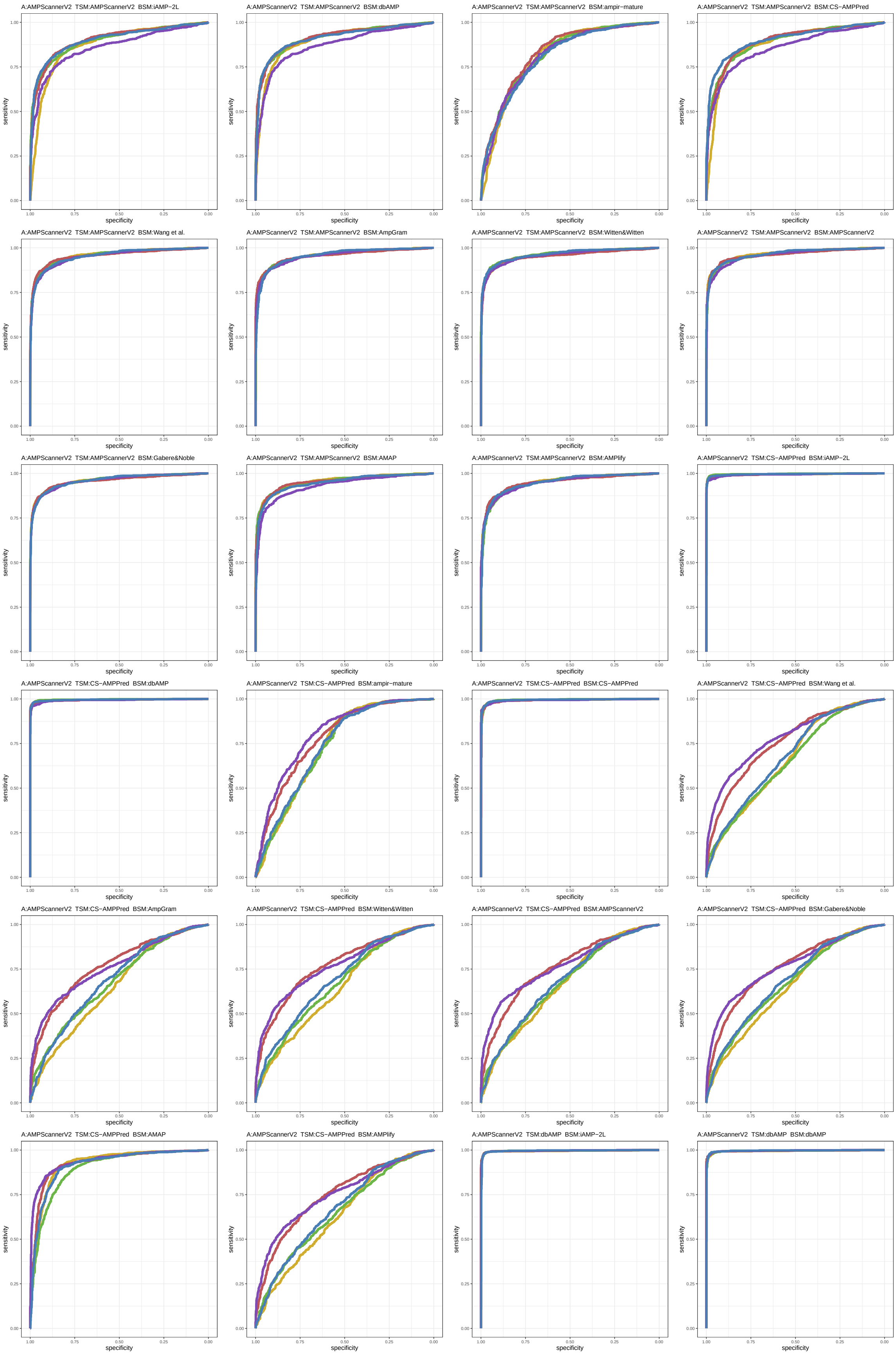

Figure S31: ROC curves 529-552 of 1452. Each subplot presents results for five replications indicated by different line colors.

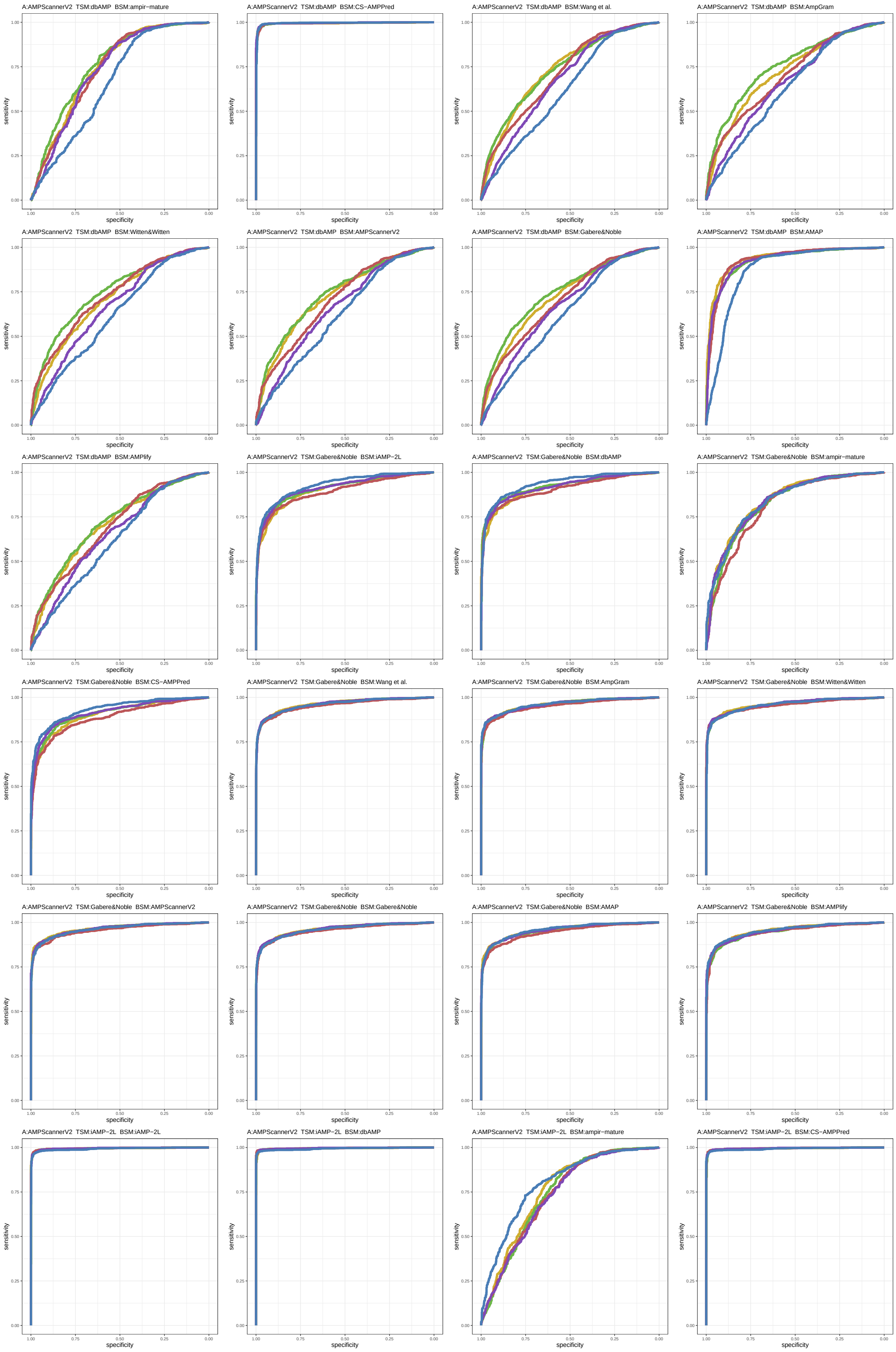

Figure S32: ROC curves 553-576 of 1452. Each subplot presents results for five replications indicated by different line colors.

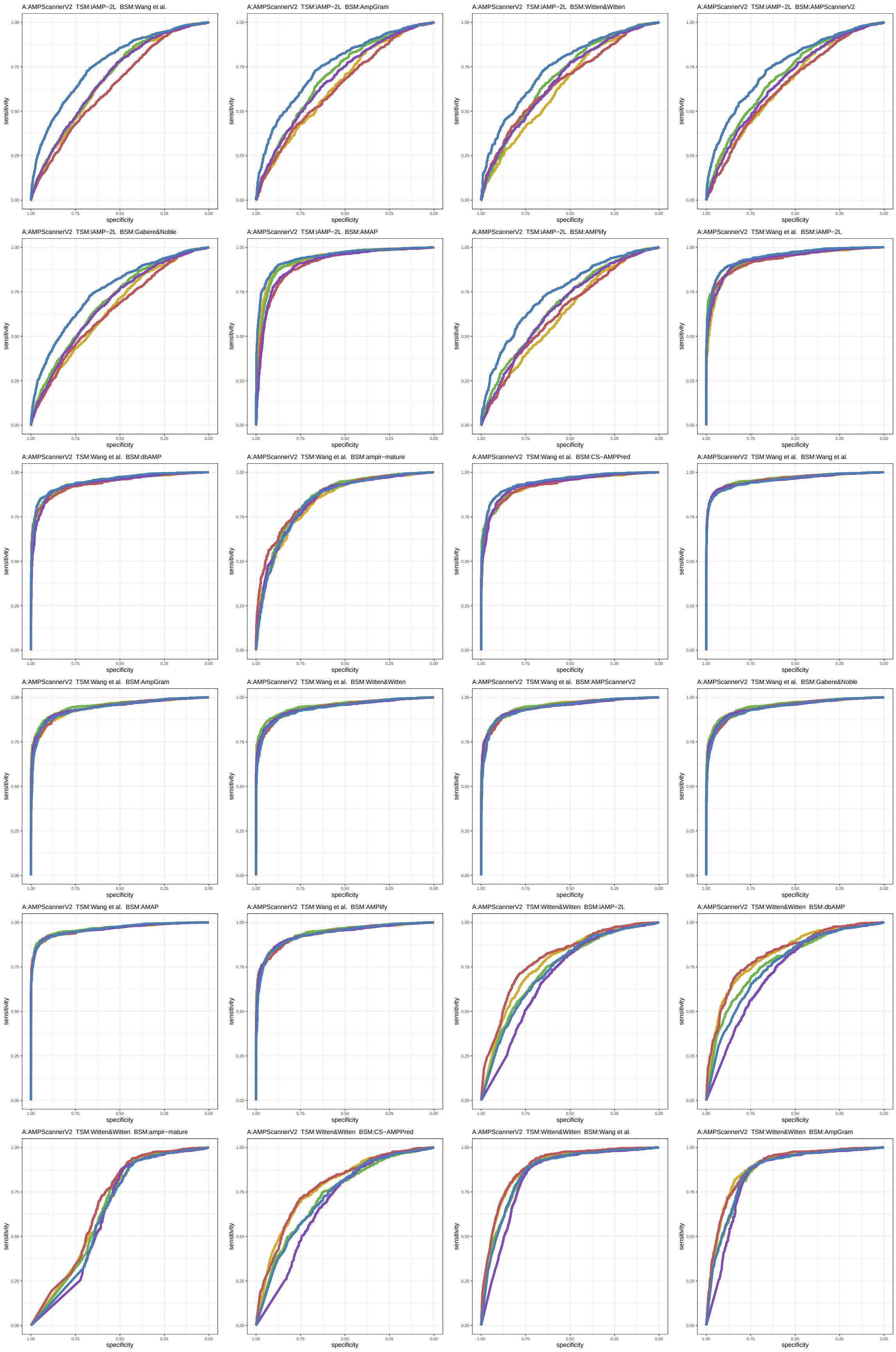

Figure S33: ROC curves 577-600 of 1452. Each subplot presents results for five replications indicated by different line colors.

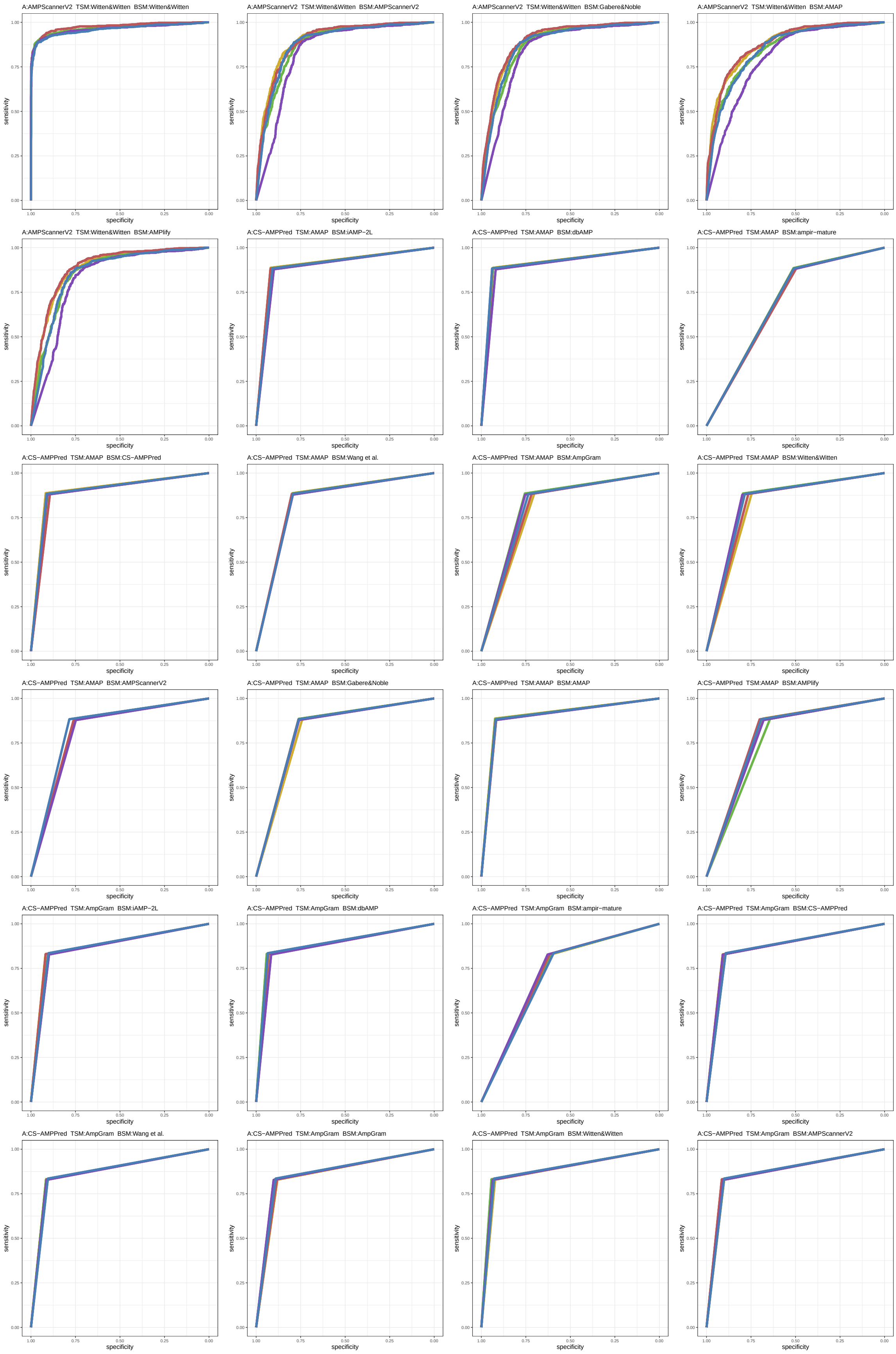

Figure S34: ROC curves 601-624 of 1452. Each subplot presents results for five replications indicated by different line colors.

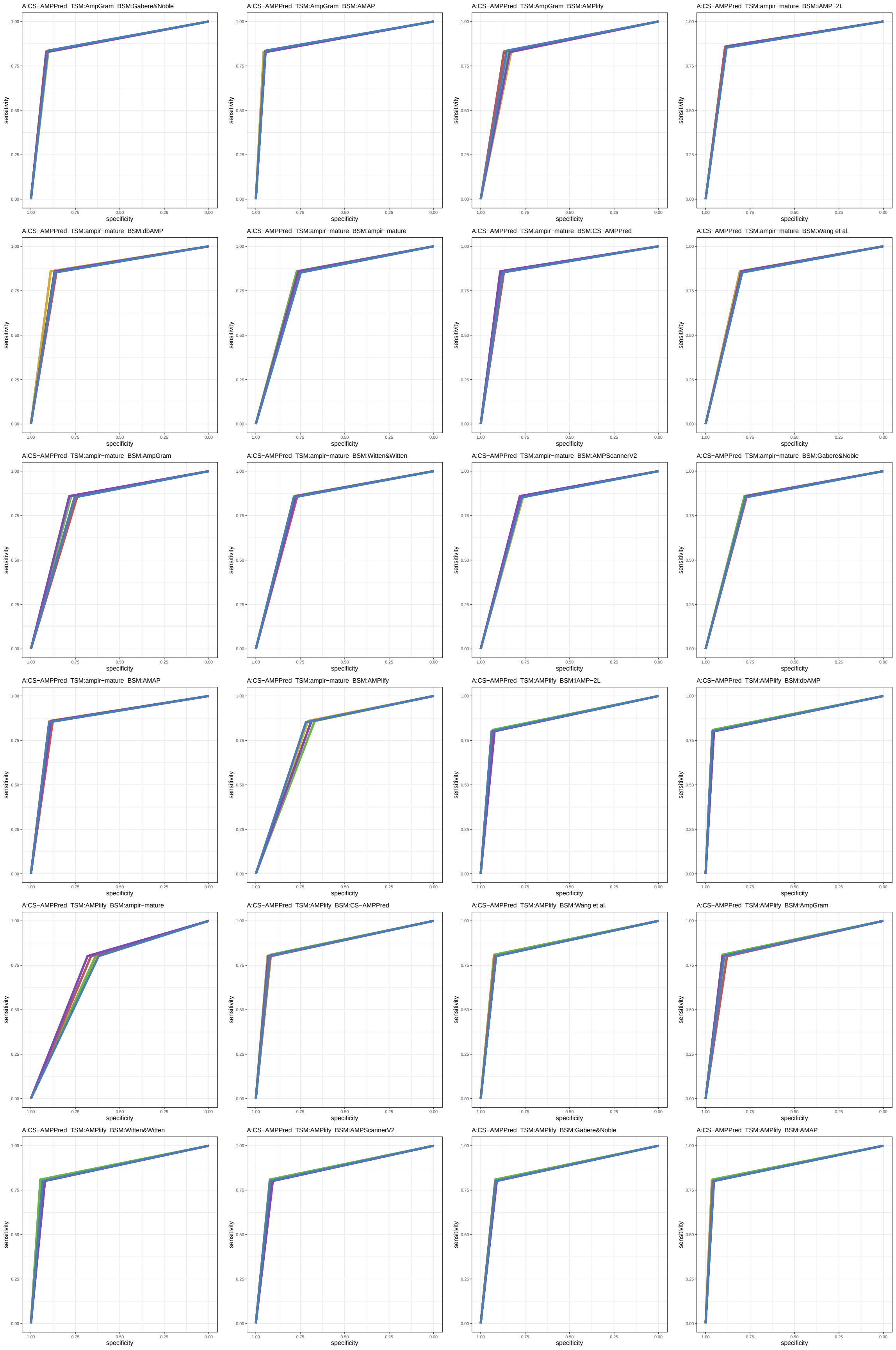

Figure S35: ROC curves 625-648 of 1452. Each subplot presents results for five replications indicated by different line colors.

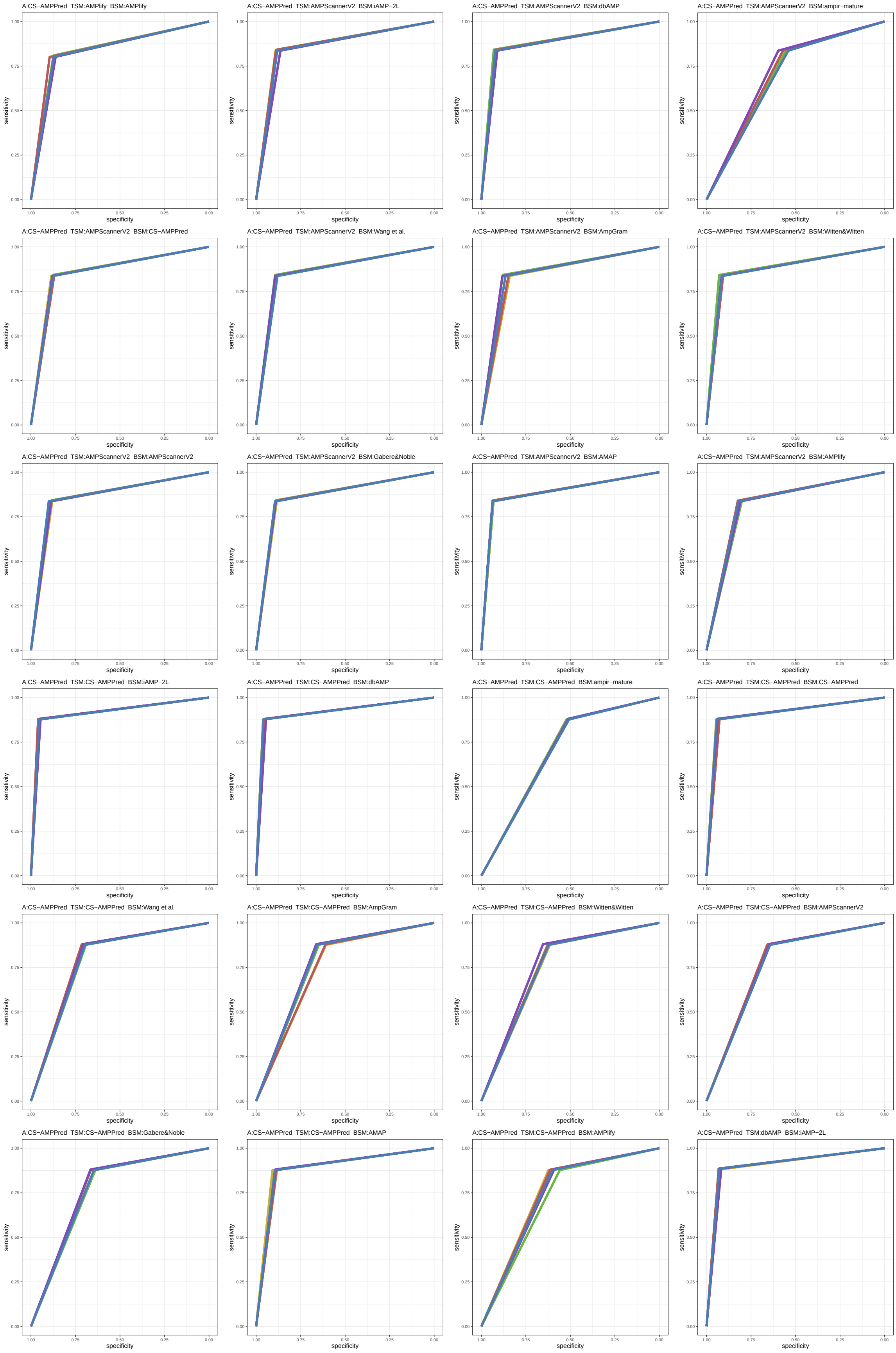

Figure S36: ROC curves 649-672 of 1452. Each subplot presents results for five replications indicated by different line colors.

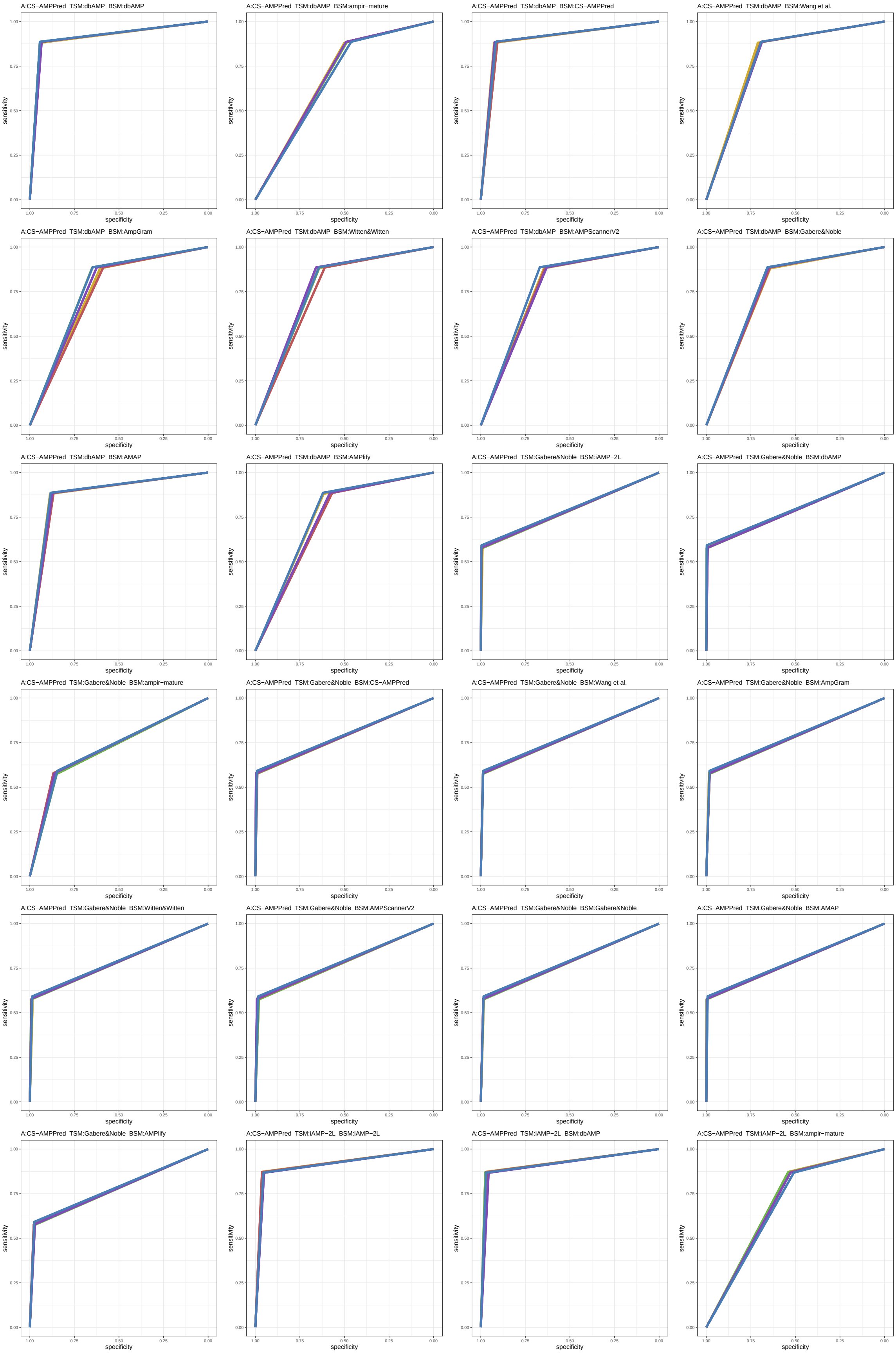

Figure S37: ROC curves 673-696 of 1452. Each subplot presents results for five replications indicated by different line colors.

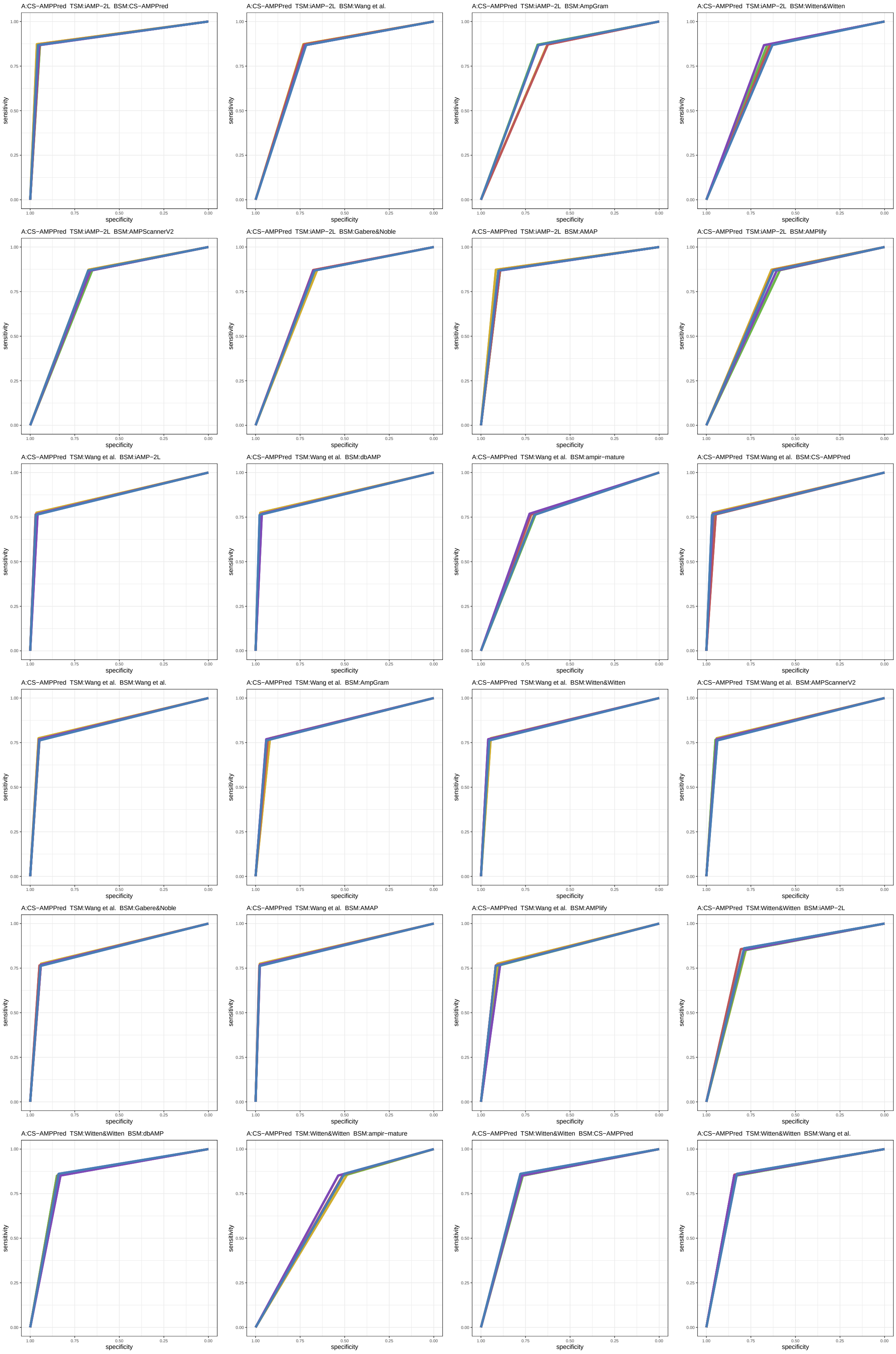

Figure S38: ROC curves 697-720 of 1452. Each subplot presents results for five replications indicated by different line colors.

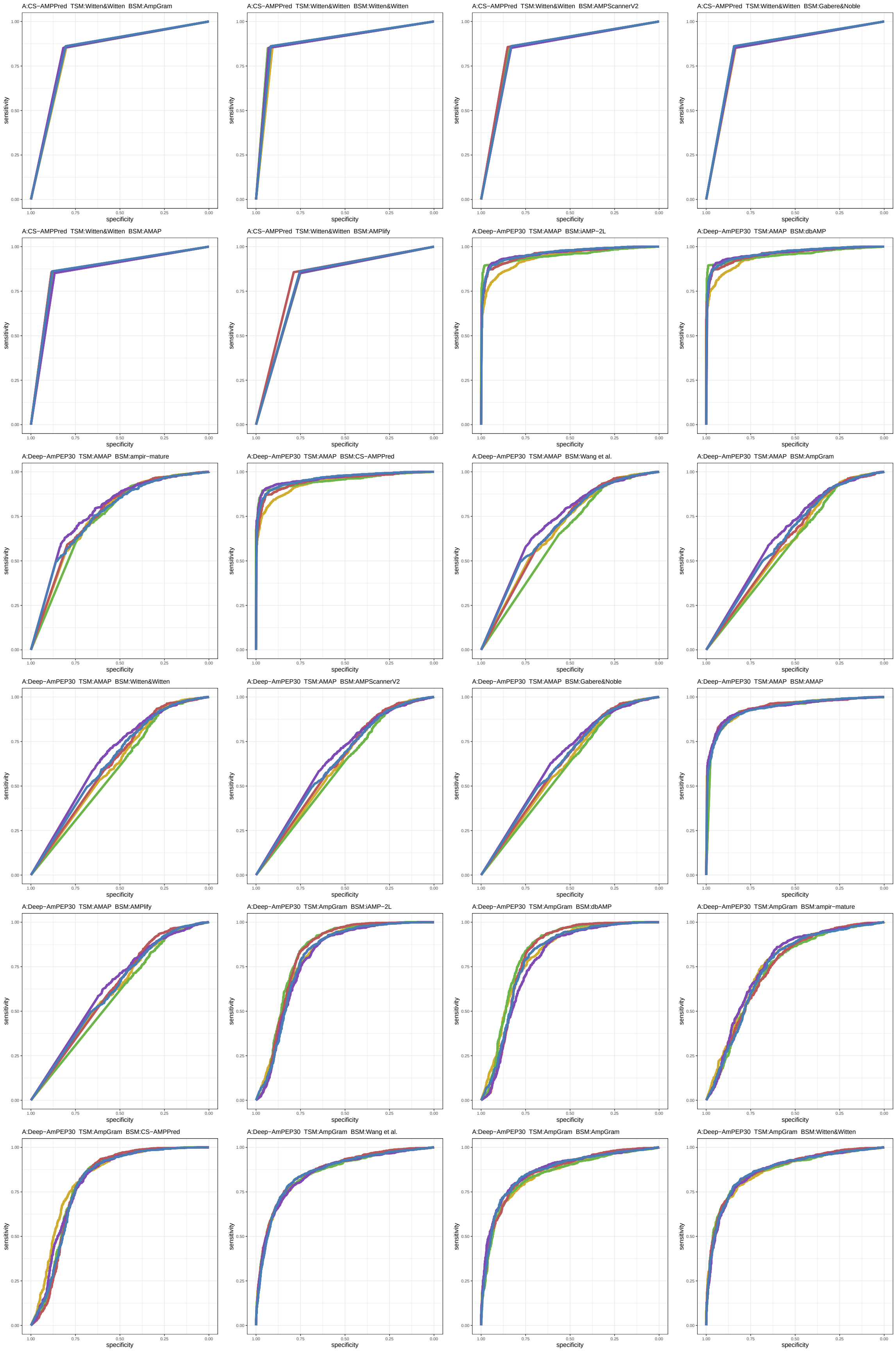

Figure S39: ROC curves 721-744 of 1452. Each subplot presents results for five replications indicated by different line colors.

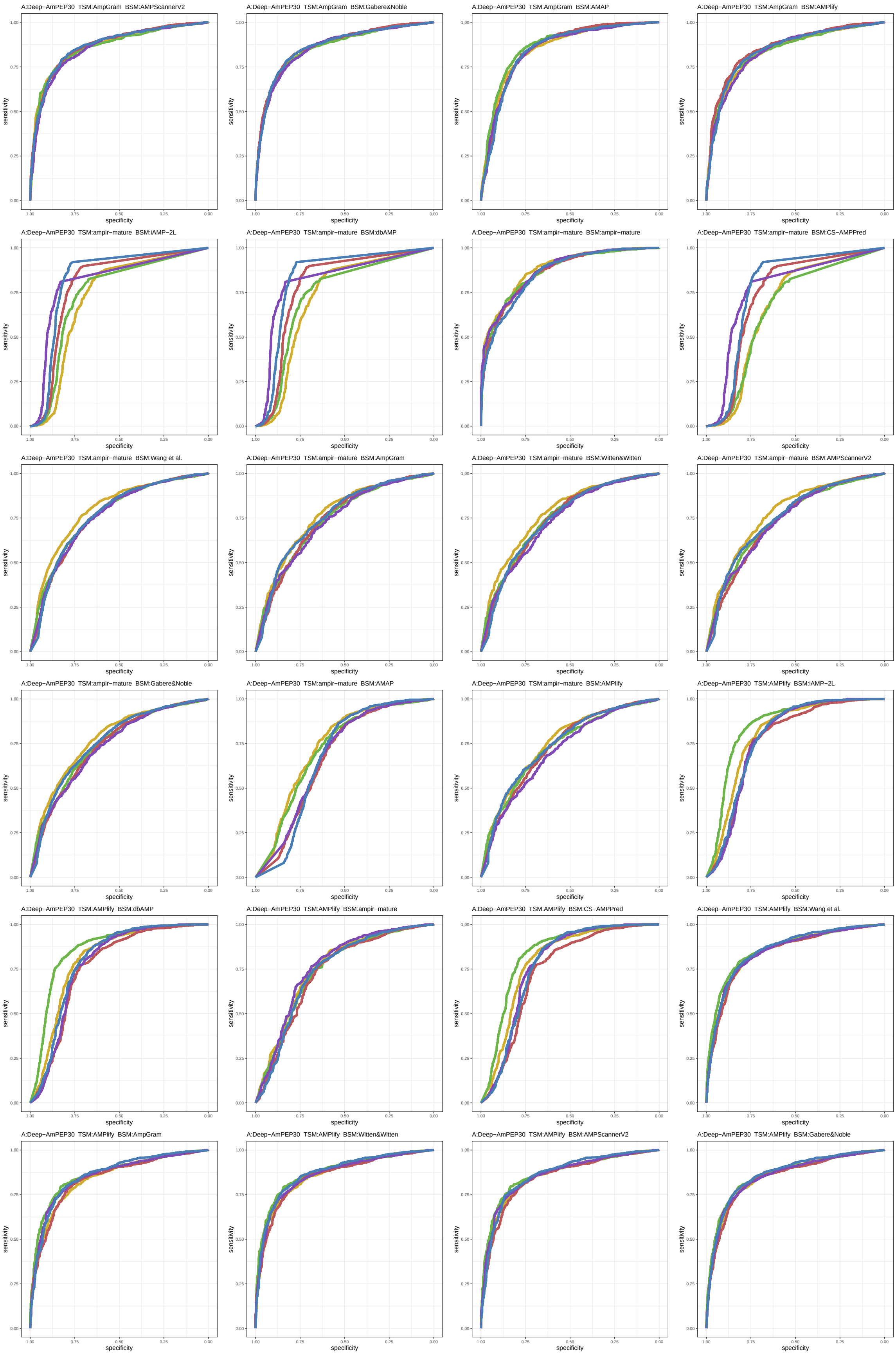

Figure S40: ROC curves 745-768 of 1452. Each subplot presents results for five replications indicated by different line colors.

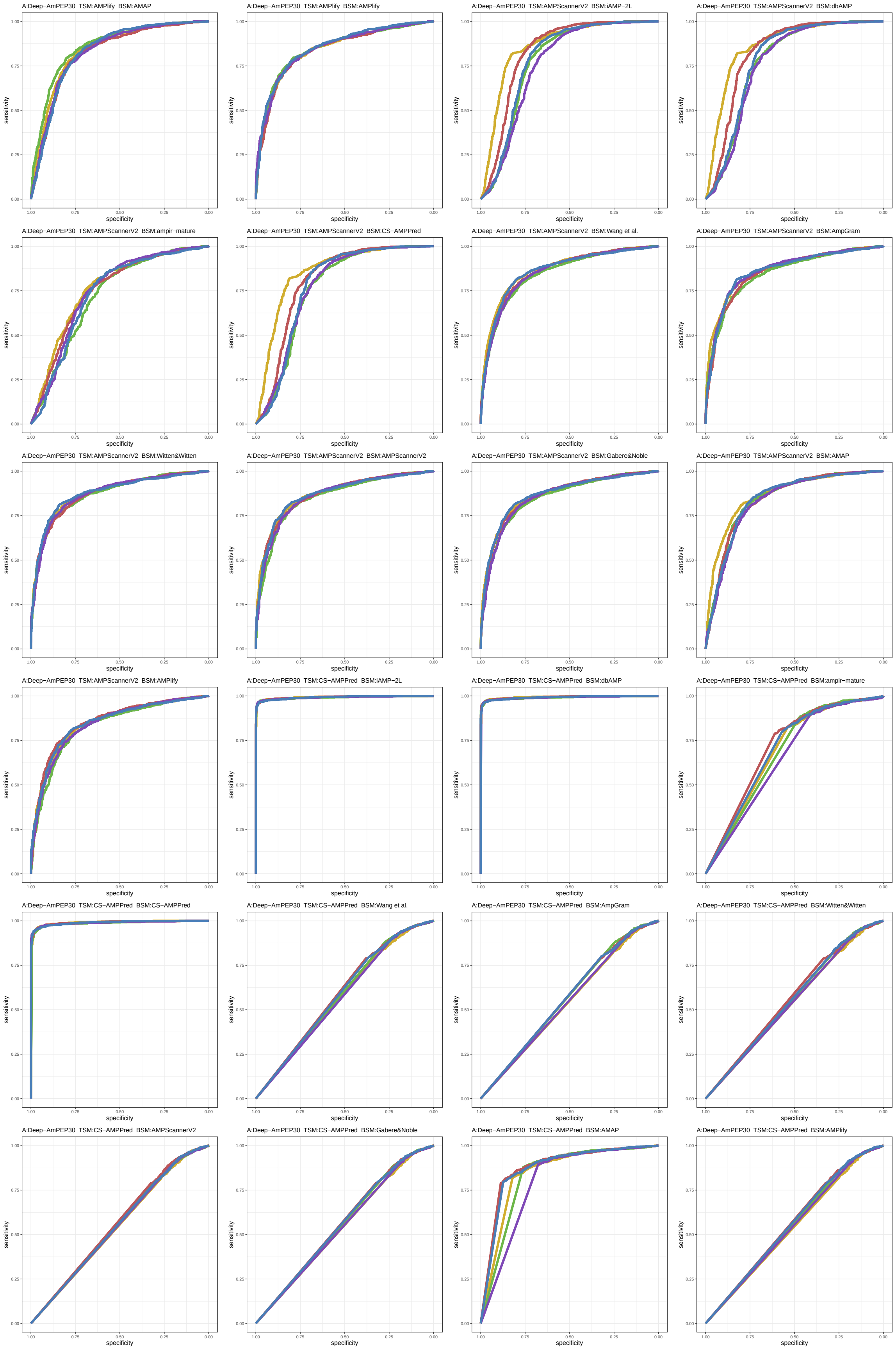

Figure S41: ROC curves 769-792 of 1452. Each subplot presents results for five replications indicated by different line colors.

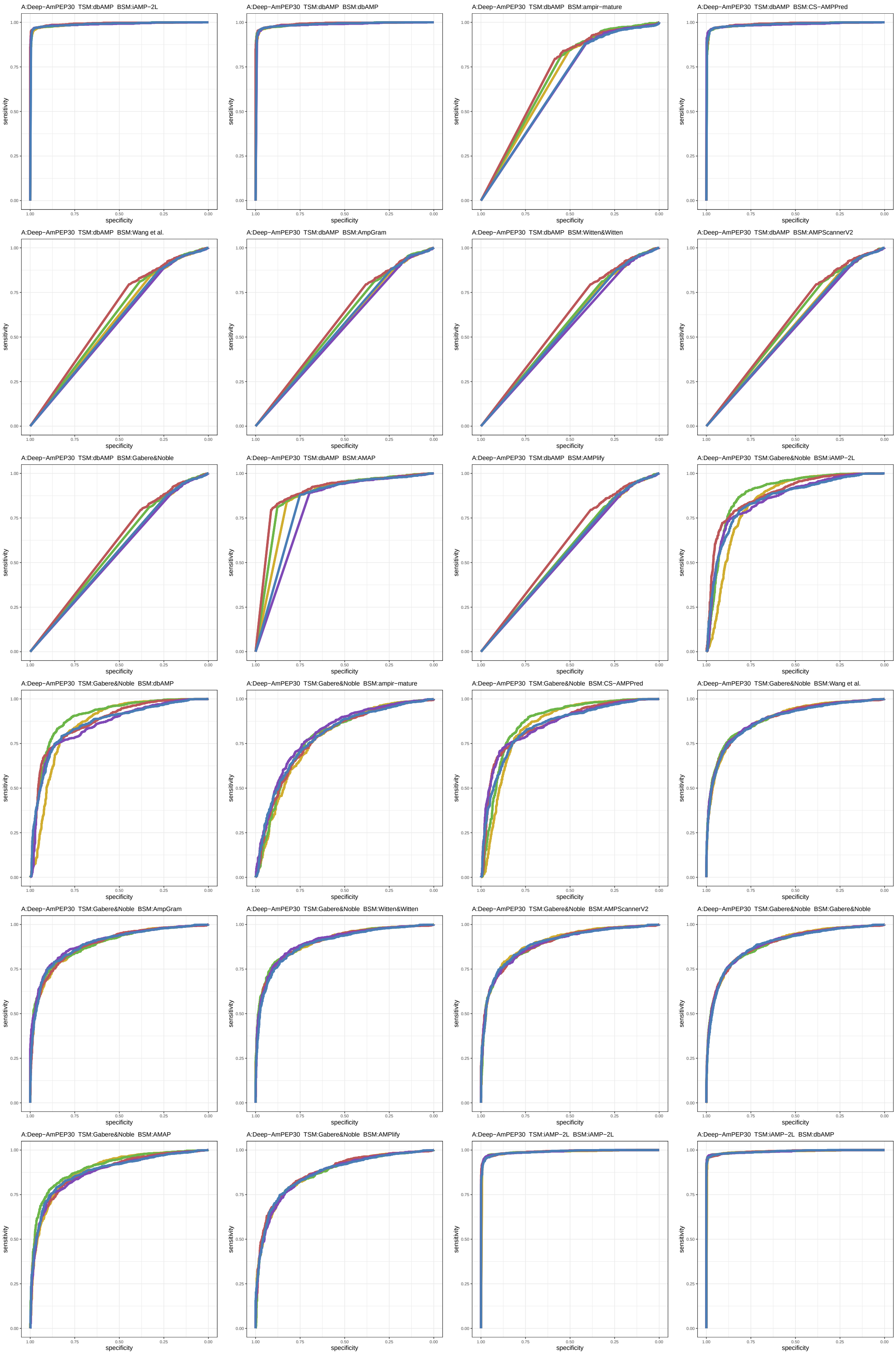

Figure S42: ROC curves 793-816 of 1452. Each subplot presents results for five replications indicated by different line colors.

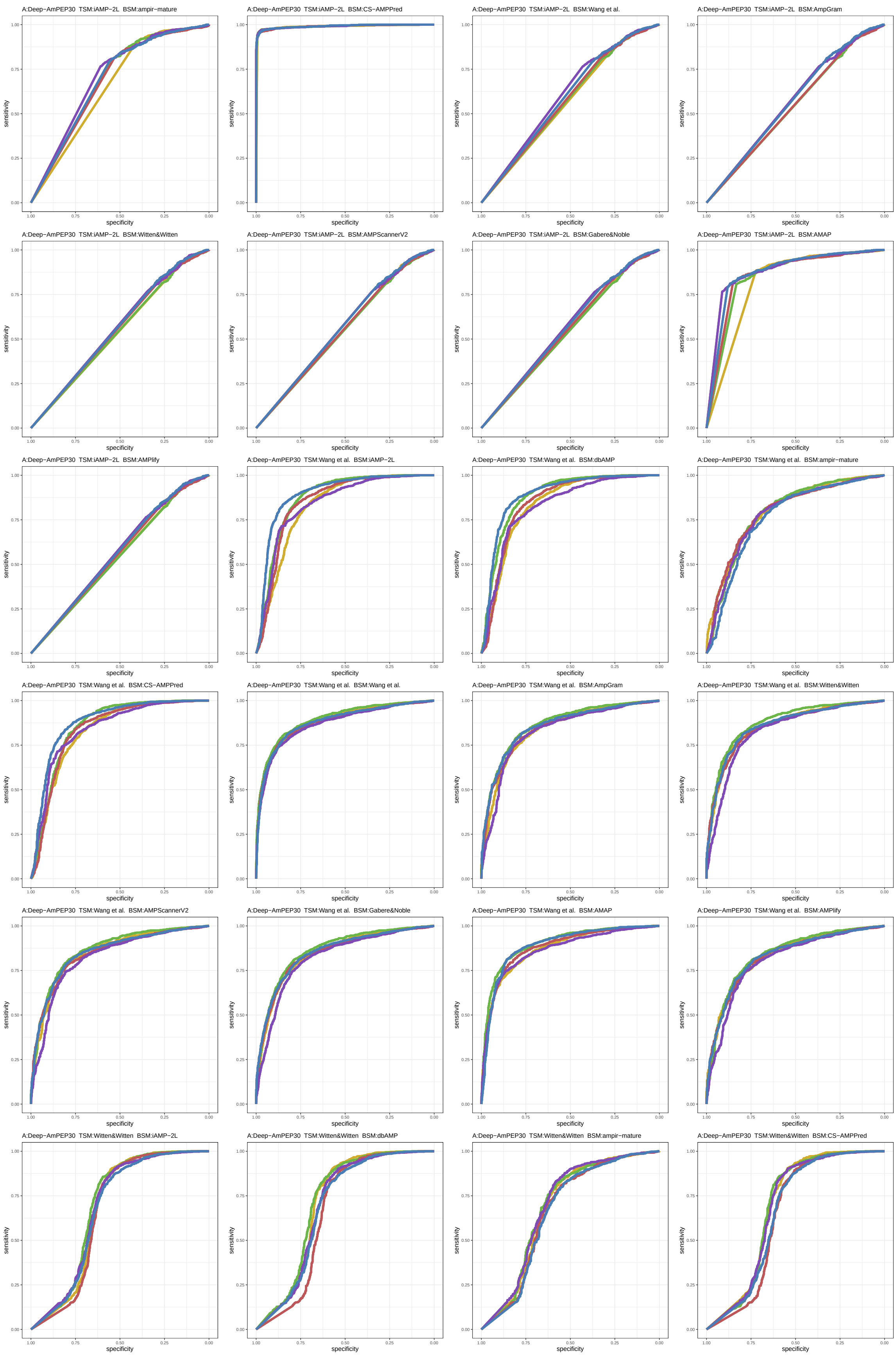

Figure S43: ROC curves 817-840 of 1452. Each subplot presents results for five replications indicated by different line colors.

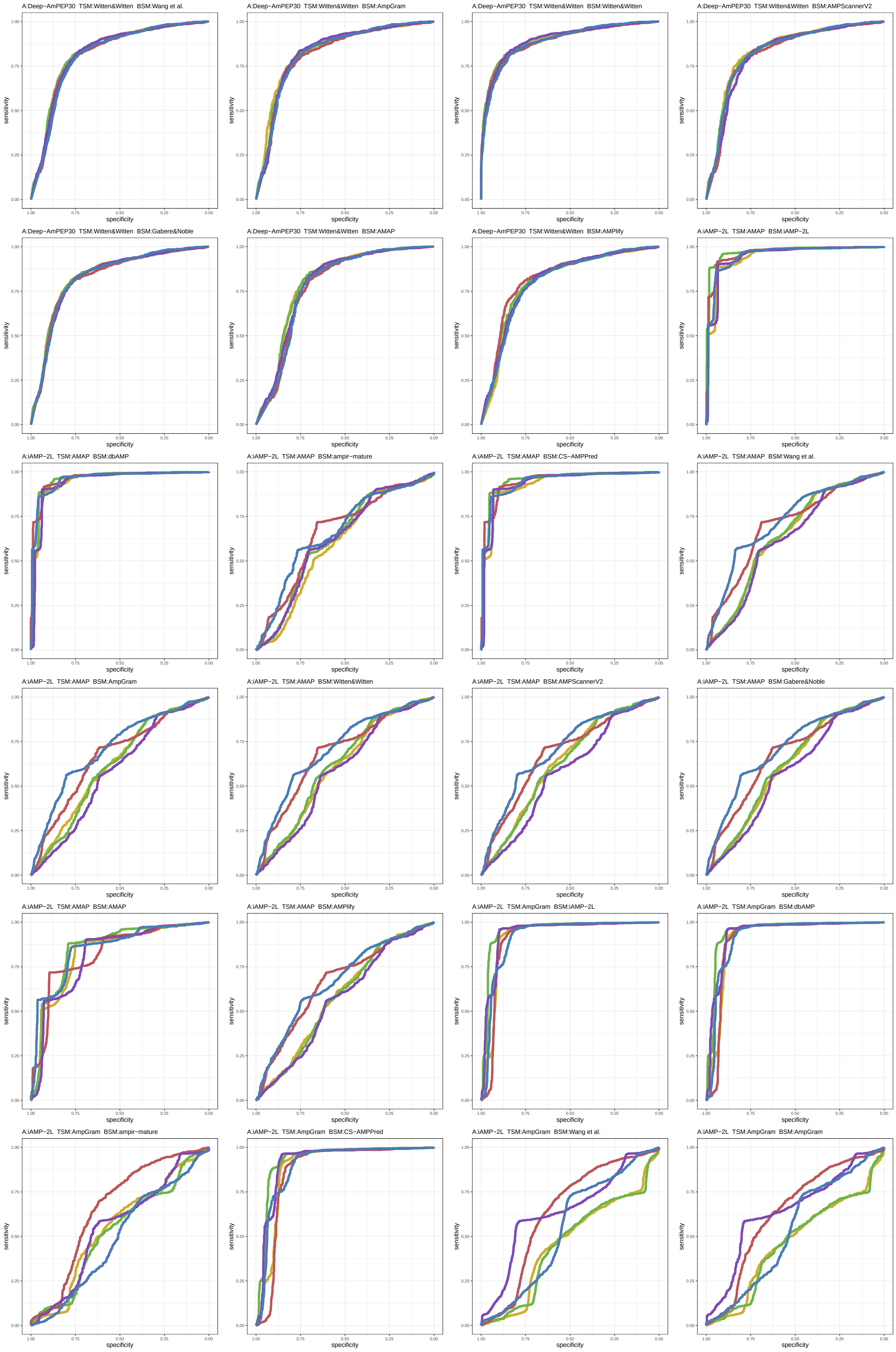

Figure S44: ROC curves 841-864 of 1452. Each subplot presents results for five replications indicated by different line colors.

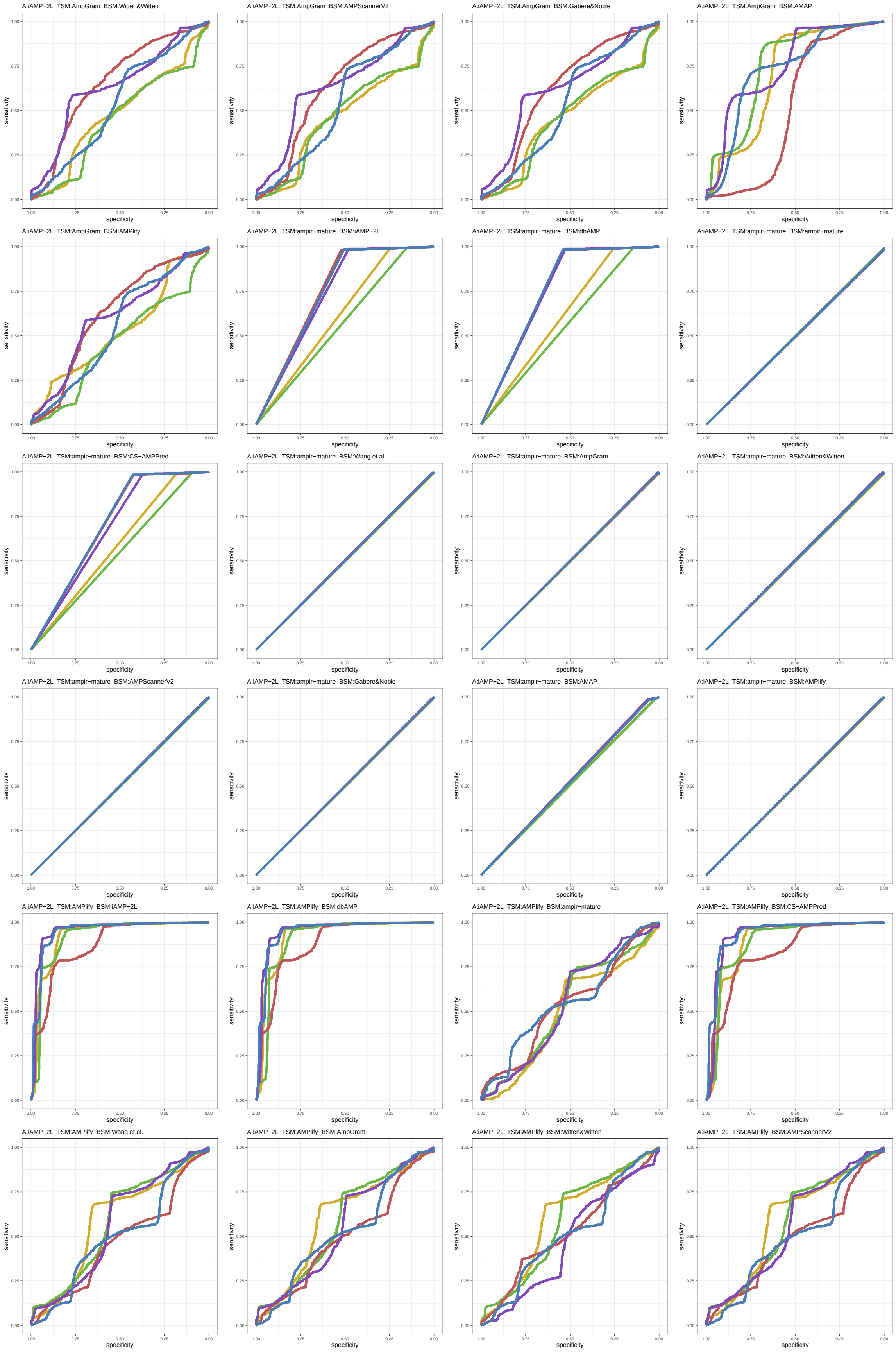

Figure S45: ROC curves 865-888 of 1452. Each subplot presents results for five replications indicated by different line colors.

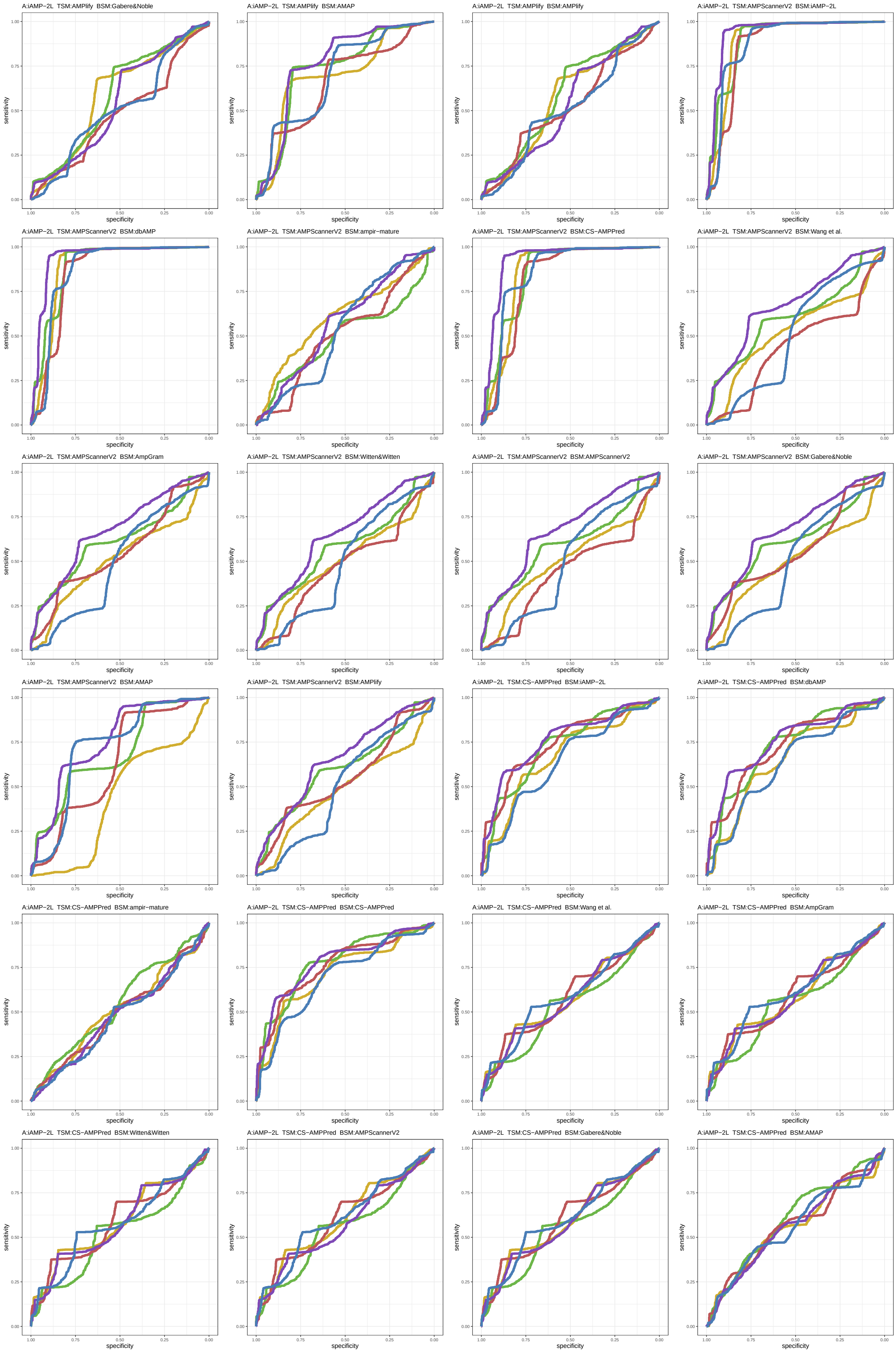

Figure S46: ROC curves 889-912 of 1452. Each subplot presents results for five replications indicated by different line colors.

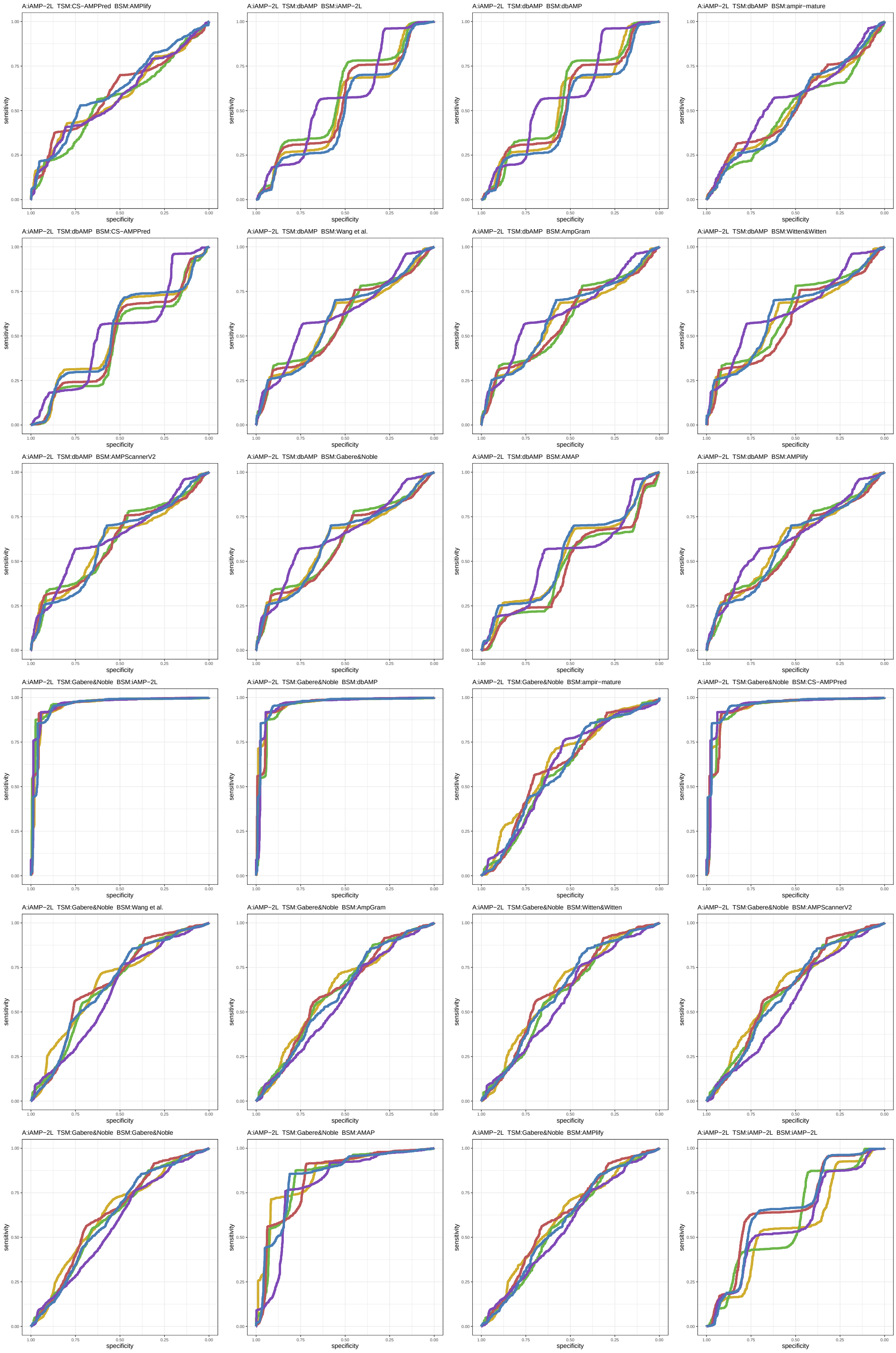

Figure S47: ROC curves 913-936 of 1452. Each subplot presents results for five replications indicated by different line colors.

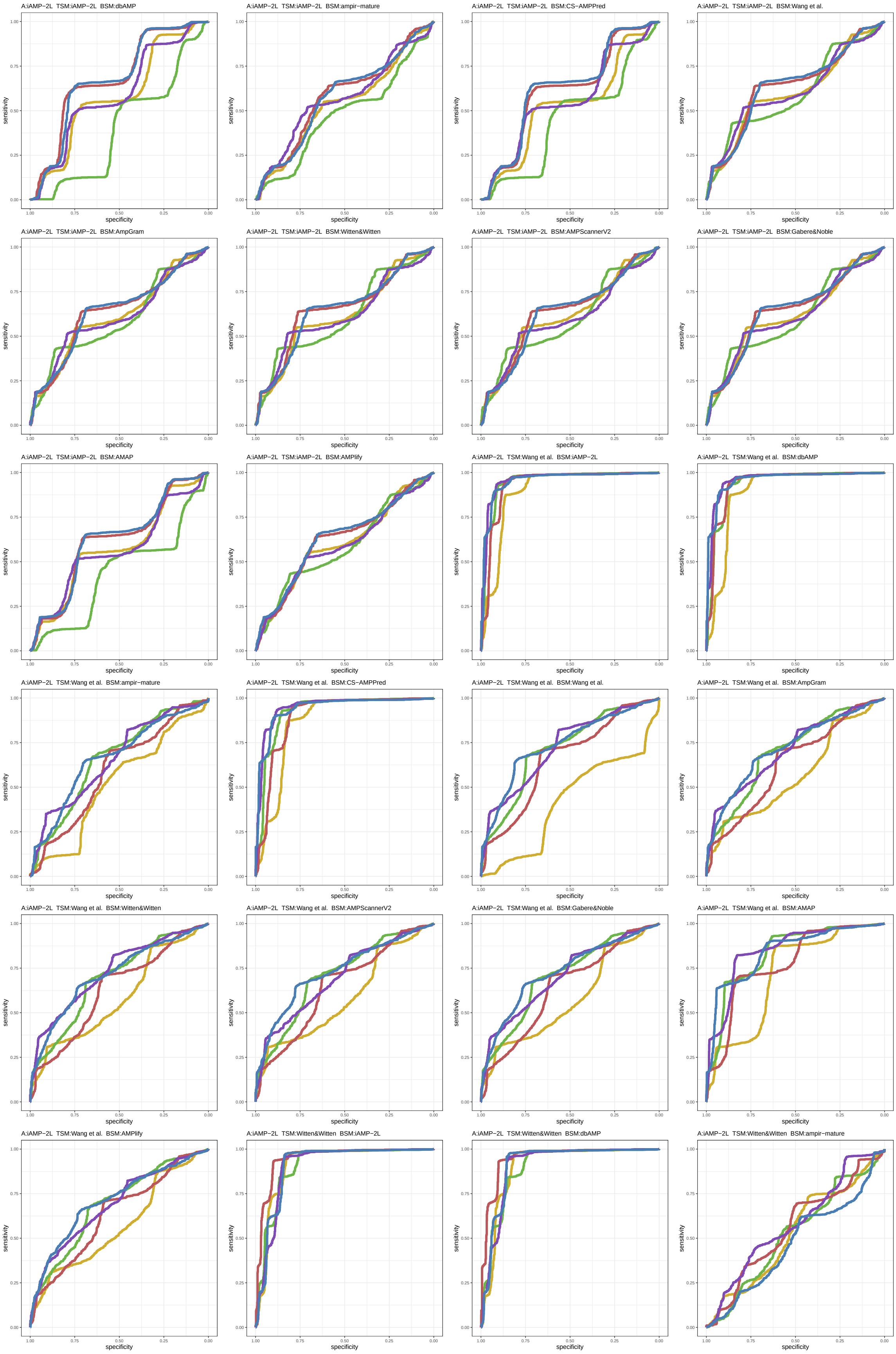

Figure S48: ROC curves 937-960 of 1452. Each subplot presents results for five replications indicated by different line colors.

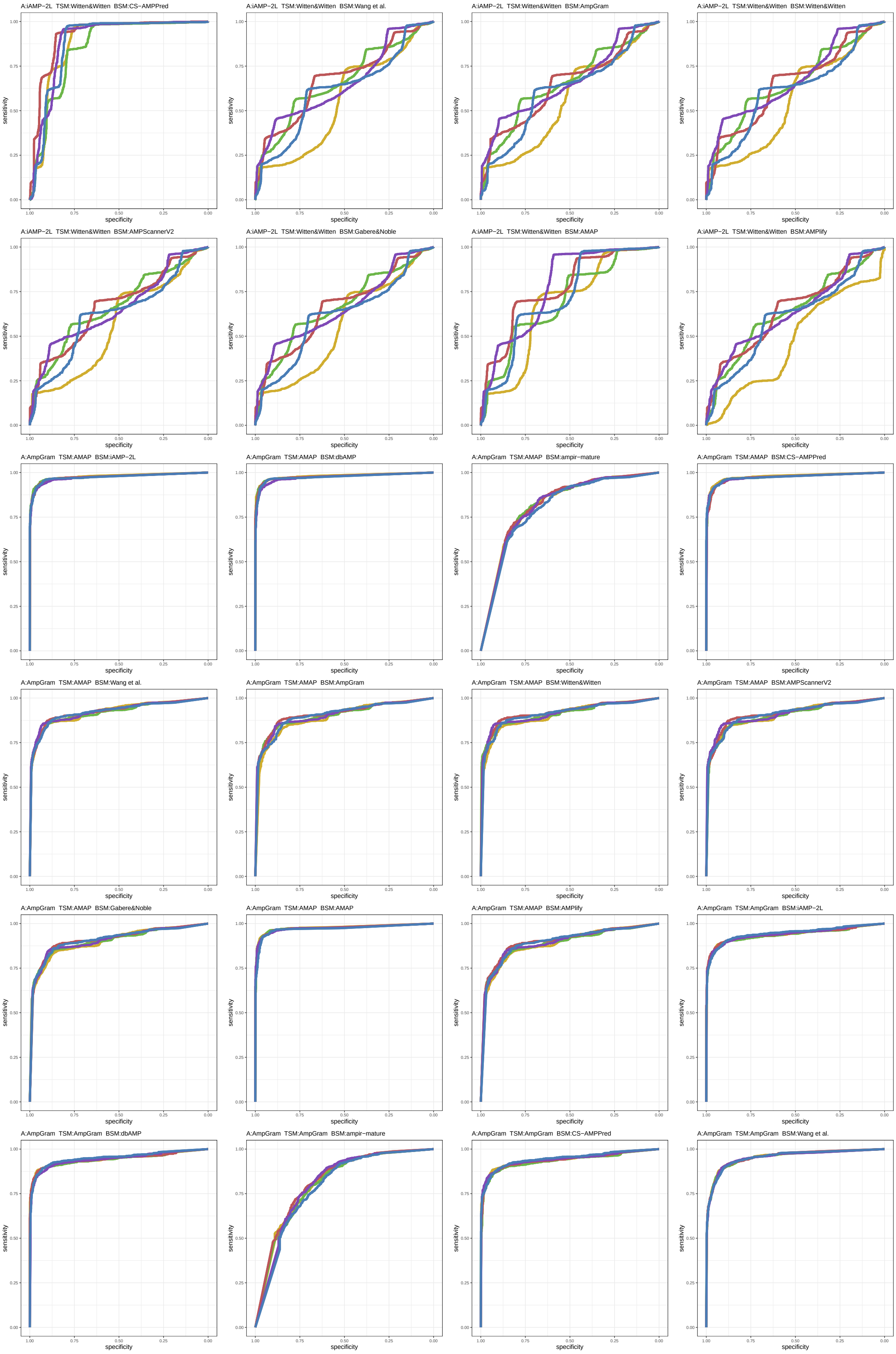

Figure S49: ROC curves 961-984 of 1452. Each subplot presents results for five replications indicated by different line colors.

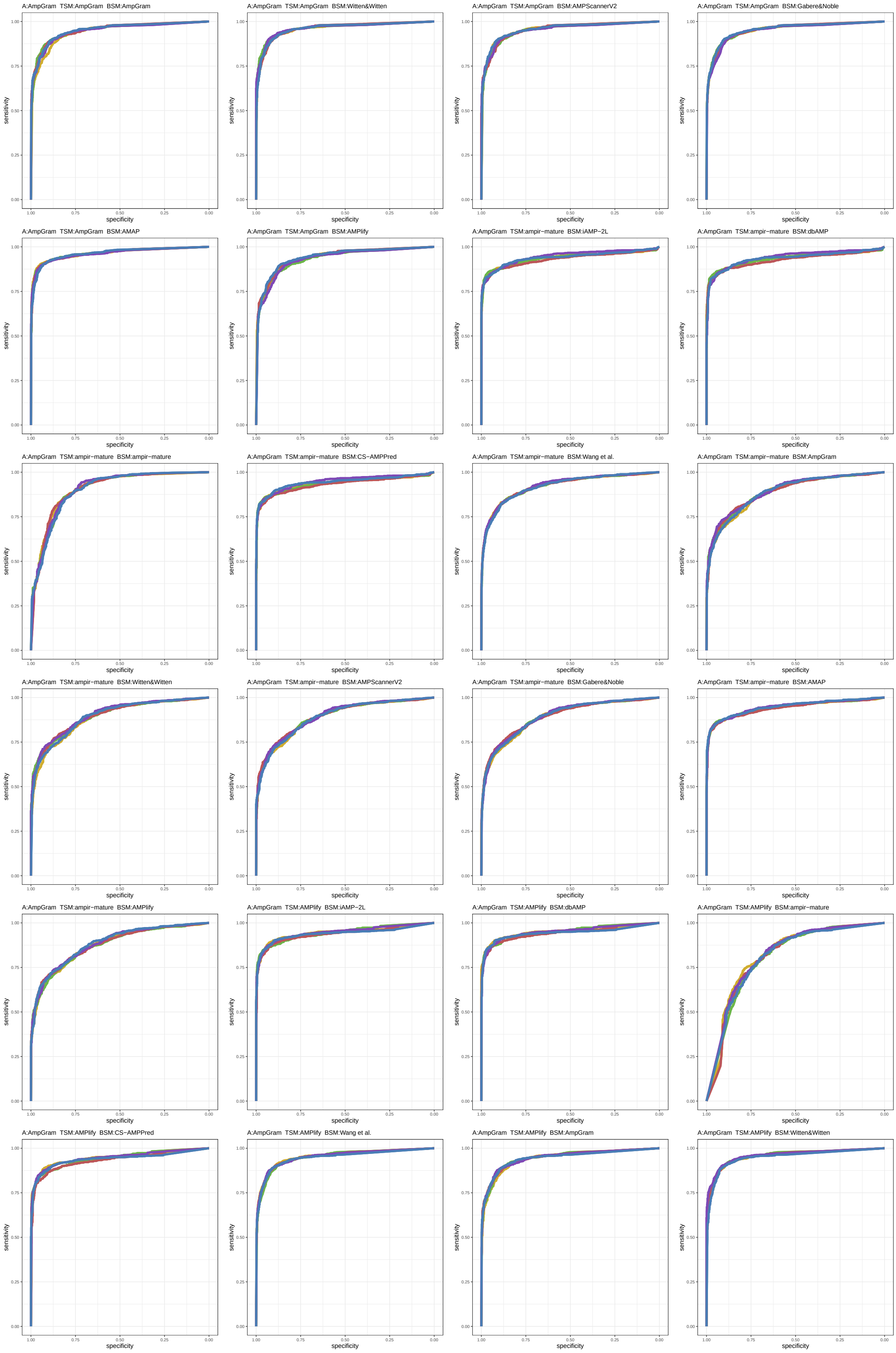

Figure S50: ROC curves 985-1008 of 1452. Each subplot presents results for five replications indicated by different line colors.

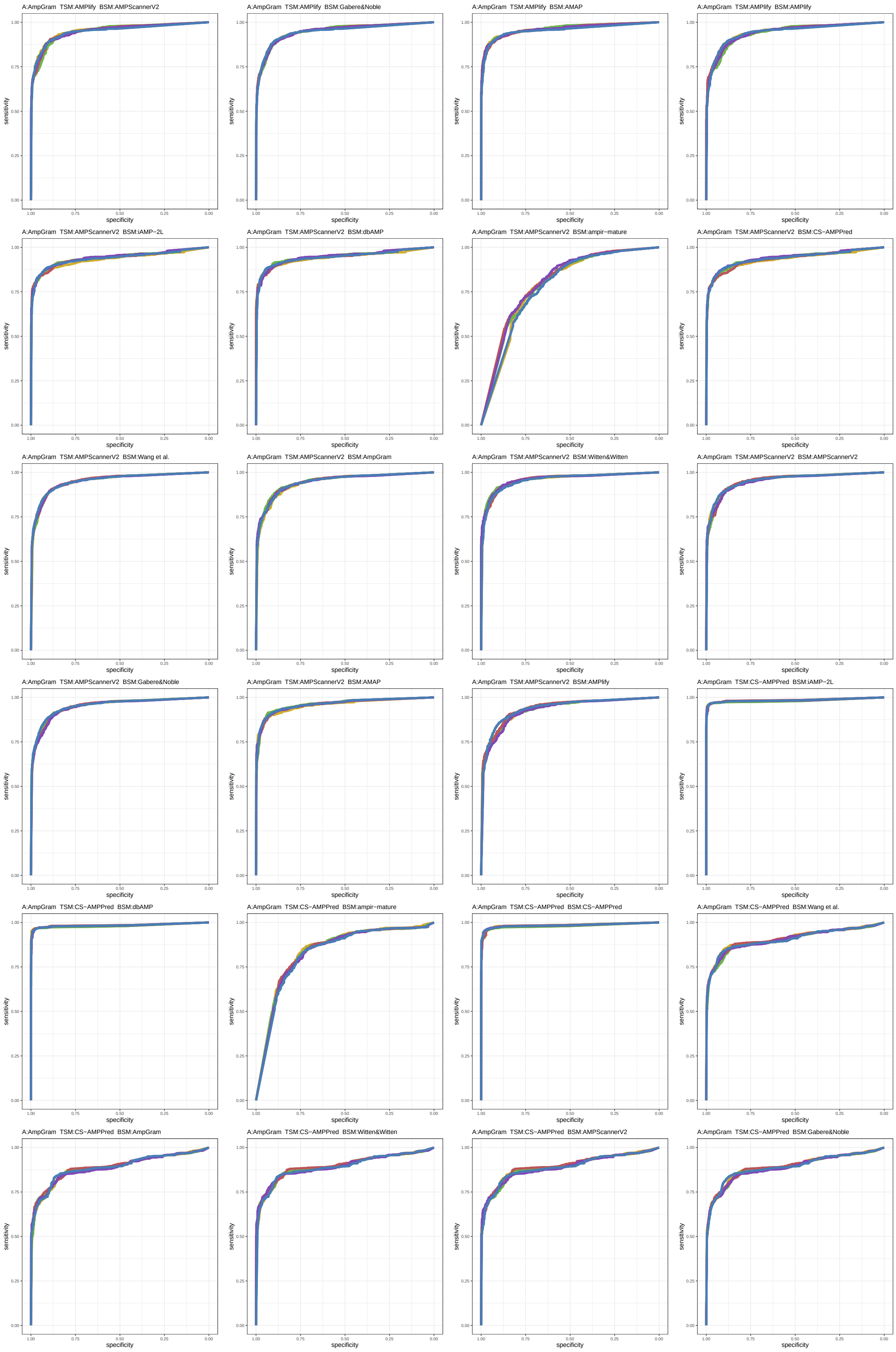

Figure S51: ROC curves 1009-1032 of 1452. Each subplot presents results for five replications indicated by different line colors.

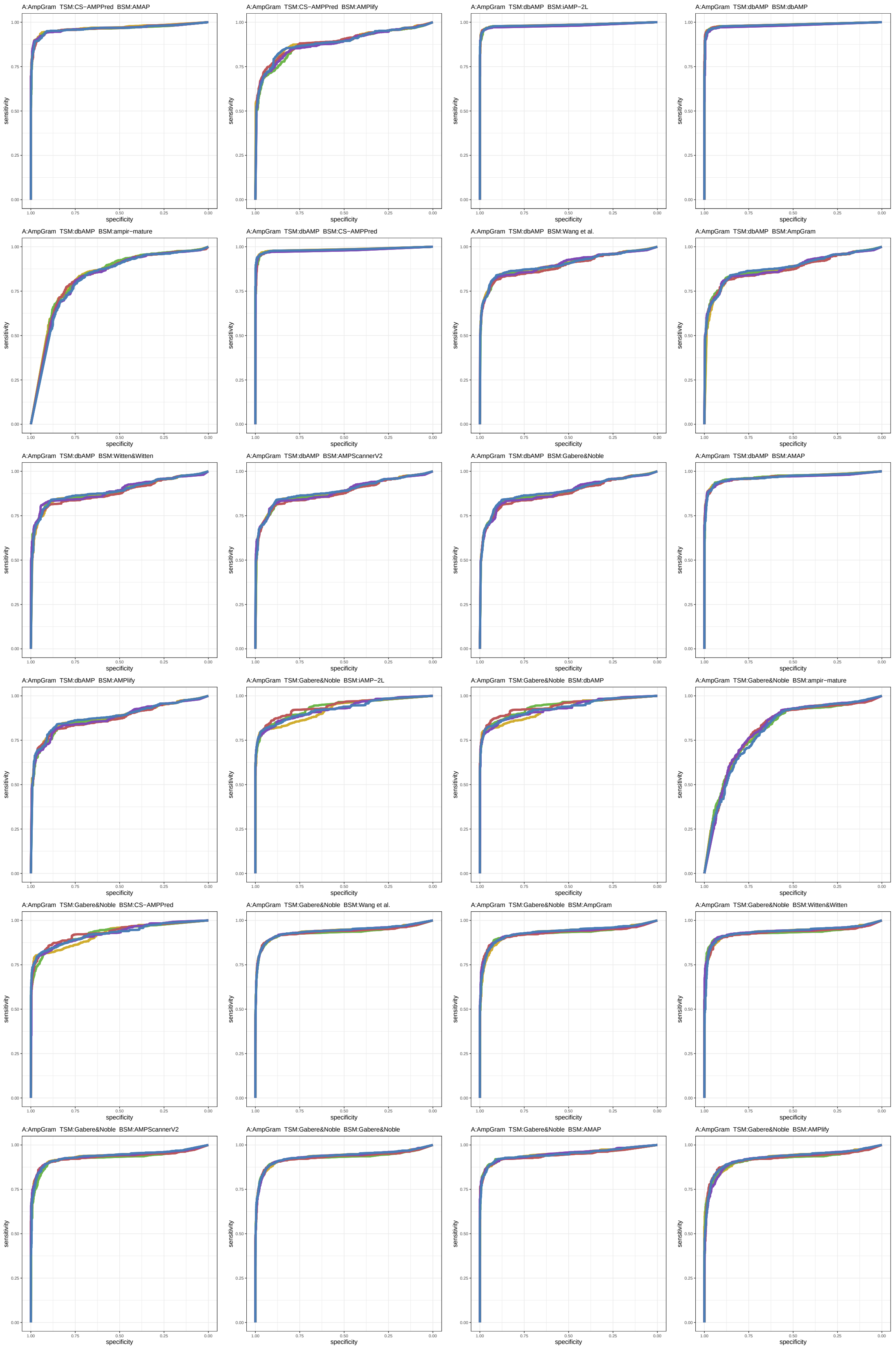

Figure S52: ROC curves 1033-1056 of 1452. Each subplot presents results for five replications indicated by different line colors.

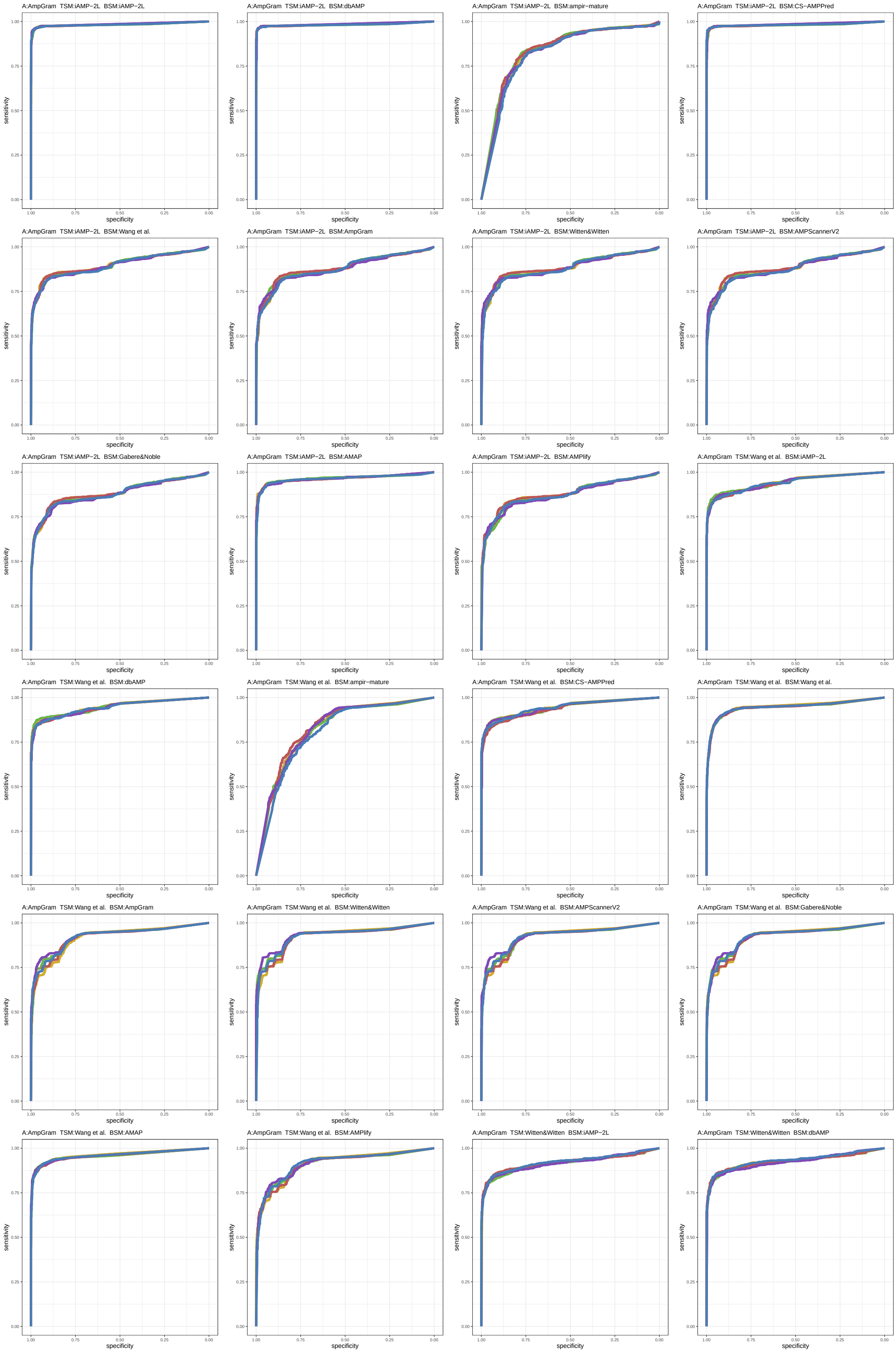

Figure S53: ROC curves 1057-1080 of 1452. Each subplot presents results for five replications indicated by different line colors.

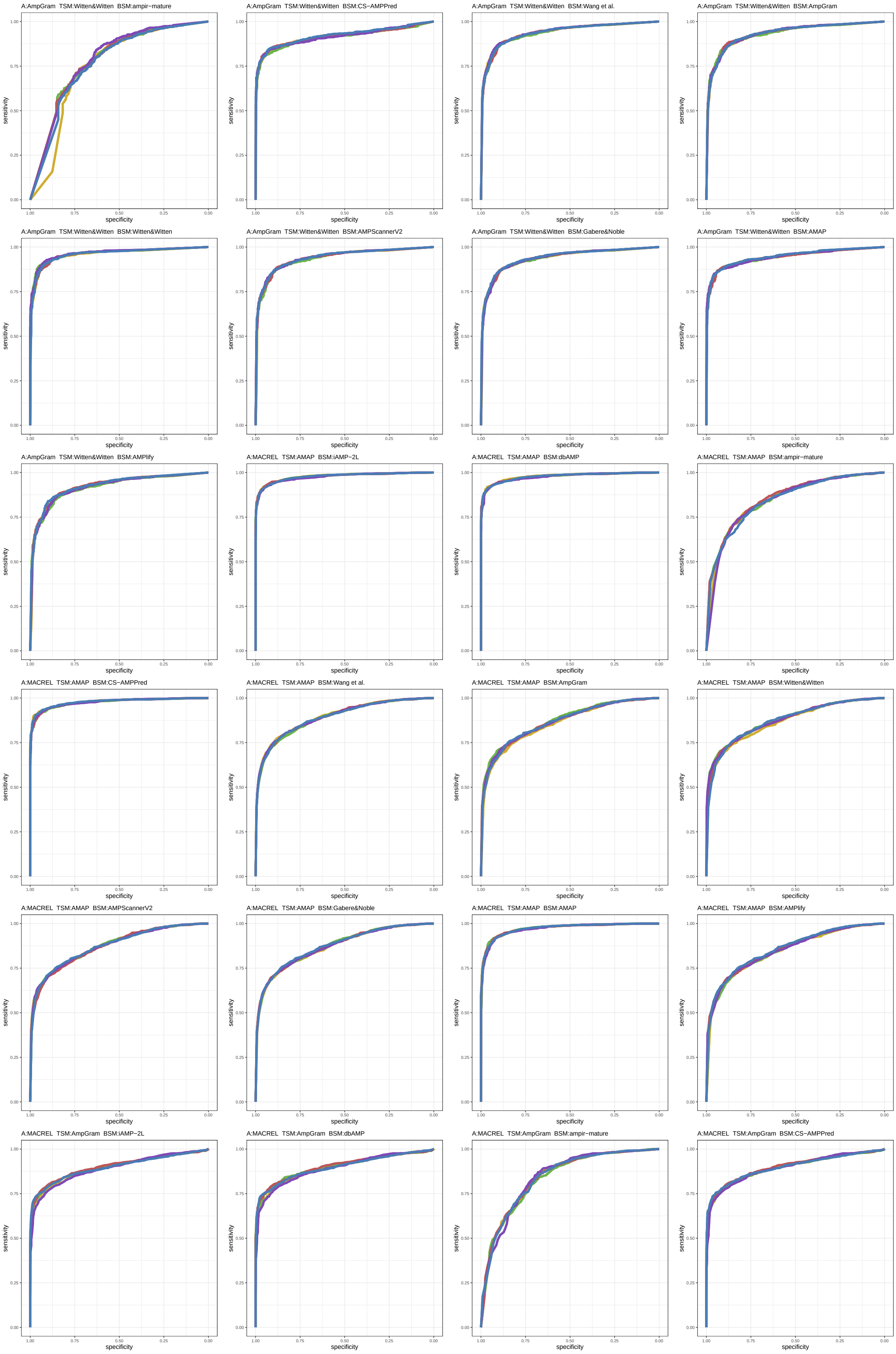

Figure S54: ROC curves 1081-1104 of 1452. Each subplot presents results for five replications indicated by different line colors.

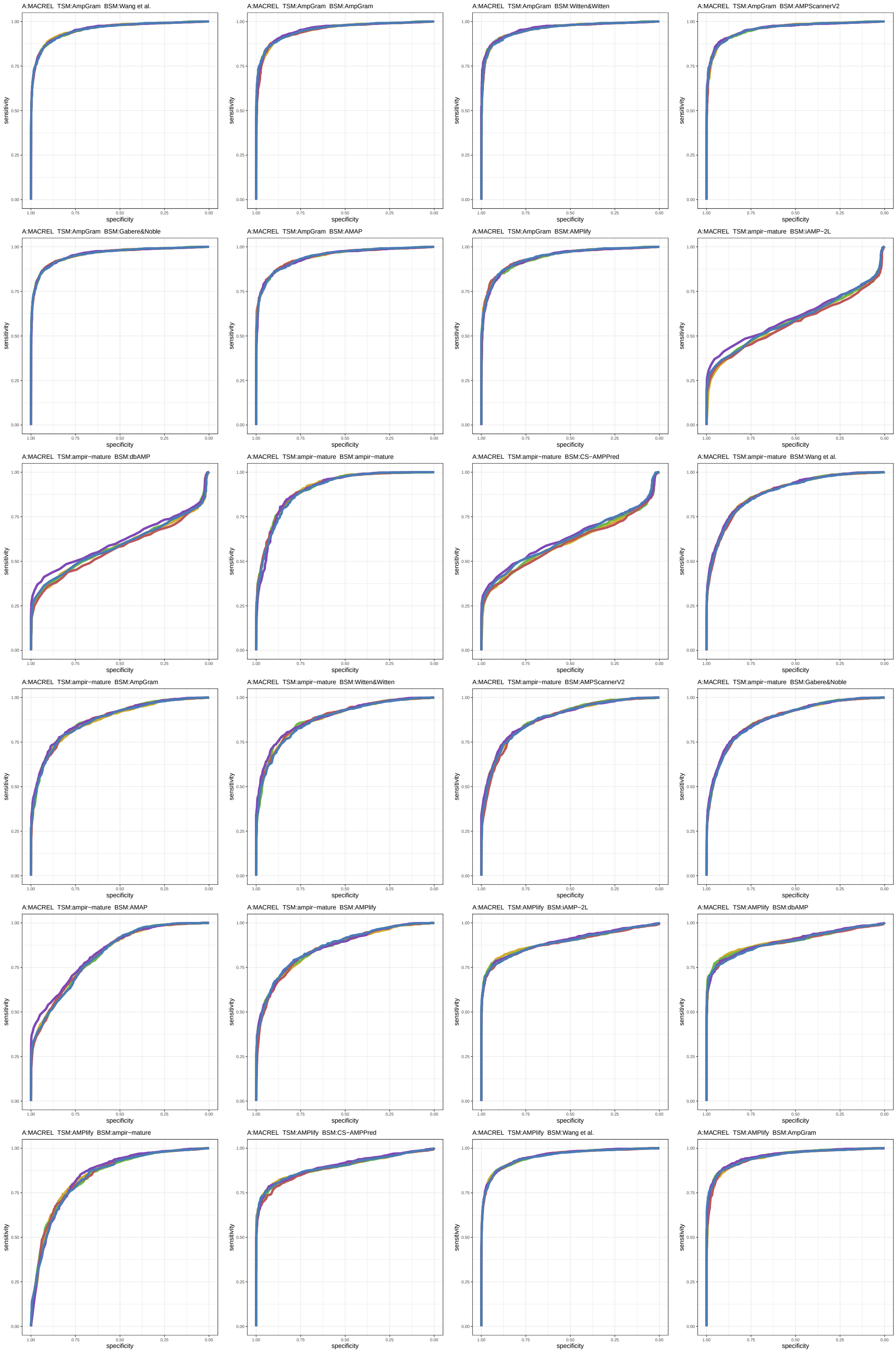

Figure S55: ROC curves 1105-1128 of 1452. Each subplot presents results for five replications indicated by different line colors.

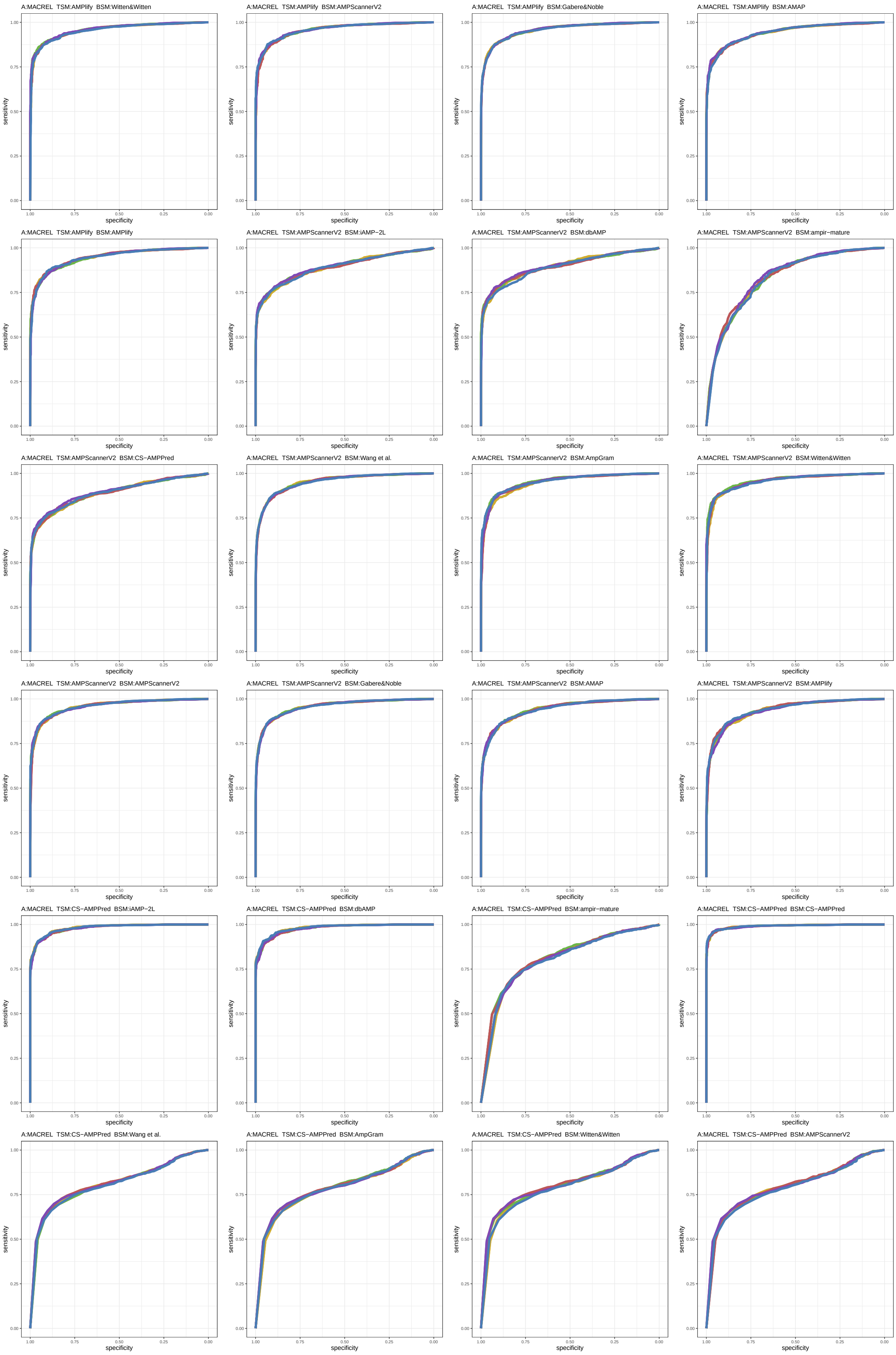

Figure S56: ROC curves 1129-1152 of 1452. Each subplot presents results for five replications indicated by different line colors.

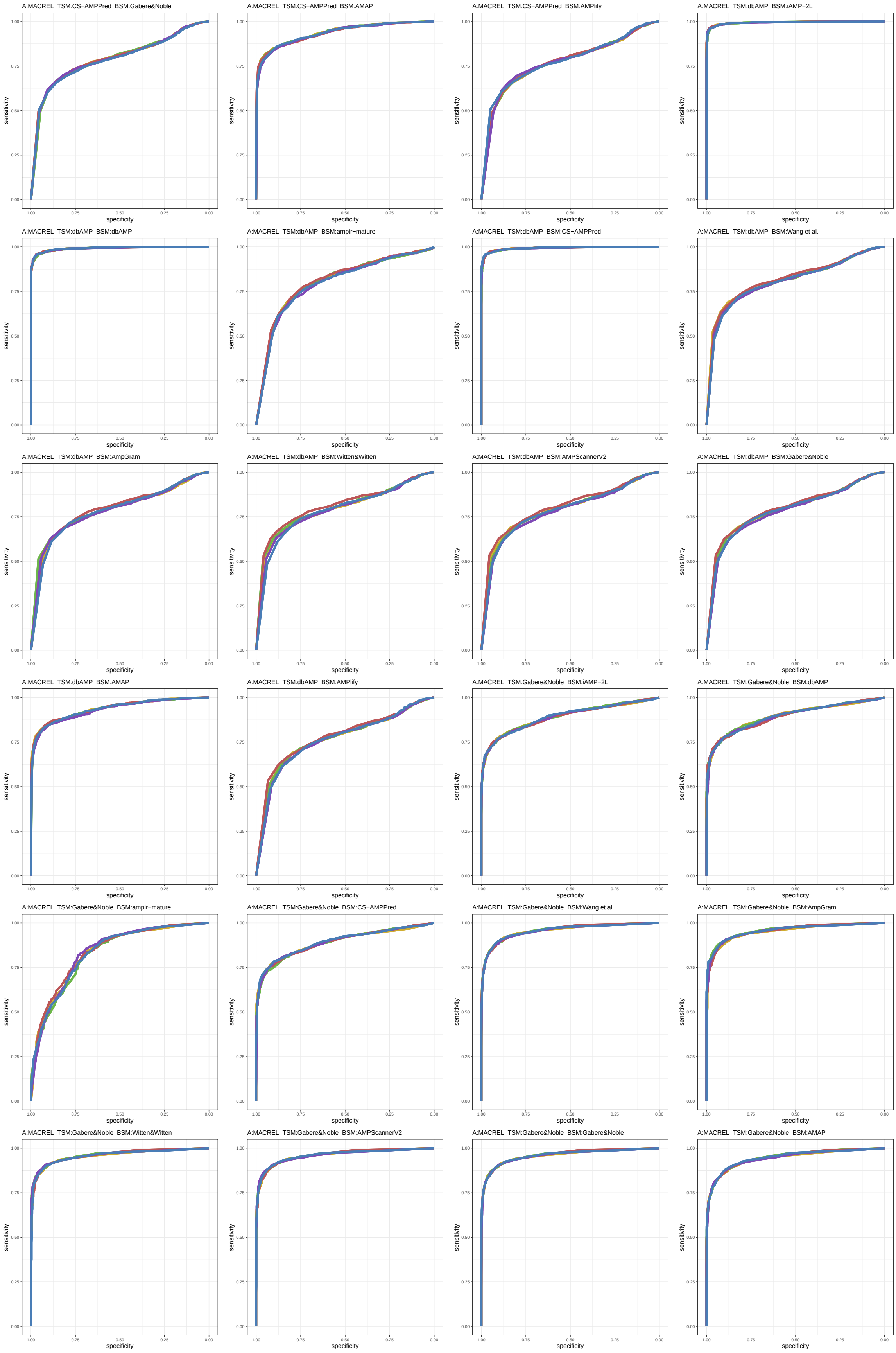

Figure S57: ROC curves 1153-1176 of 1452. Each subplot presents results for five replications indicated by different line colors.

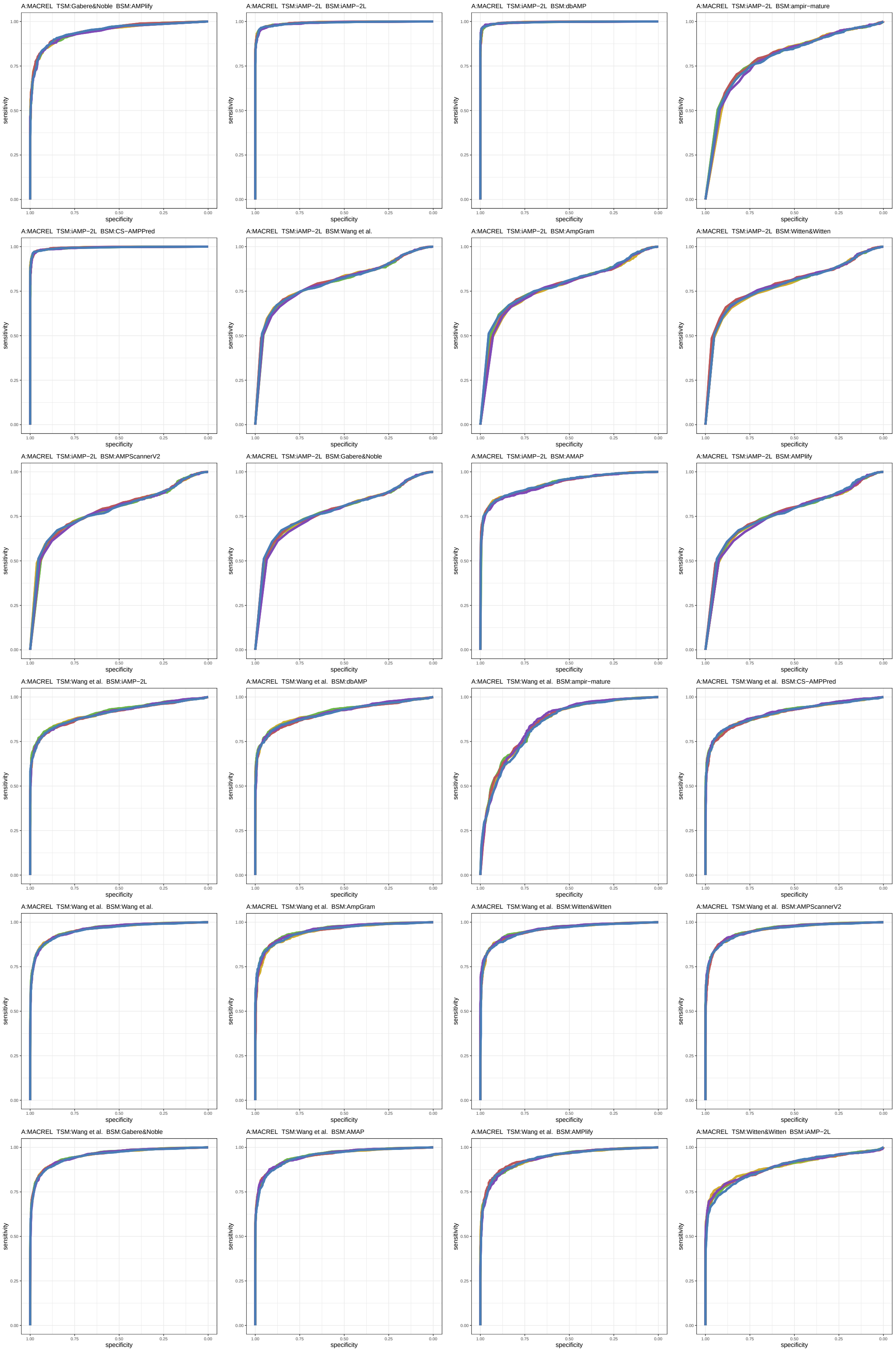

Figure S58: ROC curves 1177-1200 of 1452. Each subplot presents results for five replications indicated by different line colors.

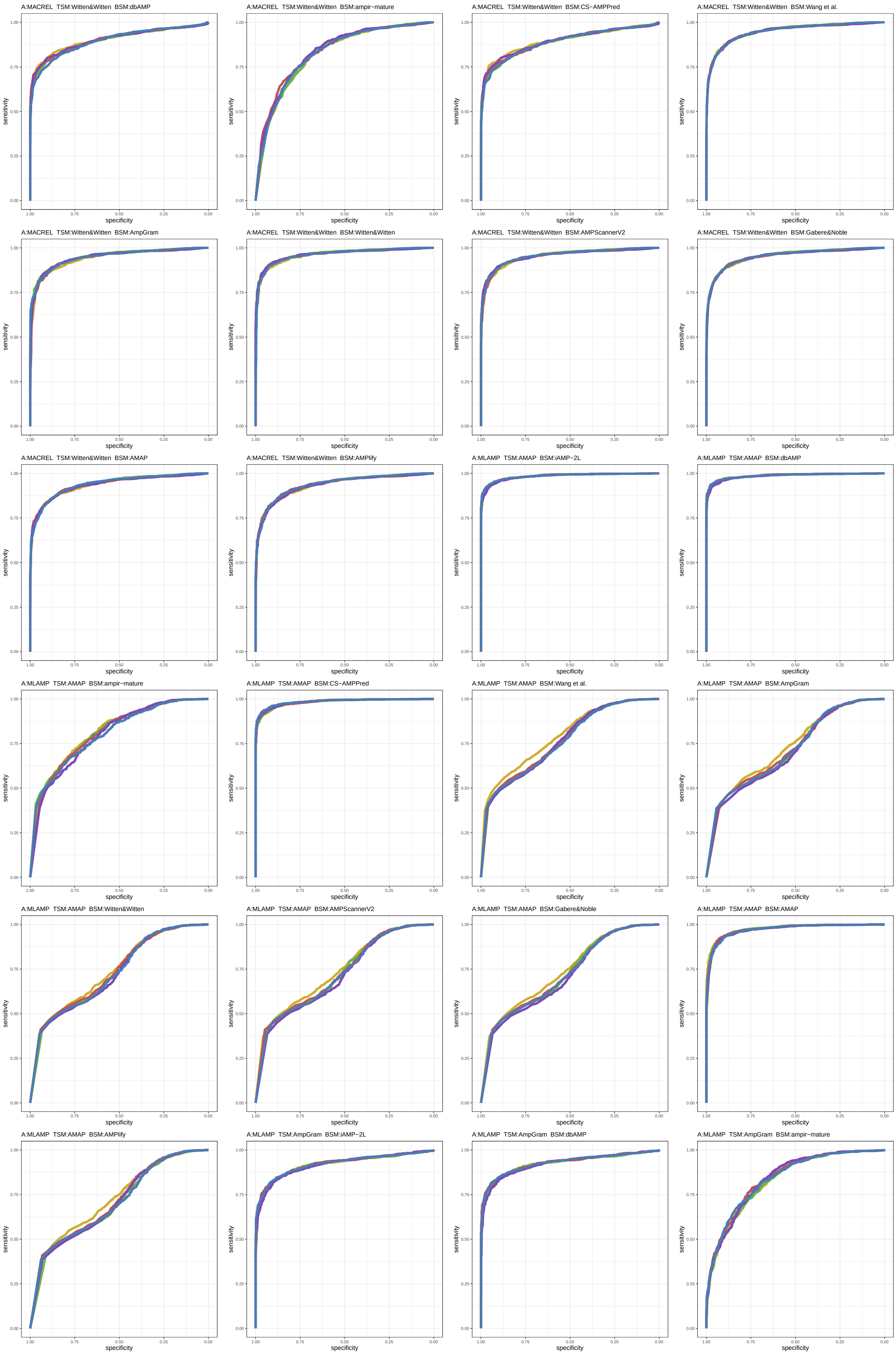

Figure S59: ROC curves 1201-1224 of 1452. Each subplot presents results for five replications indicated by different line colors.

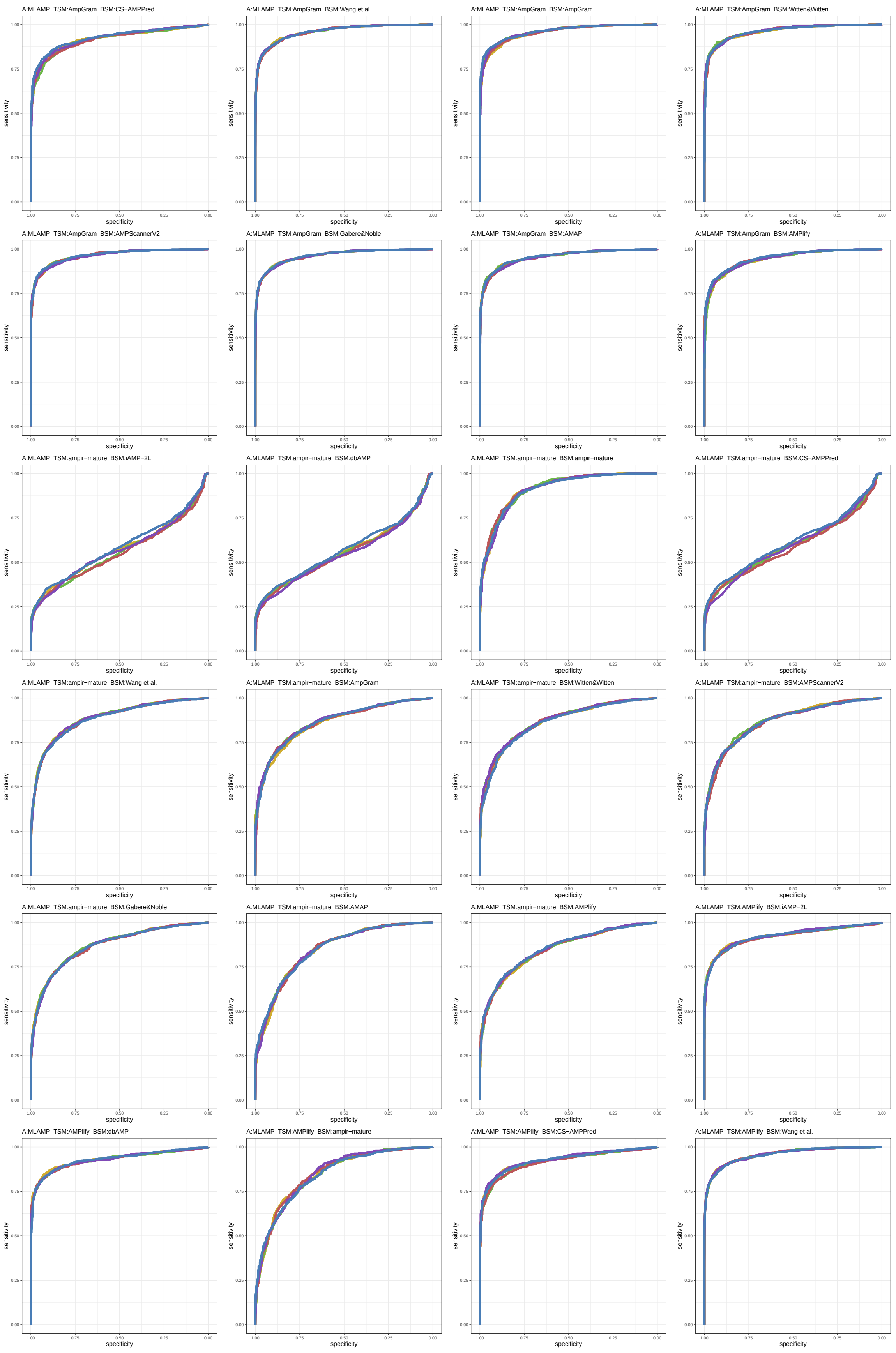

Figure S60: ROC curves 1225-1248 of 1452. Each subplot presents results for five replications indicated by different line colors.

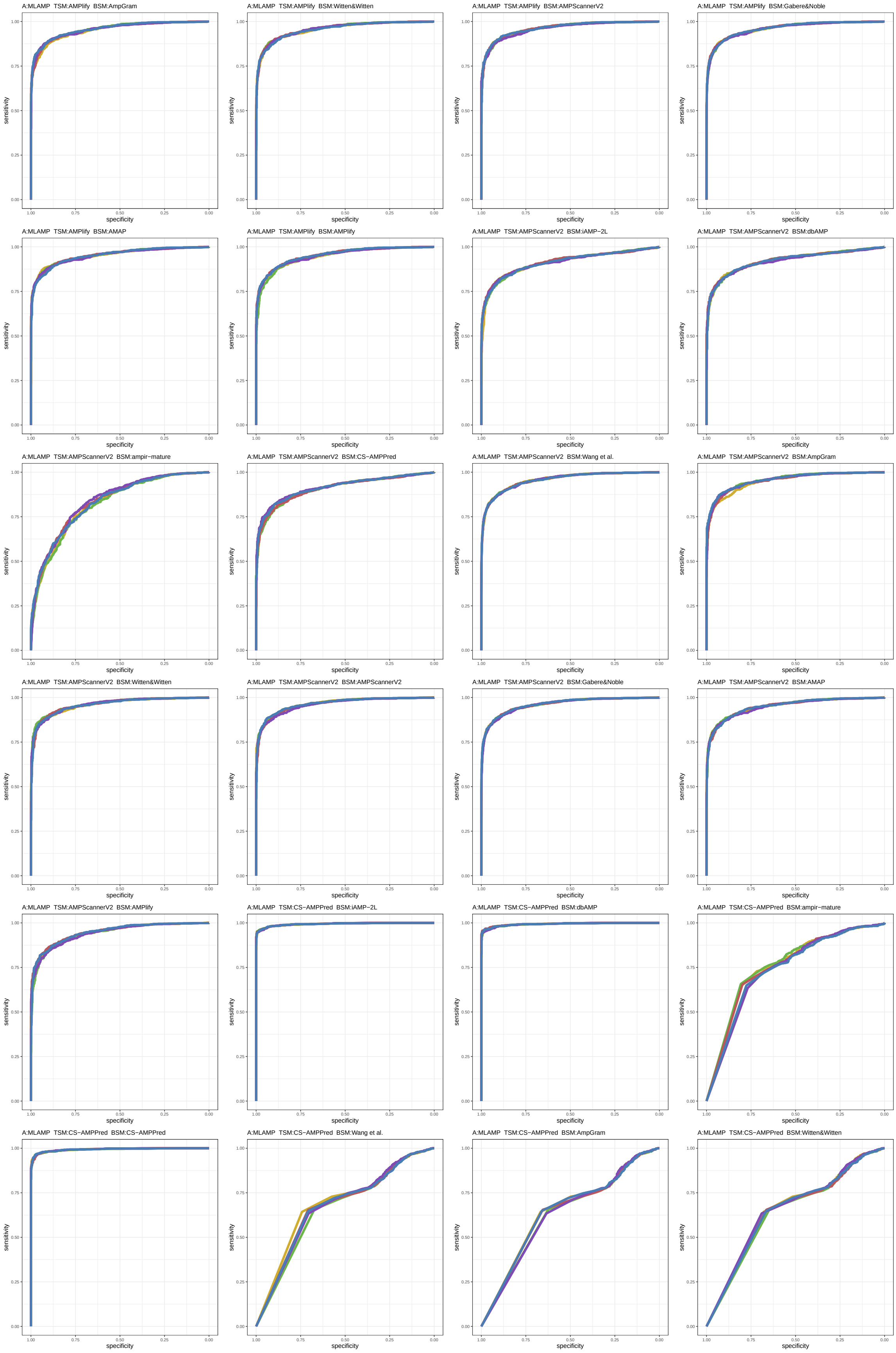

Figure S61: ROC curves 1249-1272 of 1452. Each subplot presents results for five replications indicated by different line colors.

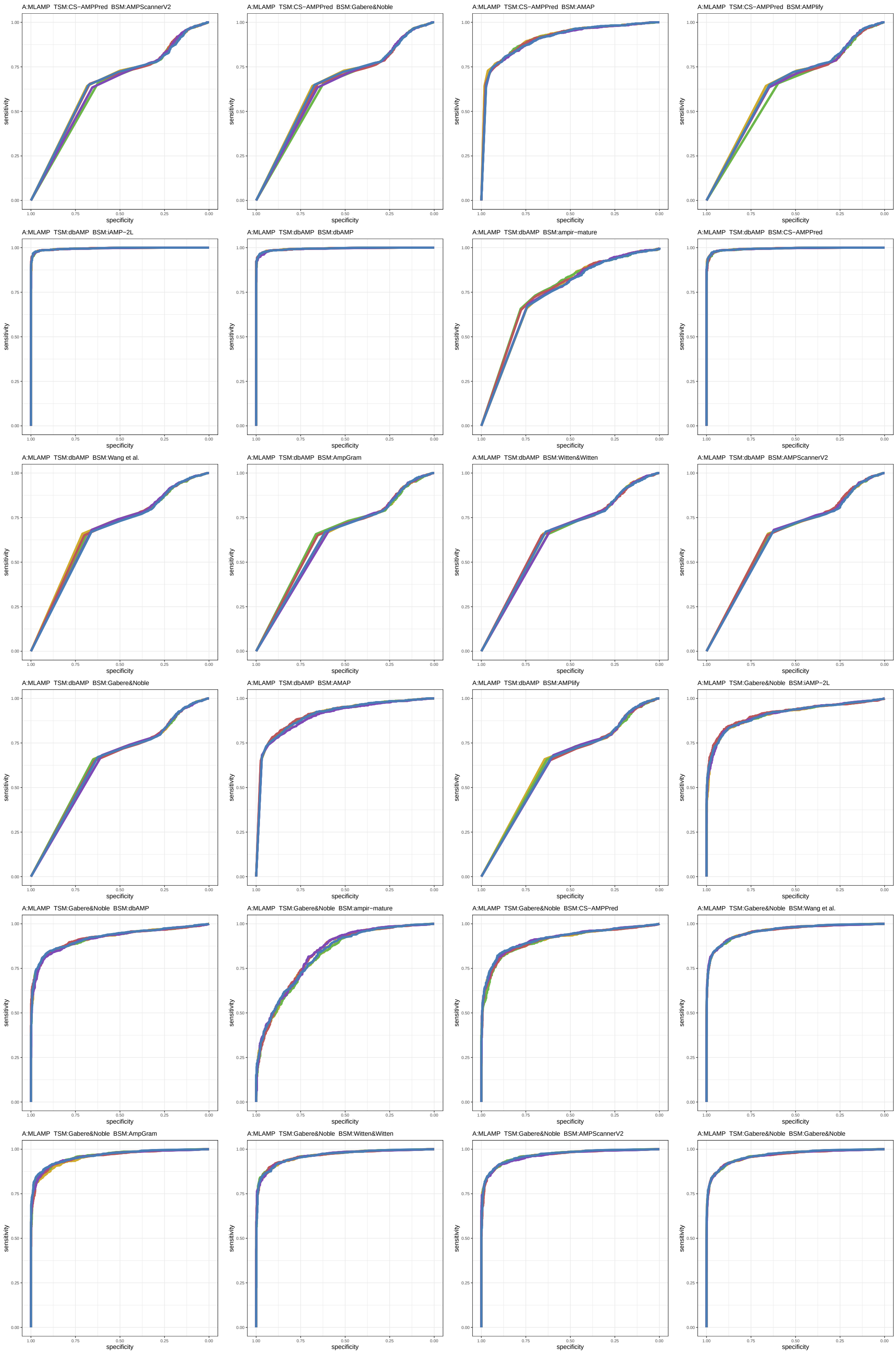

Figure S62: ROC curves 1273-1296 of 1452. Each subplot presents results for five replications indicated by different line colors.

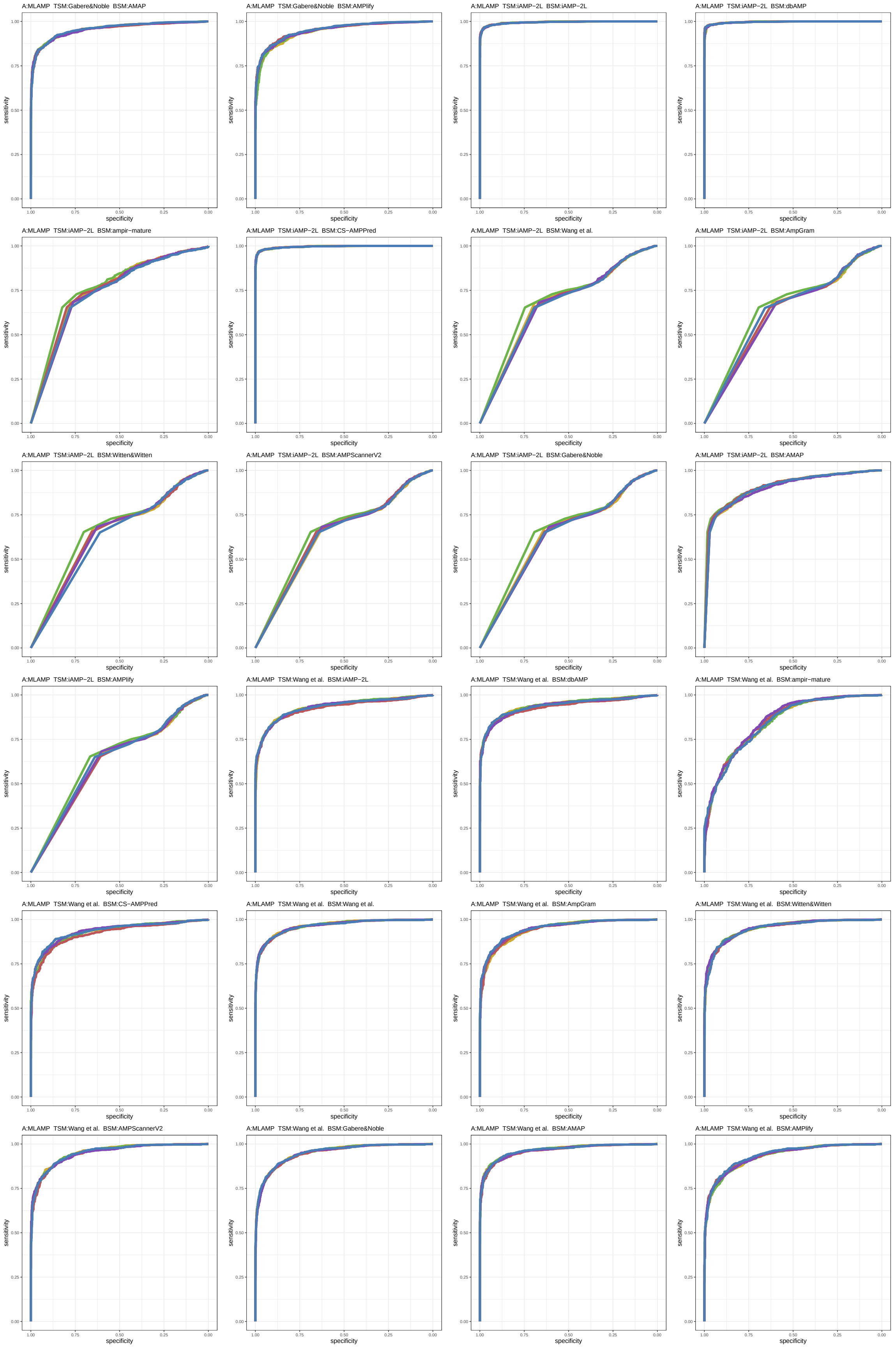

Figure S63: ROC curves 1297-1320 of 1452. Each subplot presents results for five replications indicated by different line colors.

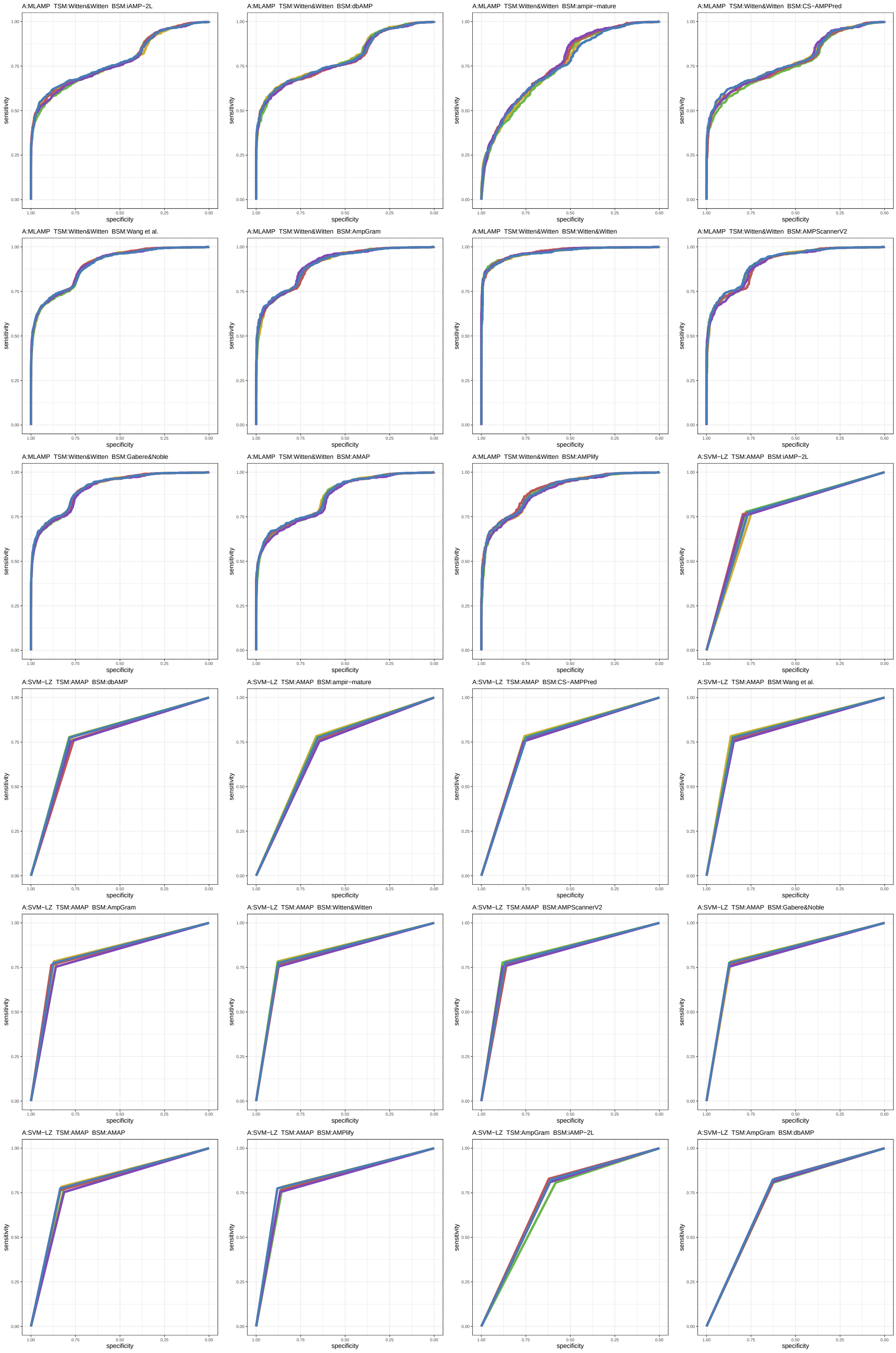

Figure S64: ROC curves 1321-1344 of 1452. Each subplot presents results for five replications indicated by different line colors.

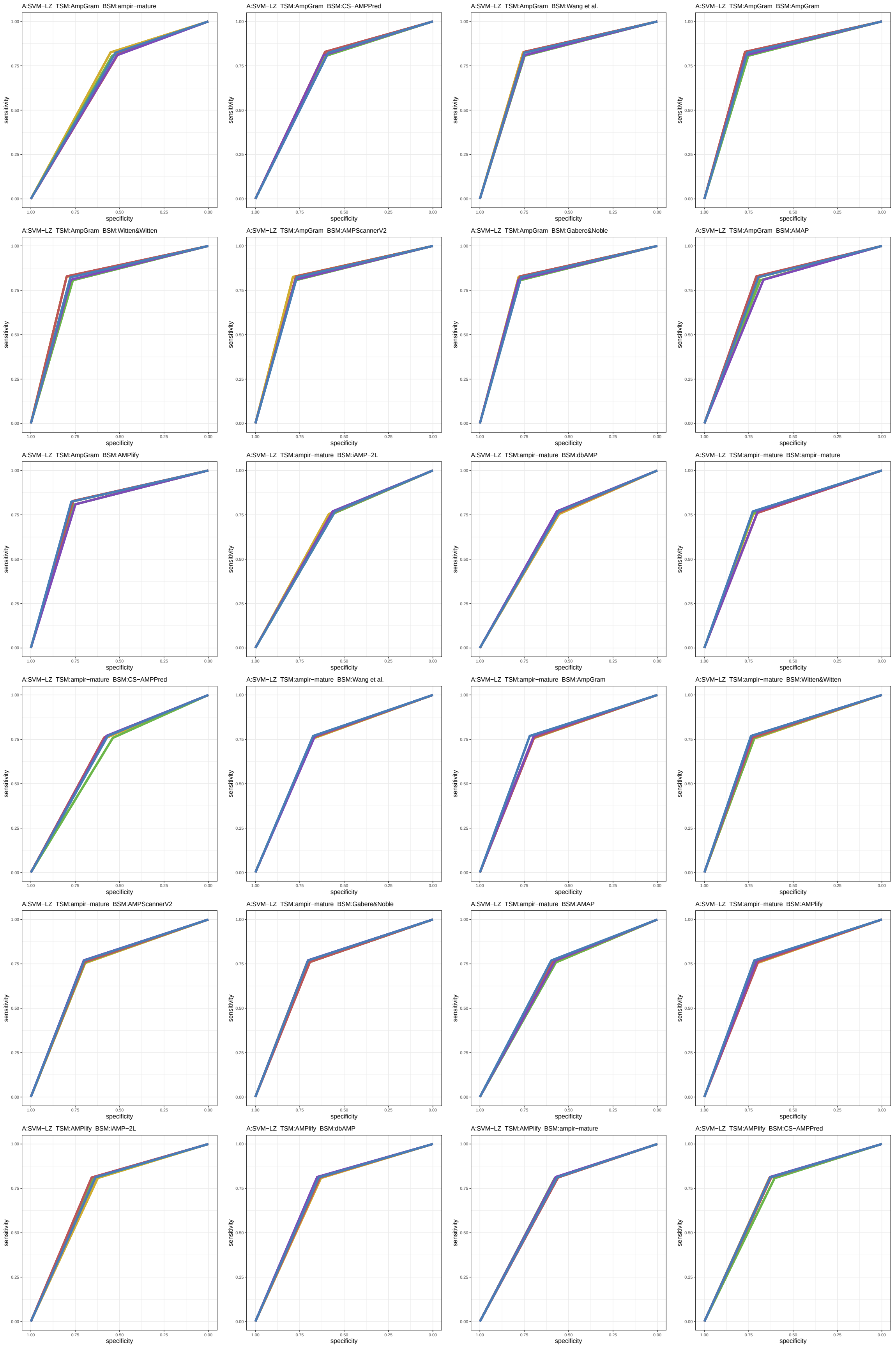

Figure S65: ROC curves 1345-1368 of 1452. Each subplot presents results for five replications indicated by different line colors.

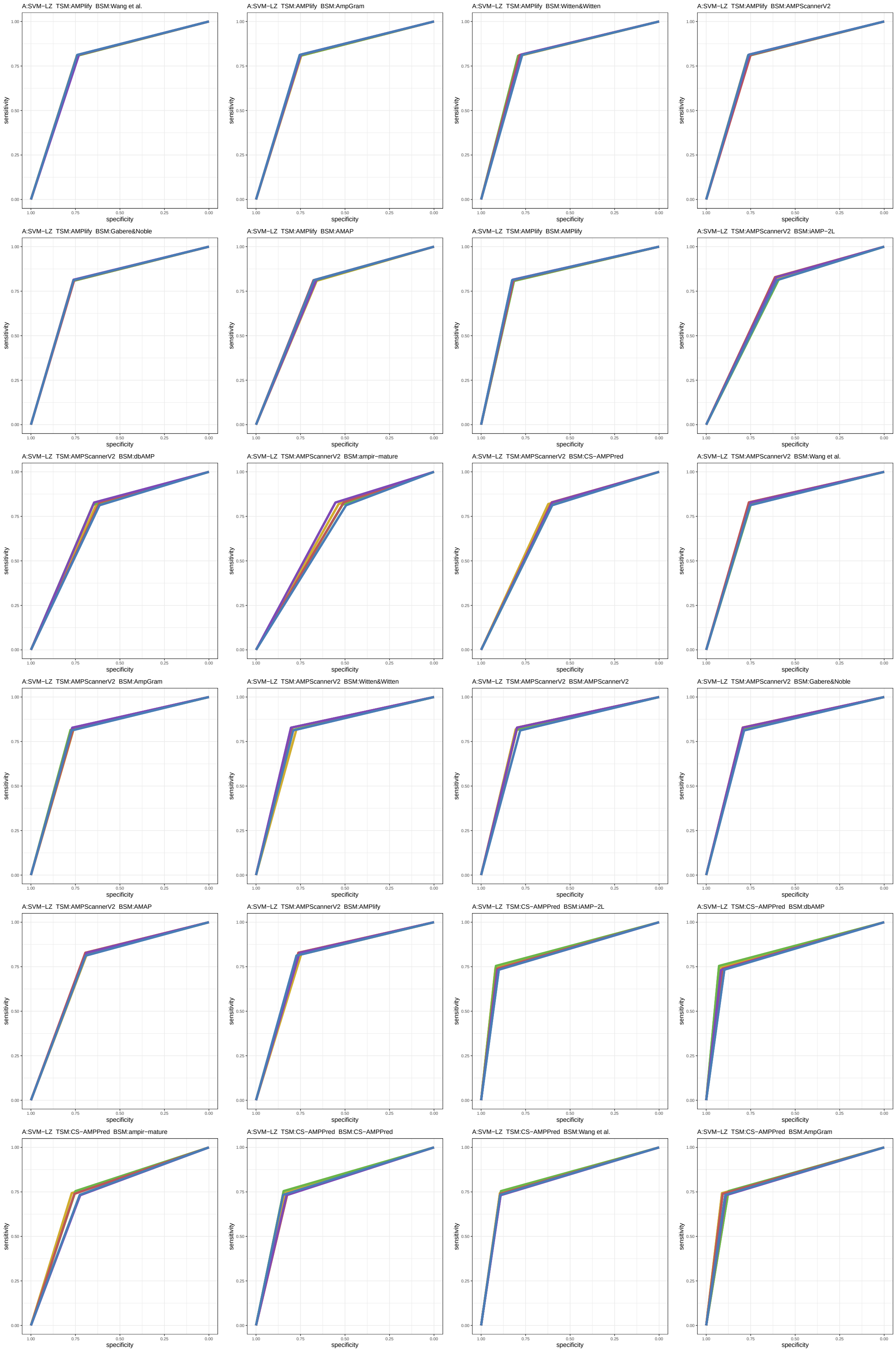

Figure S66: ROC curves 1369-1392 of 1452. Each subplot presents results for five replications indicated by different line colors.

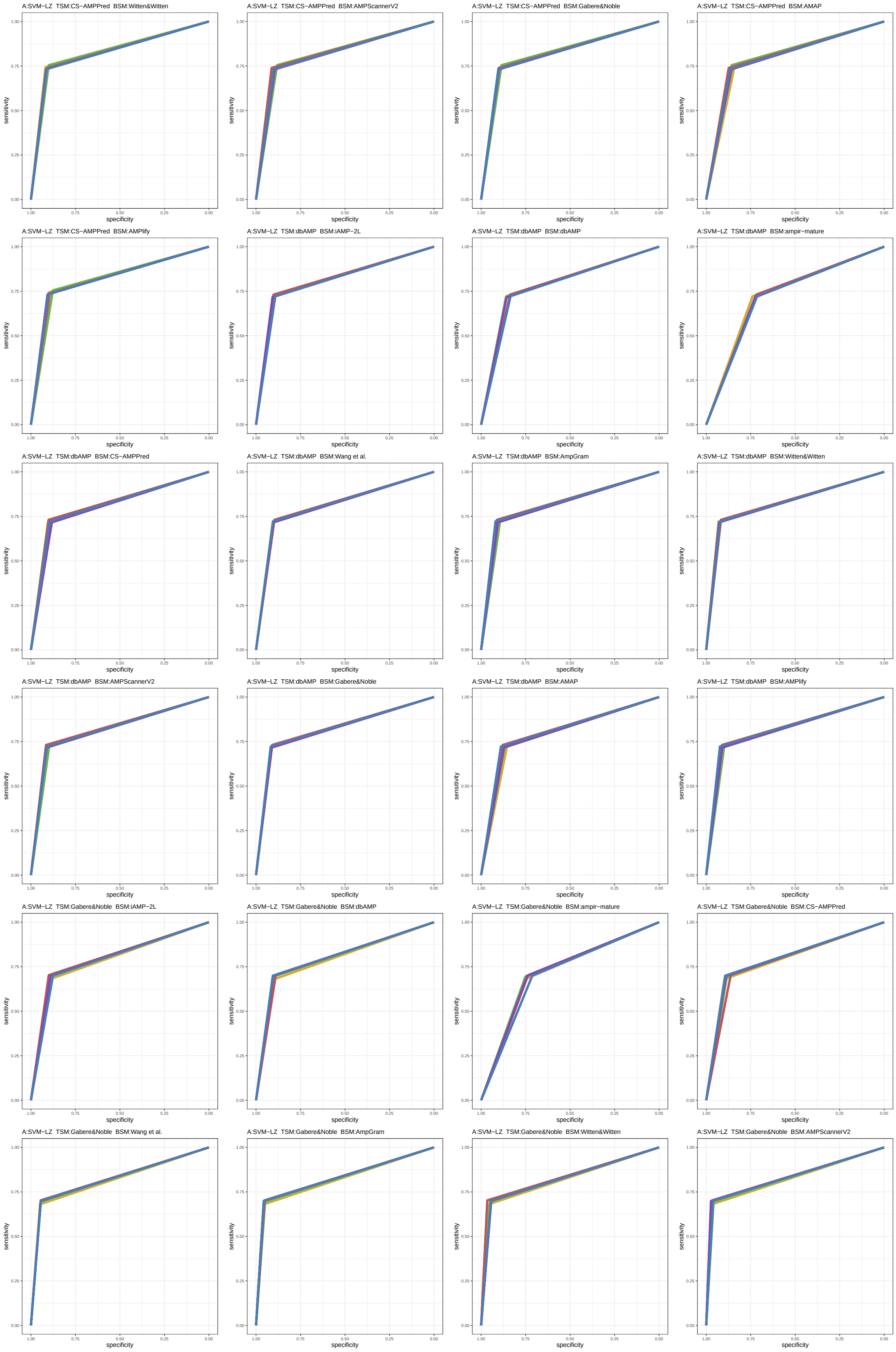

Figure S67: ROC curves 1393-1416 of 1452. Each subplot presents results for five replications indicated by different line colors.

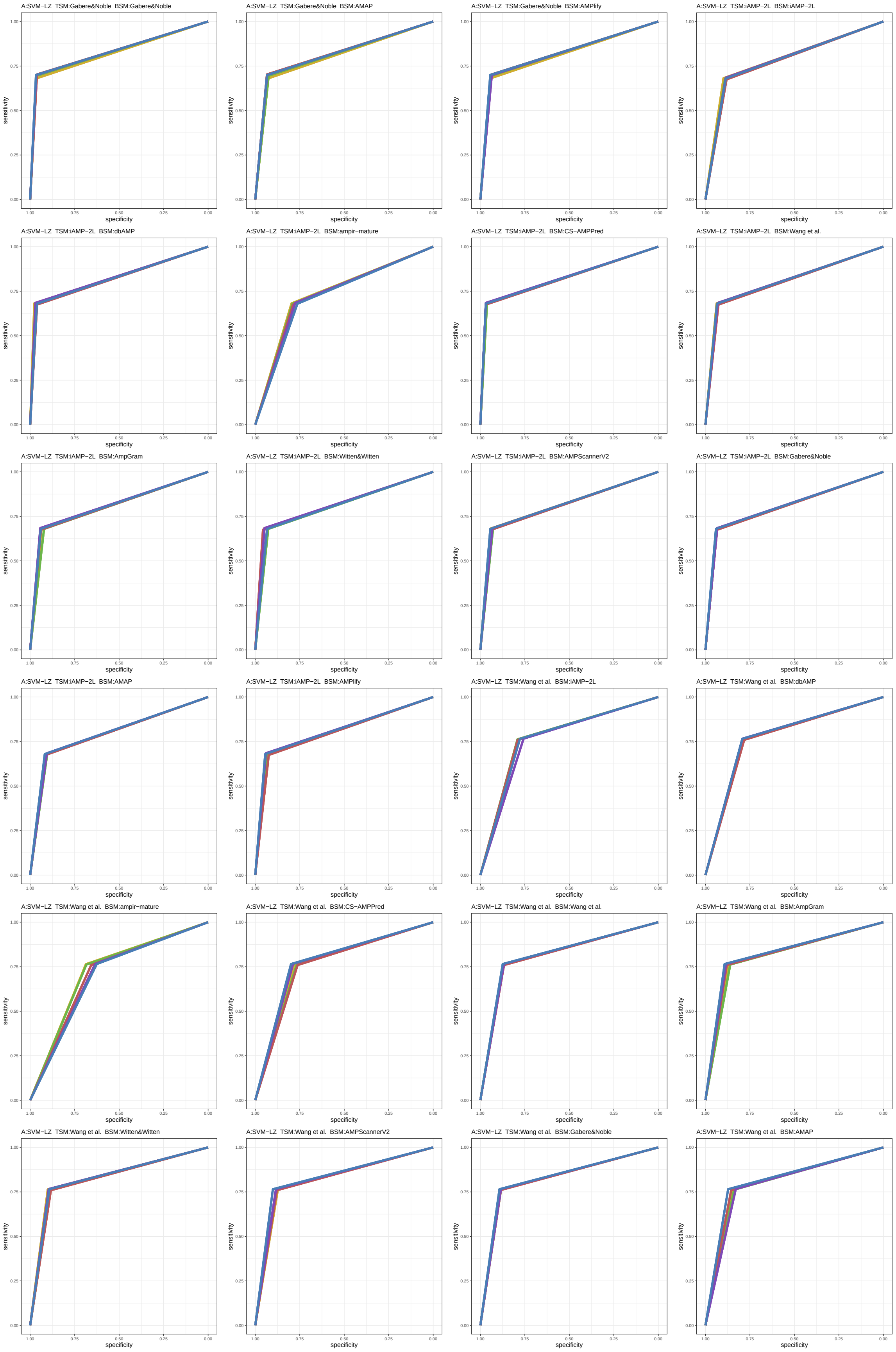

Figure S68: ROC curves 1417-1440 of 1452. Each subplot presents results for five replications indicated by different line colors.

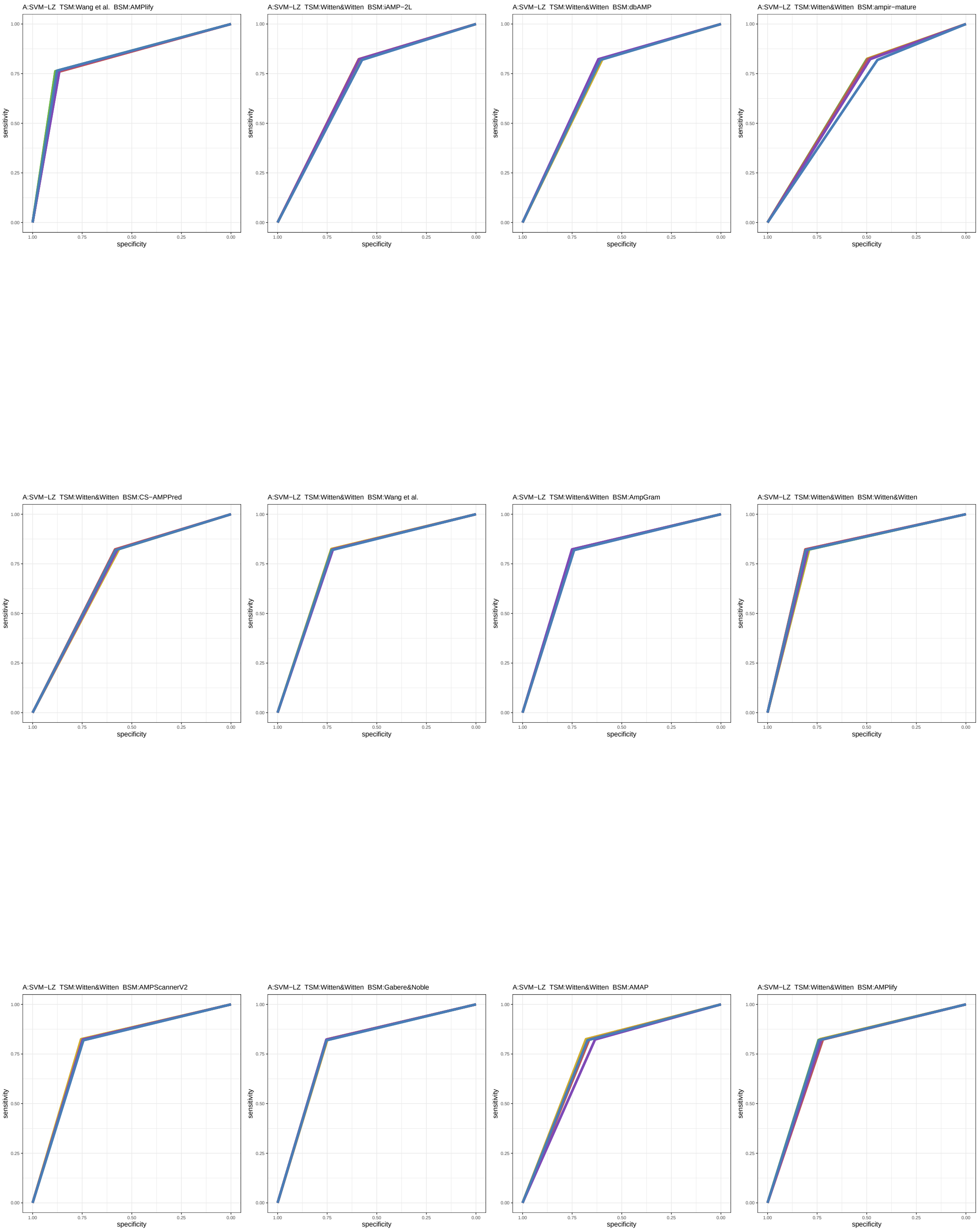

Figure S69: ROC curves 1441-1452 of 1452. Each subplot presents results for five replications indicated by different line colors.

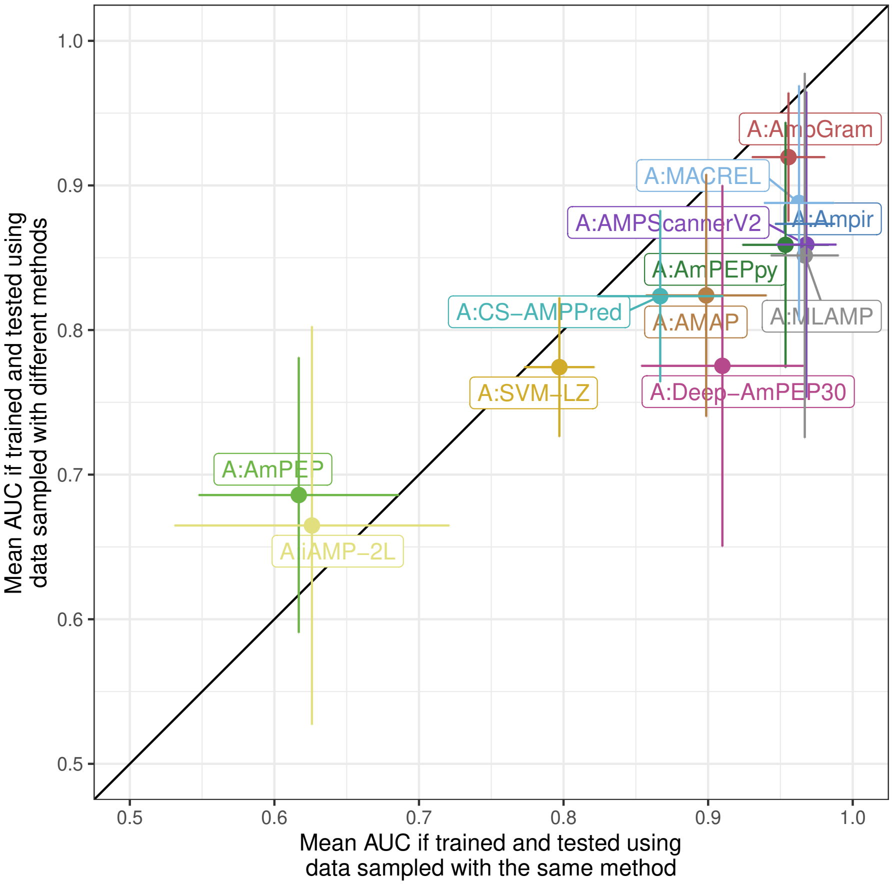

Figure S70: Model architecture performance with standard deviations. The x-axis represents mean AUC for architectures trained and tested on sets generated by the same negative data sampling method. The y-axis represents mean AUC for architectures trained and tested on sets generated by different negative data sampling methods. The architectures on the right of the diagonal perform better when the training and benchmark sample are produced by the same method while the architectures on the left when the methods are different. Horizontal and vertical lines indicate standard deviations.

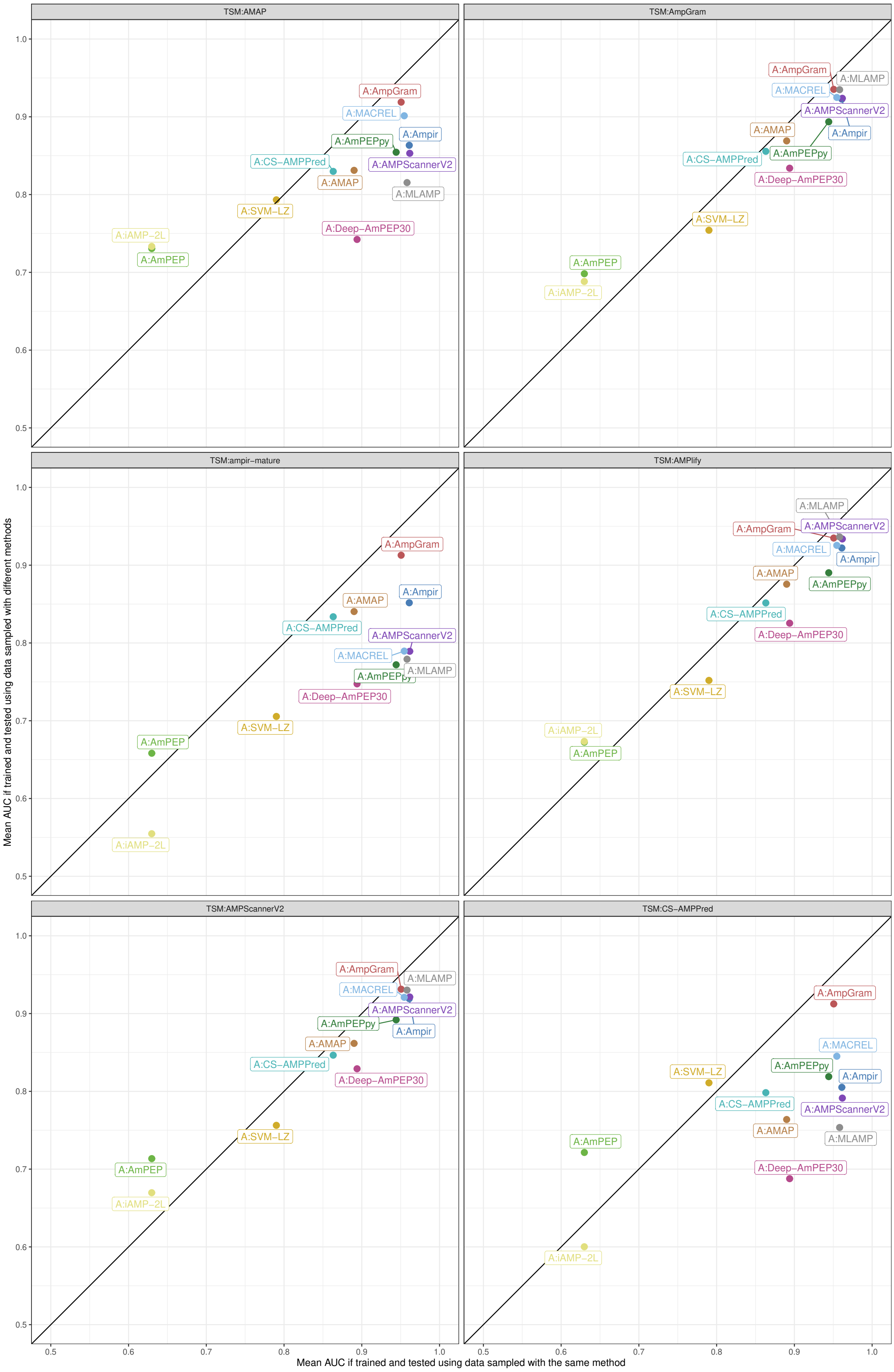

Figure S71: Model architecture performance depending on the negative data sampling method used for training. The x-axis represents mean AUC for architectures trained and tested on sets generated by the same negative data sampling method. The y-axis represents mean AUC for architectures trained and tested on sets generated by different negative data sampling methods. The architectures on the right of the diagonal perform better when the training and benchmark sample are produced by the same method while the architectures on the left when the methods are different.

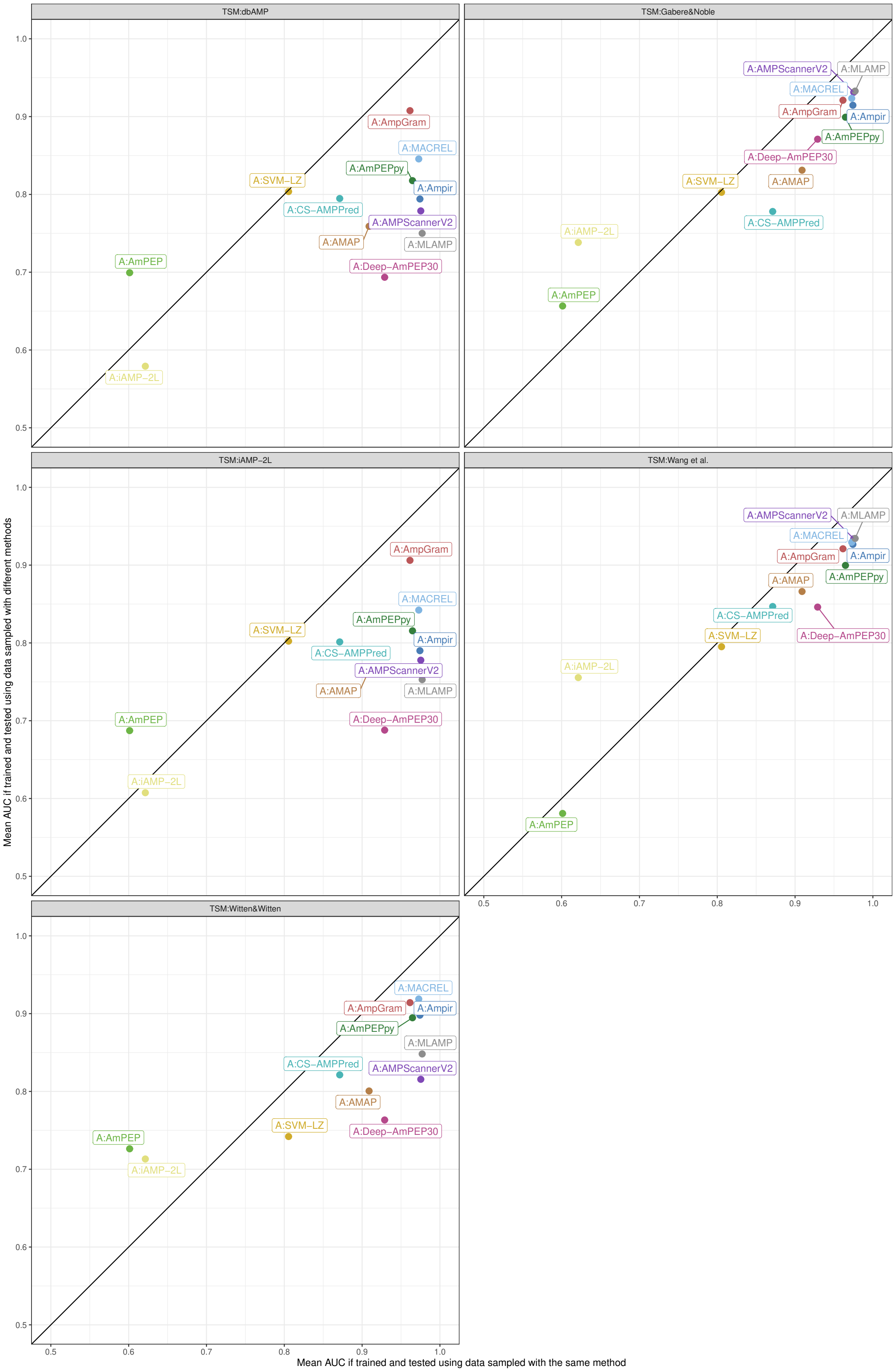

Figure S71 (continued): See description on the previous page.

[15] Sneh Lata, Nitish K Mishra, and Gajendra PS Raghava. AntiBP2: improved version of antibacterial peptide prediction. *BMC Bioinform.*, 11(S1), 2010.

[16] Faiza Hanif Waghu, Ram Shankar Barai, Pratima Gurung, and Susan Idicula-Thomas. CAMPR3: a database on sequences, structures and signatures of antimicrobial peptides. *Nucleic Acids Res.*, 44(D1):D1094–D1097, 2016.

[17] Shaini Joseph, Shreyas Karnik, Pravin Nilawe, V. K. Jayaraman, and Susan Idicula-Thomas. ClassAMP: A prediction tool for classification of antimicrobial peptides. *IEEE/ACM Trans Comput Biol Bioinform*, 9(5):1535–1538, 2012.

[18] William F Porto, Állan S Pires, and Octavio L Franco. CS-AMPPred: an updated SVM model for antimicrobial activity prediction in cysteine-stabilized peptides. *PLoS One*, 7(12):e51444, 2012.

[19] Jhih-Hua Jhong, Yu-Hsiang Chi, Wen-Chi Li, Tsai-Hsuan Lin, Kai-Yao Huang, and Tzong-Yi Lee. dbAMP: an integrated resource for exploring antimicrobial peptides with functional activities and physicochemical properties on transcriptome and proteome data. *Nucleic Acids Res.*, 47(D1):D285–D297, 2019.

[31] Jacob Witten and Zack Witten. Deep learning regression model for antimicrobial peptide design. *BioRxiv*, page 692681, 2019.

[32] Wararat Chiangjong, Somchai Chutipongtanate, and Suradej Hongeng. Anticancer peptide: Physicochemical property, functional aspect and trend in clinical application. *International Journal of Oncology*, 57(3):678–696, 2020.

[33] Jasmeet S Khara, Ying Wang, Xi-Yu Ke, Shaoqiong Liu, Sandra M Newton, Paul R Langford, Yi Yan Yang, and Pui Lai Rachel Ee. Anti-mycobacterial activities of synthetic cationic $\alpha$-helical peptides and their synergism with rifampicin. *Biomaterials*, 35(6):2032–2038, 2014.
